# Genetic influence on resting state networks in young male and female adults

**DOI:** 10.1101/2021.02.13.430156

**Authors:** Gyujoon Hwang, Arman P. Kulkarni, Rosaleena Mohanty, Cole J. Cook, Veena A. Nair, Barbara B. Bendlin, Elizabeth Meyerand, Vivek Prabhakaran

**Affiliations:** Department of Medical Physics, University of Wisconsin-Madison, 1111 Highland Ave, Rm 1005, Madison, WI 53705, USA; Department of Biomedical Engineering, University of Wisconsin-Madison, 1415 Engineering Drive, Madison, WI 53706, USA; Division of Clinical Geriatrics, Department of Neurobiology, Care Sciences and Society, Karolinska Institutet, Stockholm, Sweden; Department of Radiology, University of Wisconsin-Madison, E3/366 Clinical Science Center, 600 Highland Avenue, Madison, WI 53792, USA; Department of Medicine, University of Wisconsin-Madison, 1685 Highland Ave, 5158 Medical Foundation Centennial Building, Madison, WI 53705, USA; Department of Neurology, University of Wisconsin-Madison, Medical Foundation Centennial Building, 1685 Highland Avenue, Madison, WI 53705, USA

**Keywords:** Heritability, Twin Study, Connectome, Resting-state fMRI

## Abstract

Determining genetic versus environmental influences on the human brain is of crucial importance to understand the healthy brain as well as in a variety of disease and disorder states. Here we propose a unique, minimal assumption, approach to investigate genetic influence on the functional connectivity of the brain using 260 subjects” (65 monozygotic (MZ) and 65 dizygotic (DZ) healthy young adult twin pairs) resting state fMRI (rsfMRI) data from the Human Connectome Project (HCP). For any given resting state connection between twin pairs, the connection strengths across pairs were subtracted from each other in both directions. By applying the F-Test for equality of variances per connection, we found that there were a number of significant connections that demonstrated greater variance among dizygotic pairs in comparison to monozygotic pairs, implying these connections were under significant genetic influence. These population (DZ-MZ) results remained true irrespective of gender, with the caveat that certain connections were significant on a gender-specific basis. This is the first study to our knowledge to assess the heritability across young healthy adults both in general and specific to gender.

**Population Results & Discussion:** At the population level, there appears to be a posterior to anterior gradient of more to less genetic influence on brain connections and networks with visual > temporal, parietal > frontal. There was a high density of genetically-influenced functional connections predominantly involving posterior regions or networks of the brain: Visual Networks (VNs - primary visual, early visual, dorsal stream and ventral stream visual cortices, MT+ complex). These posterior regions of the brain with greater genetic influence are implicated for example in visual, perceptual, dorsal (“where”) and ventral (“what”) visuospatial processing streams (VNs).

There was a low-density or paucity of genetically-influenced functional connections predominantly involving anterior regions or networks of the brain comprising Task Positive Networks (TPNs): FrontoParietal Networks (FPNs - dorsolateral prefrontal, orbital and polar frontal, midcingulate, insular and frontal opercular, superior and inferior parietal cortices); FrontoTemporal Networks (FTNs - inferior frontal, posterior opercular, early auditory, auditory association cortices); Sensorimotor Networks (SMNs - premotor, somatosensory, paralobular, and motor cortices); These anterior regions of the brain with lesser genetic influence are implicated in various TPN processes; for example in high-level cognitive and affective processes such as working memory, executive function, reasoning, attentional and impulse control, emotional judgement and decision making (FPNs); language and auditory processes (FTNs); action-planning and movement processes (SMN).

There was a mix of high (posterior) and low (anterior) density of genetically influenced functional connections involving the extended Default Mode Network (eDMN). Specifically, there was a high density of genetically-influenced functional connections involving predominantly posterior-medial regions of eDMN - hippocampus and precuneus/posterior cingulate cortices; There was a low density of genetically influenced connections involving anterior regions (anterior cingulate and medial prefrontal) and lateral (inferior parietal, temporoparietooccipital) regions of the eDMN. The eDMN is involved in low-level cognitive and affective processes such as those involved in episodic memory retrieval, mental imagery, introspection, rumination, evaluation of self and others.

These differences in genetic influence on posterior (more) vs. anterior (less) brain regions may have implications in terms of the environmental influence (e.g., education, school and work environment, family and home environment, social interaction with friends and peers, medications, nutrition, sports and physical exercise) on posterior (less) vs. anterior (more) portions of the brain during development and later in life.

**Gender-Specific Results & Discussion:** As noted at the population level, both males and females were under extensive genetic influence in terms of network interactions involving visual cortices. In addition, males were more genetically influenced in terms of network interactions involving auditory-language related cortices compared to females. This finding suggests that males may be more functionally “hard-wired” and females may be more environmentally influenced and shaped in terms of auditory-language systems than males.

As noted at the population level, both males and females were under extensive genetic influence in terms of interactions involving the eDMN which is considered a central hub of the brain for various processes such as internal monitoring, rumination and evaluation of self and others, as noted previously. In addition, males also were more genetically influenced compared to females in terms of intranetwork and internetwork interactions of eDMN and other brain regions (occipital, temporal, parietal, and frontal regions) involved in various task-oriented processes and attending to and interacting with the environment which comprise part of the Task Positive Networks (TPNs). There were also nearly five times more genetically influenced functional connections in males (310) than females (64) suggesting that male brains are more genetically influenced, i.e. functionally “hard-wired”, than females. This result suggests differences in genetic predisposition in males (more) vs. females (less) in terms of interplay of attending to task-oriented interactions with the environment (TPNs) vs. internal and external interactions with self and others (eDMN). This finding may also have implications in terms of brain plasticity differences in males (less) versus females (more) in terms of ability to react or adapt/maladapt to environmental influences (e.g. task completion demands, psychosocial stressors, positive and negative feedback, meditation, cognitive behavioral therapy, pharmacotherapy) and their overall malleability.

These results reveal the similarities and differences of genetics and environmental influences on different connections, areas, and networks of the resting state functional brain in young healthy males and females with implications in development and later in life. This unique method can be applied in healthy as well as in patient populations to reveal the genetic and environmental influences on the brain.

**Significance:** There were high vs. low genetic influences on posterior vs. anterior brain regions involved in low-level visuospatial processes vs. high-level cognitive processes such as reasoning and language respectively. This finding may have implications in terms of the brain to be environmentally influenced (e.g., school, work and home environment) during development and later in life.

There were nearly five times more genetically influenced functional connections in males than females in brain regions involved in task-oriented interactions with environment vs. interactions with self and others. This finding may have implications in terms of brain plasticity differences in males (less) versus females (more) in terms of ability to adapt/maladapt to environmental influences (e.g. task completion demands, psychosocial stressors, various therapies) and their overall malleability. This is the first study to our knowledge to assess the heritability across young healthy adults both in general and specific to gender.

## 1. Introduction

Genetics and environmental factors are two key components in characterizing every individual. Quantifying these two components meaningfully is of great importance in understanding their influence on healthy individuals as well as in a variety of diseases and disorders. Studies have shown that normal and aberrant brain functional connectivity has been found to have strong genetic dependence and linkage in normal development and aging (Gao et al., 2017; Hoff et al., 2013; Yang et al., 2016) as well as with a variety of disease states and disorders (A. et al., 2016; Glahn et al., 2010). Resting state networks (RSNs) may be endophenotypes (Glahn et al., 2010), and more recently, the advent of MR fingerprinting suggests that patterns of resting state functional connectivity (rsFC) may be used to identify specific individuals (Finn et al., 2015). In order to be classified as an endophenotype, a phenotype must satisfy a set of criteria, one of which necessitates heritability (Elliott et al., 2019; Glahn et al., 2010).

Currently, modern analyses investigating heritability utilize either 1) the ACE (A = Additive Genetics, C = Shared Environment, E = Unshared Environment) Model (Zyphur et al., 2013), or Falconer”s Formula (Falconer and Mackay, 1996). The ACE model primarily evaluates narrow-sense heritability due to additive genetics (*h*^*2*^) based on monozygotic (MZ) or dizygotic (DZ) twin differences, in which this heritability is defined as the total variation attributed to genes. Falconer”s Formula instead evaluates broad sense heritability, which includes all genetic influences on phenotypic variation (Falconer and Mackay, 1996). In general, MZ twins are known to uniquely have identical genomes as opposed to DZ twins, implying phenotypic variability between these twin groups most likely is attributable to genetic effects and not environmental effects (Joseph, 2013). While models such as ACE and Falconer”s method (Falconer and Mackay, 1996; Zyphur et al., 2013) currently exist, there remains room for a more parsimonious method in the interest of characterizing individuals based on rsFC while at the same time evaluating genetic influence of the functional brain.

Therefore, the purpose of this study was two-fold: 1) to develop a novel approach in order to evaluate rsFC phenotypic variance differences between MZ and DZ twin pairs to elucidate genetic influence on the functional brain with minimal assumptions and 2) to determine the influence of gender differences on the functional brain utilizing the aforementioned twin pairs. This is the first study to our knowledge to assess the heritability across young healthy adults both in general and specific to gender. The Human Connectome Project (HCP) (Smith et al., 2013; Van Essen et al., 2013) is known for its high temporal and spatial resolution fMRI data, large sample size and a well-characterized high-quality imaging and behavioral dataset. From this dataset, we use 65 MZ and 65 DZ young healthy adult twin pairs as subjects, each scanned during four sessions over two days. We then demonstrate that all resting state connectivity phenotypic variances of those significantly different are smaller for MZ twins as opposed to DZ twins at the population level, and that gender-wise differences in significant connections exist at the gender-specific level. These results demonstrate crucial findings that verify the overall genetic contributions that would be expected for monozygotic twins and for dizygotic twins. These findings further elucidate which brain regions are most influenced by genetics as opposed to those which may be unhindered and malleable to environmental influences. This outcome encourages further implementation of this novel methodology in more aspects of functional connectivity, genetics, and characterization of individuals, or as a precursor to more complicated analyses such as those aforementioned.

## 2. Materials and Methods

### 2.1. Demographics

65 MZ (age=29.1±3.6 years, 35 female twin pairs) and 65 DZ (age=29.1±3.6 years, all same-sex twins, 37 female twin pairs) healthy twin pairs from the HCP were analyzed. Table 1 summarizes the demographics. There were no significant differences in age, gender ratio, education, DVARS as well as in education score differences (Wilcoxon Rank Sum Test) between twin pairs across any group (p>0.15, MZ versus DZ, Male MZ (MMZ) versus Male DZ (MDZ), Female MZ (FMZ) versus Female DZ (FDZ)).

**Table 1.**
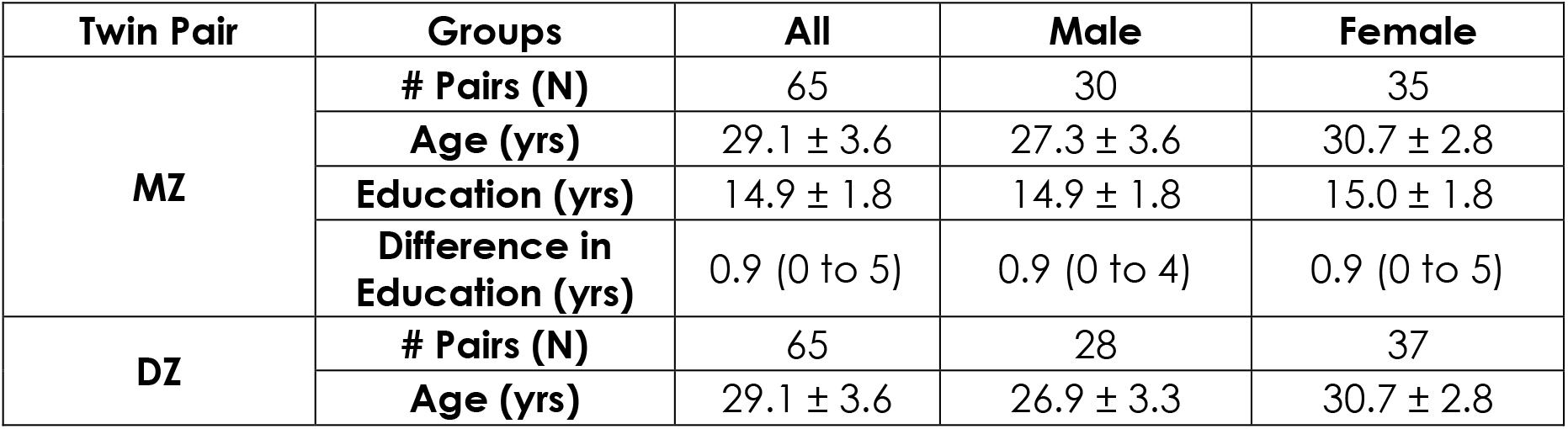
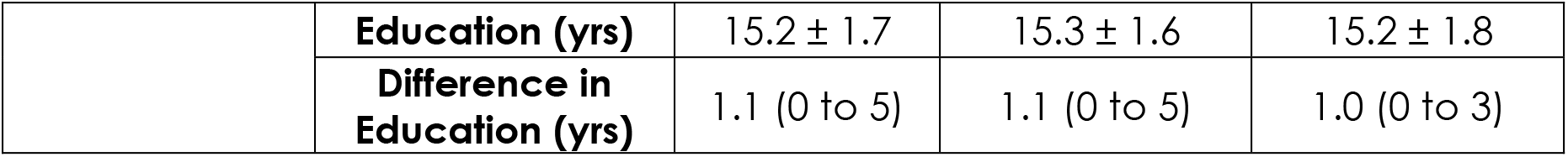
Demographics summary. Age, gender, education, DVARS and the differences in education were all matched statistically (p>0.15) between groups in comparison.

### 2.2. Data Collection

60 minutes of eyes fixation rsfMRI data which had been acquired over four scanning sessions across two days as part of the HCP and had been minimally processed were used (Glasser et al., 2013). The processed images are publicly available on the ConnectomeDB web database (Hodge et al., 2016). Permission was obtained from the HCP Connectome Coordination Facility to access restricted demographic and behavioral data. MR images were acquired using a 3T Connectome scanner adapted from Siemens Skyra at Washington University, in St. Louis using a 32-channel head coil with simultaneous multi-slice imaging (SMS, 8 bands, 72 slices, TR=720ms, TE=33.1ms, 2.0 mm isotropic voxels). The pre-processing included the HCP minimal preprocessing pipelines (Glasser et al., 2013) and FMRIB”s Independent-component-analysis-based X-noisifier (FIX) correction method (Salimi-Khorshidi et al., 2014). There were no significant differences in the means of absolute and relative root-mean-squared motion, as well as DVARS, between MZ and DZ groups.

The only additional preprocessing steps performed were concatenation of the four 15-minute scans (2 pairs with opposite phase encoding gradients) and band-pass filtering (0.01-0.1Hz). Time series from 360 cortical brain regions defined by HCP”s Glasser parcellation (Glasser et al., 2016) and 19 subcortical regions from FreeSurfer”s subcortical segmentation (Fischl et al., 2002) were extracted per subject. Raw pairwise Pearson”s correlations were computed and then transformed to Fisher Z scores to generate the connectivity matrices.

### 2.3. Twin Analysis

For a given twin pair, the differences in the connectivity matrices were calculated. To account for the arbitrary nature in the order of subtraction, every difference was duplicated with its opposite sign value. Essentially, then, two difference matrices were generated per twin pair: twin A – twin B and twin B – twin A. The resulting difference matrices for the monozygotic and dizygotic groups were compared using the F-Test of equality of two variances. Multiple comparison corrections were performed using Bonferroni correction (Abdi, 2007). Normality for each significant connection was tested via the subtracted connection distributions using the Kolmogorov-Smirnov (KS) Test (Massey, 1951). The twin analysis and subsequent analyses were repeated to compare between male MZ and DZ pairs, as well as female MZ and DZ pairs.

### 2.4. Network Categorization

Network categorization was based on Glasser defined networks (Glasser et al., 2016). The subcortex was further expanded based on the FreeSurfer based parcellation (Desikan et al., 2006) of anatomical regions (i.e., cerebellum, thalamus, caudate, putamen, etc). The resulting significant network-network connections were binned based on each respective network into a newly defined network connectivity matrix to demonstrate the network effect of the results. Connectogram visualizations of these results were generated using Circos (Krzywinski et al., 2009).

### 2.5. Region Validation Analysis

Glasser et al. (Glasser et al., 2016) defines 22 cortical regions (or networks) that are constituted by 360 parcels (or areas). FreeSurfer subcortical parcellation has 10 regions. A permutation test (Nichols and Holmes, 2002) was performed to reveal which of these 32 regions contain a significantly large or conversely small number of the significant connections from the F-Test analysis. For example, if the number of total whole-brain significant connections was N, then N connections were randomly selected from 71,631 possible connections, and their starting and ending regions were analyzed. If a connection started and ended in the same region, the region was counted twice. This procedure was repeated 10,000 times and the distributions of the occurrences of parcel counts were generated per region. Then, the *p*-values for the actual results were determined based on the significant region count results and their comparison against these null distributions.

### 2.6. Intra-class Correlation Validation Analysis

To validate our findings, we computed from the rsFC measures—which were shown to be significant at the whole brain level from the F-Test analyses—the average rsFC value on a subject by subject basis. The intra-class correlation (ICC) of this average measure was calculated across MZ twin pairs and DZ twin pairs separately, for a single ICC measure per grouping.

## 3. Results

### 3.1. Population Differences

Between MZ and DZ twin groups, 542 out of 71,631 connections showed significant difference in variance (corrected *p*<0.05), and all connections had greater variances in the DZ group. Out of 542, 300 were intrahemispheric (131 connections were left-hemispheric, 169 right-hemispheric) and 242 interhemispheric (Figure 1).

**Figure 1.**
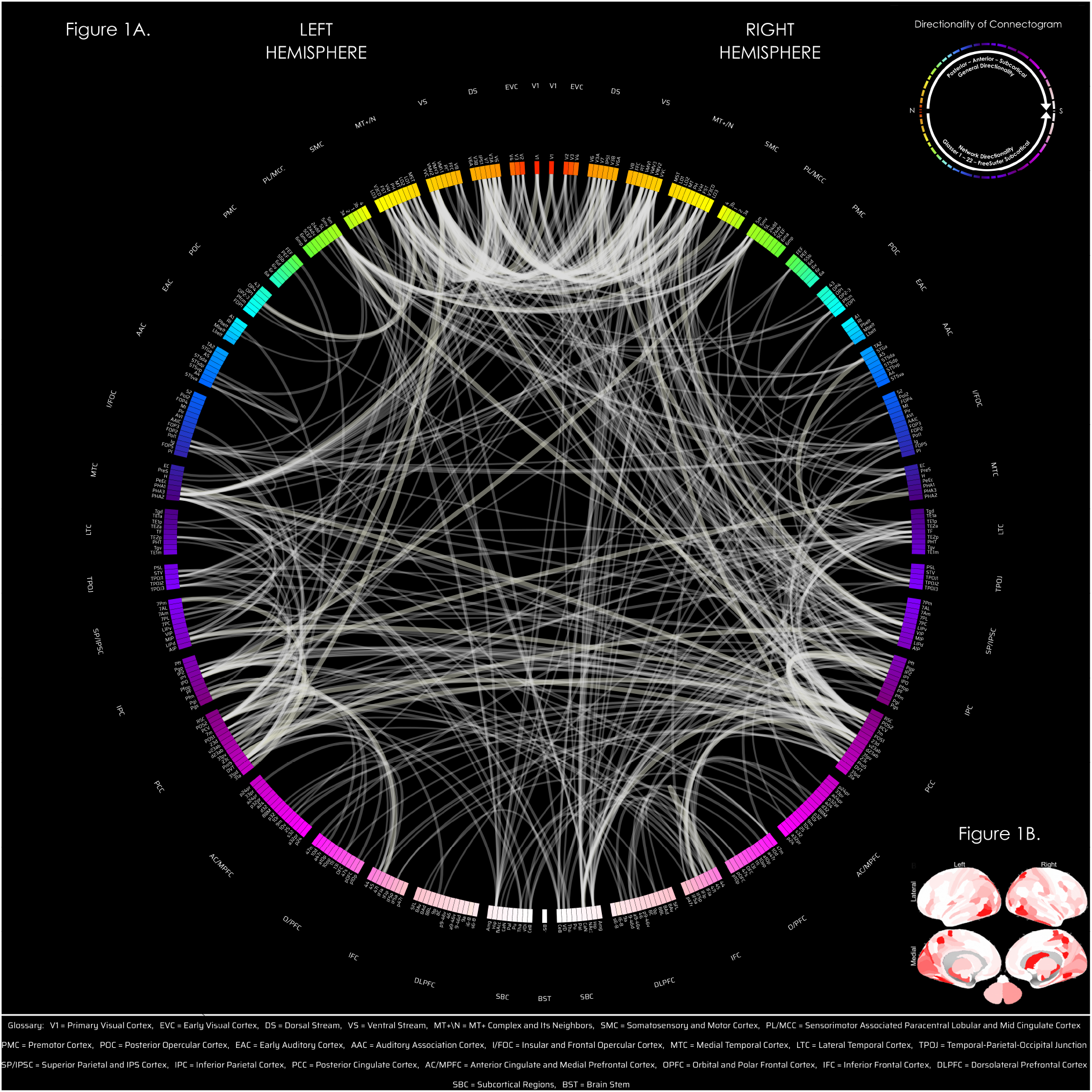
Connectogram of resting state functional connections that demonstrated significantly larger variances (corrected, thin white 0.01≤p<0.05; medium thick white 0.0001≤p<0.01; thick white p<0.0001) in the DZ twin group compared to the MZ twin group (Figure 1A). These brain connections were found to be influenced by genetics. Regions with a higher number of connections involving that region (that were more frequently influenced by genetics) are colored increasingly red (Figure 1B). At the population level, there was a high density of genetically-influenced functional connections involving posterior regions of the brain and a low density or paucity of genetically-influenced functional connections involving anterior regions of the brain. Directionality of the connectogram (top right) is shown to emphasize the exact direction in terms of networks and general direction of anatomy from north (N) to south (S). For the full list of Glasser parcels see Glasser et al., 2016. The subcortical FreeSurfer region labels are: BSt = Brain Stem, Amg = Amygdala, Hip = Hippocampus, NAcc = Accumbens, CaN = Caudate, Pal = Globus Pallidus, Pu = Putamen, Tha = Thalamus, VDi = Ventral Diencephalon, and CeB = Cerebellum.

At the population level, there appeared to be a posterior to anterior gradient of more to less genetic influence, both at the parcel- and at the network-level, with visual > temporal, parietal > frontal (Figure 1, Figure 3 left panel).

There was a high density of genetically-influenced functional connections involving posterior regions of the brain: Visual Networks (VNs) - primary visual cortex, early visual cortex, dorsal stream and ventral stream visual cortices, MT+ complex.

There was a low density or paucity of genetically-influenced functional connections involving anterior regions of the brain: FrontoParietal Networks (FPNs) - dorsolateral prefrontal, orbital and polar frontal, midcingulate, insular and frontal opercular, Superior and Inferior Parietal cortices; Frontotemporal Networks (FTNs) - inferior frontal, posterior opercular, early auditory, and auditory association cortices; and Sensorimotor Networks (SMNs) - premotor, somatosensory, sensorimotor paralobular, and motor cortices.

There was a mix of high (posterior) and low (anterior) density of genetically influenced functional connections involving the extended Default Mode Network (eDMN). Specifically, there was a high density of genetically-influenced functional connections involving predominantly posterior-medial regions of eDMN - hippocampus and precuneus/posterior cingulate cortices; There was a low density of genetically influenced connections involving anterior regions (anterior cingulate and medial prefrontal) and lateral (inferior parietal, temporoparietooccipital) regions of the eDMN.

These findings were seen both at the parcel level (Figure 1) as well as at the overall network level (Figure 3 left panel and Table 2).

**Table 2.**
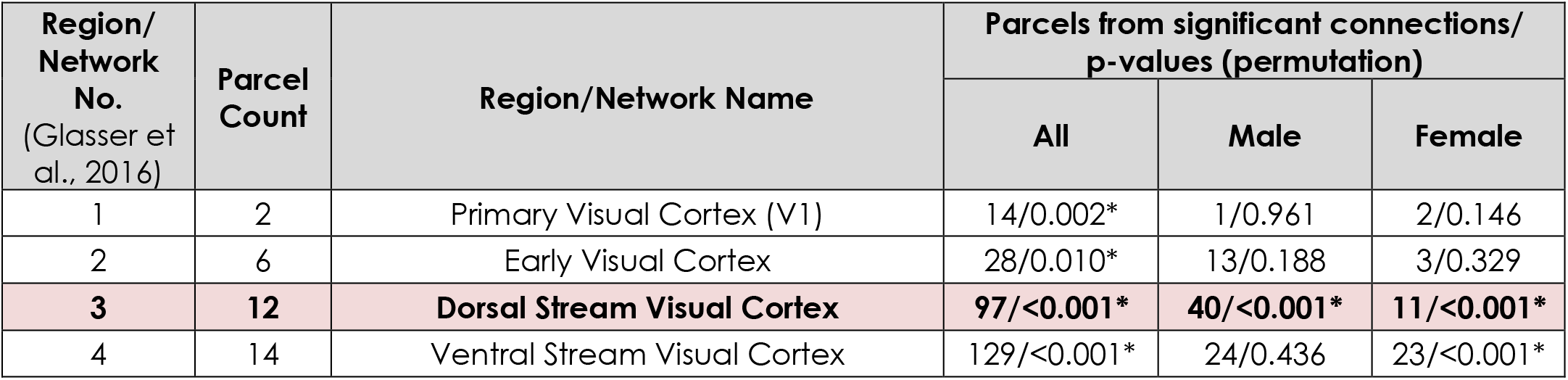
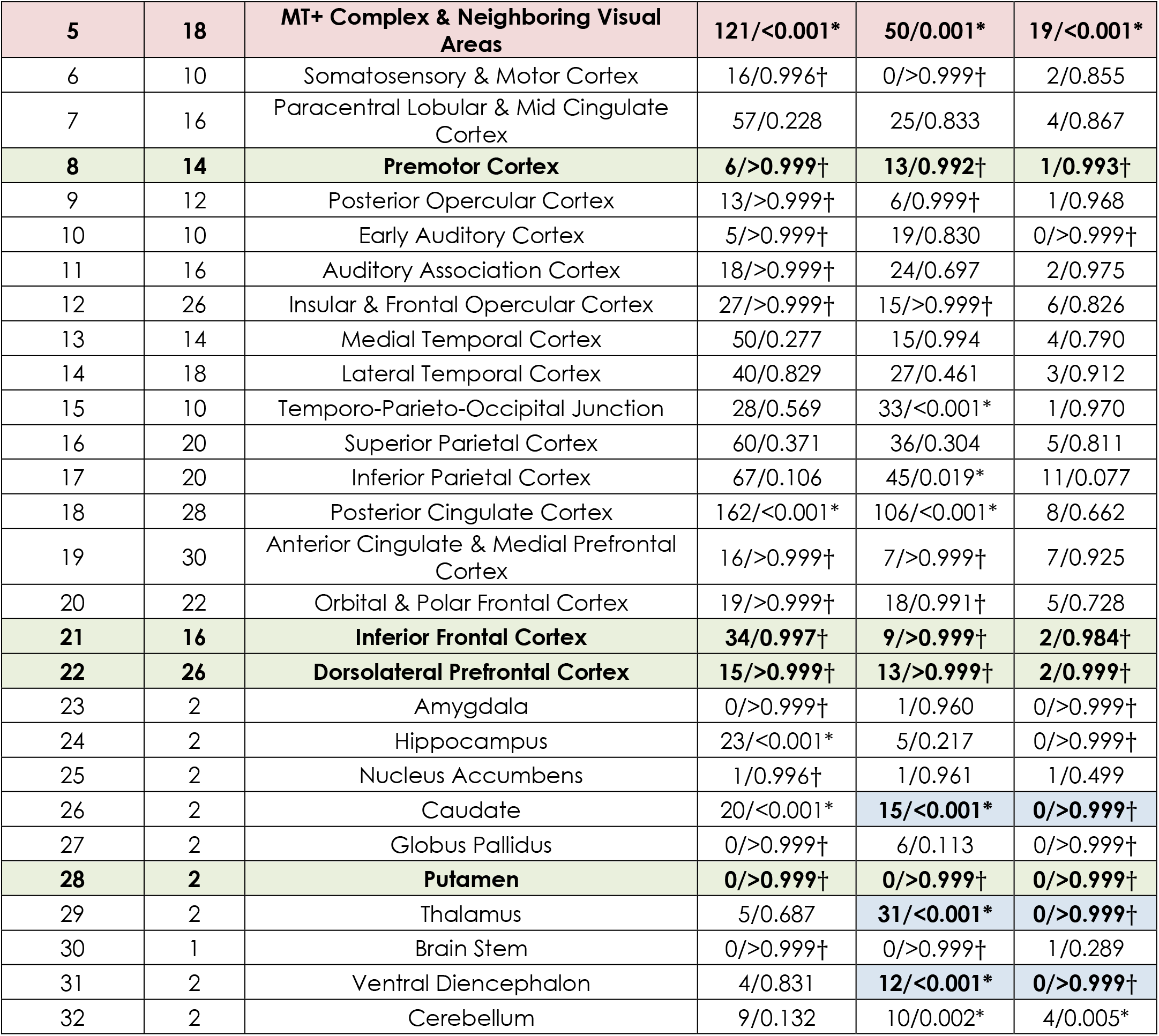
Results from the region analysis. The first 22 regions are defined by Glasser parcellation (Glasser et al., 2016) and the subcortex (regions 23-32) is from the FreeSurfer segmentation (Fischl et al., 2002). P-values were determined from a right-tailed permutation test with 10,000 iterations. *p<0.025, which indicates that there were significantly larger number of significant connections found in the region than random. † p>0.975, which indicates that there were significantly smaller number of significant connections found in the region than random.

Other regions that were significantly genetically influenced at the network level for the population included caudate regions, while those that were not significantly influenced were putamen, globus pallidus, nucleus accumbens, amygdala, and brainstem (Table 2).

### 3.2 Gender Differences

#### Males

Comparing only male twins (F-Test for MMZ versus MDZ), 310 connections (168 intrahemispheric [75 left, 93 right], 142 interhemispheric) showed significant differences in variance (corrected *p*<0.05) and all connections had greater variances in the MDZ group (Figure 3 middle panel, Table 2). 65 of these 310 connections were exclusive to the male group (these connections showed the opposite effect in females, i.e., larger variances in the FMZ twin group compared to FDZ). Figure 2 left demonstrates these 310 connections that were significant in males. The most significant connections in males involved VNs - Dorsal Stream, MT+ complex, posterior and lateral portions of eDMN - posterior cingulate, parietal (inferior parietal), and temporal cortical regions (temporoparietaloccipital). Anterior and medial portions of eDMN and other TPNs (occipital, parietal, temporal, auditory cortices, frontal cortices) were seen to be significantly involved in the network-network and parcel-parcel interactions (Figure 2 left, Figure 3, middle)

**Figure 2.**
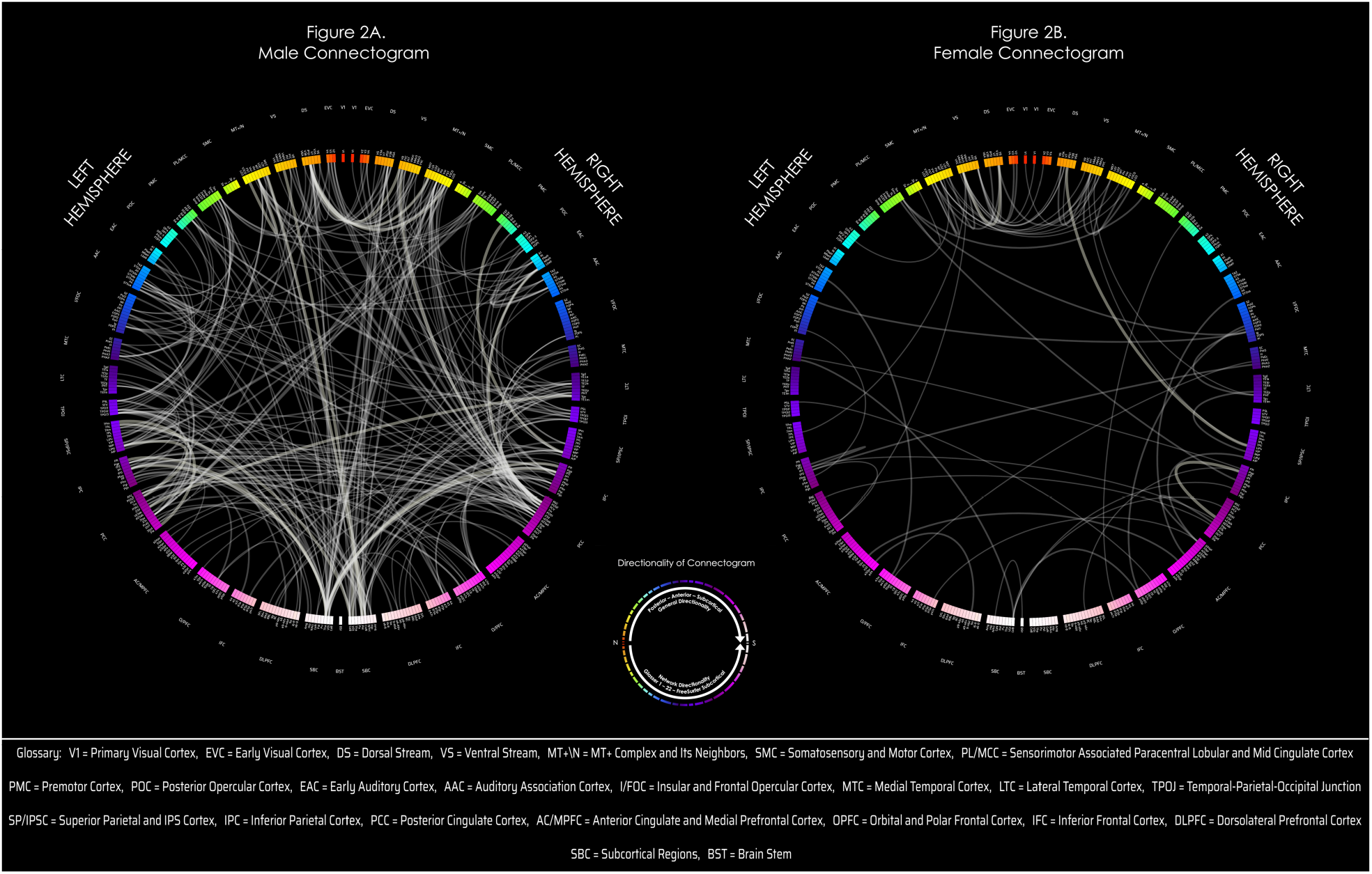
Figure 2A: male connectogram of resting state connections that demonstrated significant genetic influence (corrected p<0.05), (DZ group showing larger variance compared to MZ). Figure 2B: female connectogram of resting state connections that demonstrated significant genetic influence (corrected p<0.05) (demonstrates DZ group showing larger variance compared to MZ; one connection where DZ group shows lesser variance compared to MZ not shown). There were nearly five times as many significant connections in males (310) compared to females (64). There was also a greater interplay of eDMN with various TPNs in males, than in females. Directionality of the connectogram (middle) is shown to emphasize the exact direction in terms of networks and general direction of anatomy from north (N) to south (S). For the full list of Glasser parcels see Glasser et al., 2016. The subcortical FreeSurfer region labels are: BSt = Brain Stem, Amg = Amygdala, Hip = Hippocampus, NAcc = Accumbens, CaN = Caudate, Pal = Globus Pallidus, Pu = Putamen, Tha = Thalamus, VDi = Ventral Diencephalon, and CeB = Cerebellum.

#### Females

Comparing only female twins, 64 connections (35 intrahemispheric [19 left,16 right], 28 interhemispheric, one from left hemisphere to brainstem) showed significantly greater variance (corrected *p*<0.05) in the DZ group, and one interhemispheric connection showed the opposite effect (Figure 2 & 3 right panel, Table 2). 9 of these 64 connections were exclusive to the female group (these connections showed the opposite effect in males, i.e. larger variances in MMZ compared to MDZ). Figure 2 right demonstrates these 64 connections that were significant in females. The most significant connections and network interactions in females involved the Visual Networks – Dorsal and Ventral Stream, MT+ Complex (Figure 2 & 3 right panel, Table 2).

**Figure 3.**
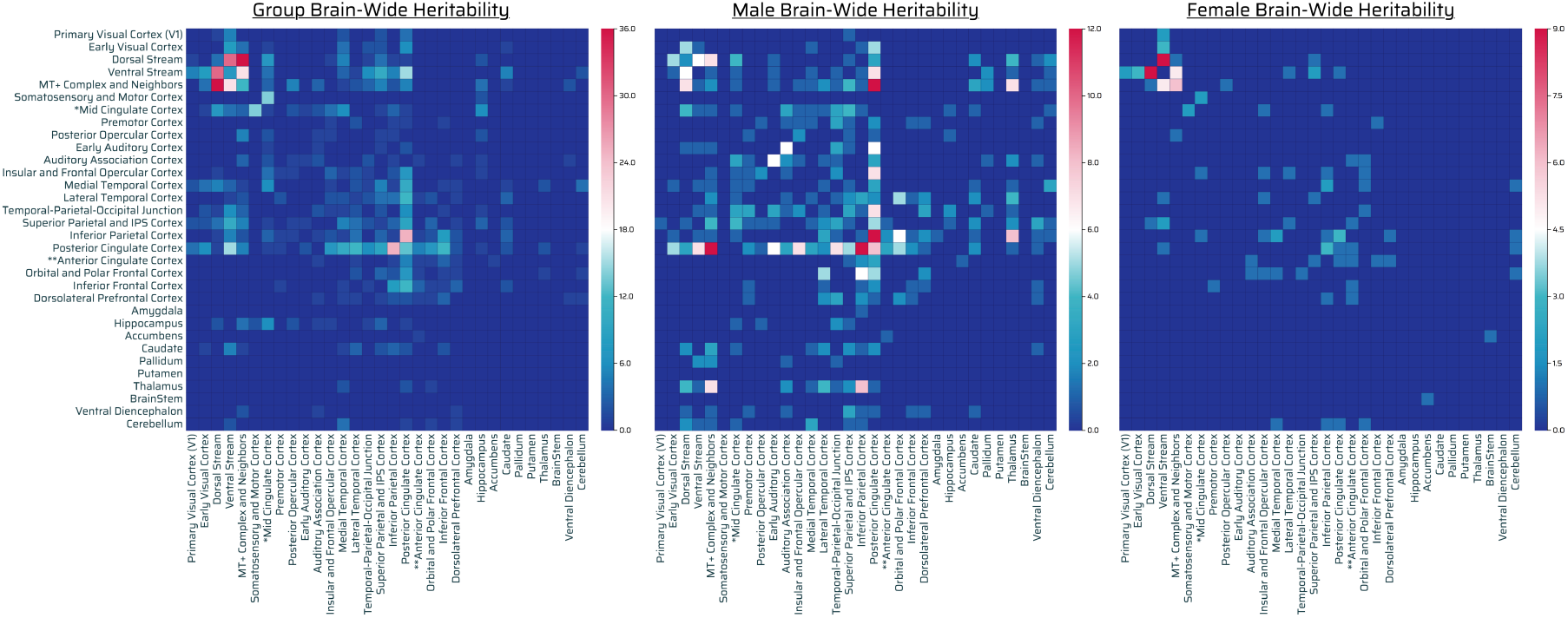
Network tabulation matrices using the Glasser defined network parcels for significantly genetically-influenced connections across all (male and female) twins (left), only male twins (middle), and only female twins (right). Overall there appears to be a gradient of more genetic influence of network-network interactions with visual > temporal, parietal > frontal (Figure 3, left). Males demonstrated a large number of genetically-influenced network-network interactions involving visual networks, TPNs, and eDMN and overall a paucity of network-network involvement of frontal regions (Figure 3, middle). Females demonstrated mainly genetically-influenced network-network interactions involving visual networks, with a small number of interactions involving TPNs and eDMN and overall a paucity of network-network involvement of frontal regions (Figure 3, right). Males overall displayed much greater network-network interactions than females. *Mid Cingulate Cortex = Sensorimotor Associated Paracentral Lobular and Mid Cingulate Cortex. **Anterior Cingulate Cortex = Anterior Cingulate and Medial Prefrontal Cortex.

#### Other Cortical Regions

Although at the network level (Table 2), dorsolateral prefrontal, inferior frontal, anterior cingulate and medial prefrontal area were not significant for males or females, specific parcels of these network were significantly involved in males and females (Figure 2). Posterior cingulate region, precuneus, hippocampus involvement was significant in males at both the network and parcel level (Table 2 and Figure 2), while it was significant in females only at the parcel level (Figure 2).

Therefore, the DMN and TPNs were significantly genetically influenced much more in males than females. See appendix for the full list of connections that were significant at the parcel level.

#### Subcortical

Genetics appeared to influence a high density of connections involving cortico-basal-ganglia-thalamic-cortical regions predominantly in males. Ventral diencephalon involvement was seen predominantly in males. Cerebellar region involvement and cortico-cerebellar connections were seen in both males and females.

#### Overall patterns

There were nearly five times as many significant connections in males (310) compared to females (64) (Figure 2 and Figure 3 middle and right). There was also a greater genetic influence in terms of intra and internetwork interactions of DMN and TPNs in males than in females (figure 2, figure 3 middle, right). In terms of sensory systems, there was genetic influence of involvement of Visual Networks in both females and males; there was greater genetic influence of involvement of Auditory Networks in males than females (figure 3 middle, right).

### 3.2. Network-Network Heritability Summary

The breakdown of network-network interactions and the extent of genetically-influenced connections per network-network interaction can be seen in Figure 3. Given the arbitrary nature of normalization methods, the raw data tabulation was picked to illustrate the interaction. Overall there appears to be a posterior to anterior gradient of more to less genetic influence of network-network interaction with visual > temporal, parietal > frontal (Figure 3, left).

Males showed a large number of genetically-influenced network-network interactions involving visual, parietal, and temporal regions with a paucity of network-network involvement of frontal regions (Figure 3, middle). Females demonstrated genetically-influenced network-network interactions involving visual regions (figure 3, right). Males overall show much greater genetically-influenced network-network interactions than females. There was genetic influence of extended Default mode network involvement in the overall population as well as in male and females (Table 2, Figure 3). In addition, males show more genetically-influenced network interactions of eDMN with TPNs (occipital, parietal, temporal and auditory, frontal cortices) than females (Figure 3 middle, right). These results were overall convergent with what was noted at the parcel level.

### 3.3. Validation with ICC

The correlations for the average constructed metric for all MZ pairs, regardless of gender, were shown to be higher than that for all DZ pairs (ICC_MZ_ ≤ 0.87, ICC_DZ_≤ 0.21), lending validation to our approach (Table 3). Albeit, it should be noted that the MDZ ICC value is negative, implying that the true ICC is low (Bentley et al., 2017; Taylor, 2009).

**Table 3.**
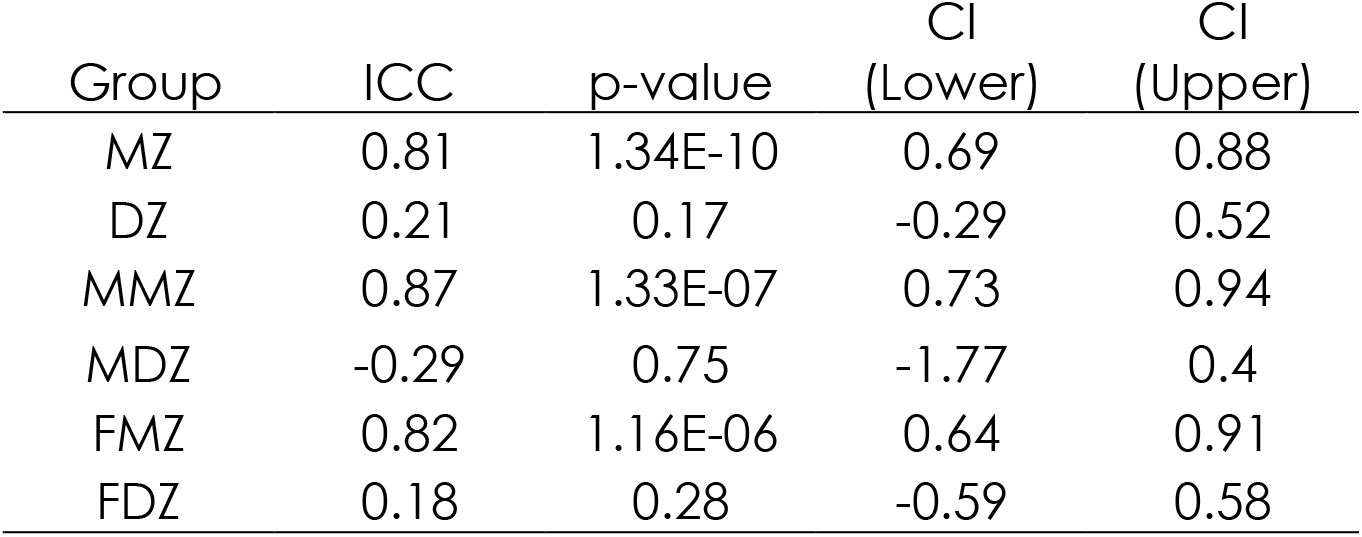
Intra-class correlations (ICCs) of average of significant rsFC measures per analysis. MZ ICCs are seen to be greater than that of DZ ICCs.

## 4. Discussion

### 4.1. Novelty of Method

The proposed method models the phenotypic variance in pair-wise connectivity similarity of MZ and DZ twins separately, and then compares these two variations with one another. Phenotypic variance is known to be composed of V_P_ =V_G_ + V_E_, (Falconer and Mackay, 1996) where V_G_ is the impact on the variance due to genetic sources and V_E_ is the impact on the variance due to environmental sources. Given that MZ twins have identical genomes, and DZ twins have half identical genomes, we presuppose that the influence of environmental effects in this analysis is moderate to negligible if the phenotypic variances in MZ twins is not equal to, or smaller than, that of DZ twins. This approach provides insight into the likelihood of genetic influence on the phenotypic variance differences. Overall, although it does not resolve these confounds as does Falconer”s Method to a limited degree, or the ACE model, it avoids the assumptions and issues that may accompany these approaches. For example, Falconer”s Method allows for heritability estimates greater than 1 or less than 0, causing interpretability issues (Corruccini et al., 1990).

### 4.2. Overall Findings

Our study aimed to investigate the genetic and gender influence of individual rsFC connections based on the Glasser and FreeSurfer subcortical parcellation mapping. This is the first study to our knowledge that specifically investigates rsFC heritability in healthy young adults with regard to both males and females. We discuss these main findings at the population level and gender-specific level.

#### 4.2.1. Population Level

At the population level, there was a high density of genetically-influenced functional connections involving posterior regions of the brain: VNs (primary visual cortex, early visual cortex, dorsal stream and ventral stream visual cortices, MT+ complex), eDMN (hippocampus, precuneus and posterior cingulate cortices); These regions are implicated for example in visual, perceptual, dorsal (“where”), ventral (“what”) stream visuospatial processing (VNs); cognitive and affective processes such as those involved in, mental imagery, introspection and rumination (eDMN). There was also a paucity of genetically-influenced functional connections involving anterior regions of the brain: Frontoparietal Networks (FPNs) - dorsolateral prefrontal, mid cingulate, insular and frontal opercular, orbital and polar frontal, superior and inferior parietal cortices; Frontotemporal Networks (FTNs) - inferior frontal, posterior opercular, early auditory, auditory association cortices, Sensorimotor networks SMNs - (premotor, somatosensory, paralobular, and motor cortices). These anterior regions are implicated for example in high-level cognitive and affective processes such as working memory, executive function, reasoning, attentional and impulse control, emotional judgement and decision making (FPNs) (Bor et al., 2003; Duncan, 2001; Duncan et al., 2012; Prabhakaran et al., 2001, 2000, 1997); language and auditory processes (FTNs) (Blau et al., 2010; Friederici, 2009; Hernandez, 2009; Hernandez et al., 2001; Hodges et al., 1992; Hommel et al., 2001; Matsumoto et al., 2004; Morillon et al., 2010; Spitsyna et al., 2006; Van Atteveldt et al., 2004), action-planning and movement processes (SMNs) (Andersen and Cui, 2009; Grafton and Hamilton, 2007; Janes et al., 2012; Johnson-Frey et al., 2005; Miller and Cohen, 2001; Penfield and Boldrey, 1937). This result was corroborated by our network analyses which showed a gradient of higher to lesser genetic influence of network-network interaction from posterior to anterior regions respectively: visual > temporal, parietal > frontal networks.

There was a gradient of high to low density of genetically-influenced functional connections involving the posterior to anterior components of the extended Default Mode Network (eDMN) respectively. Specifically, there was a high density of genetically-influenced functional connections involving predominantly posterior-medial regions of eDMN - hippocampus and precuneus/posterior cingulate cortices (Amft et al., 2015). There was a low density of genetically-influenced connections involving anterior (anterior cingulate and medial prefrontal) and lateral (inferior parietal, temporoparietooccipital) regions of the eDMN. The eDMN is involved in low-level cognitive and affective processes such as those involved in episodic memory retrieval, mental imagery, introspection, rumination, evaluation of self and others (Amft et al., 2015; Göttlich et al., 2017).

Other regions that were significantly genetically influenced at the network level for the population included caudate which has been implicated in psychomotor processing speed, procedural learning (Apps et al., 2013); and the hippocampus which has been implicated in long term memory processes (Squire et al., 2004).

#### 4.2.2. Gender-specific Level

The results found at the population level extend to males and females in differing ways.

##### Overall Patterns

There were nearly five times more significant functional connections that are genetically influenced in males (310) than females (64). There were also greater genetically-influenced network-network interactions noted in males than females. These patterns suggest males are more functionally “hard-wired” by genetic influence than females.

##### Cortical regions

As noted at the population level, both males and females were under extensive genetic influence in terms of network interactions involving visual cortices. In addition, males were more genetically-influenced in terms of network interactions involving auditory-language related cortices compared to females. This finding suggests that males may be more functionally “hard-wired” and that females may be more environmentally-influenced and shaped in terms of auditory-language systems than males. This outcome may explain some of the differences in auditory-language system functions reported between males and females (Clements et al., 2006).

As noted at the population level, both males and females were under extensive genetic influence in terms of interactions involving the eDMN which is considered a central hub of the brain for various low-level cognitive and affective processes such as internal monitoring, rumination and evaluation of self and others, as noted previously (Hagmann et al., 2008; Leech and Sharp, 2014)(Ames et al., 2008; Buckner et al., 2008; De Brigard et al., 2015; Denny et al., 2012; Jenkins et al., 2008; Lois and Wessa, 2016; Mitchell et al., 2005; Nejad et al., 2013; Posner et al., 2013; Raposo et al., 2011). In addition, males also were more genetically-influenced compared to females in terms of intranetwork and internetwork interactions between eDMN and other brain regions (occipital, temporal, parietal, and frontal regions) involved in various task-oriented processes and attending to and interacting with the environment which comprise part of the Task Positive Networks (TPNs) (Adolphs, 2006; De Brigard et al., 2015; Denny et al., 2012; Jenkins et al., 2008; Mahy et al., 2014; Raposo et al., 2011; Saxe, 2006; Schurz and Perner, 2015).

There were also nearly five times more genetically influenced functional connections in males (310) than females (64) suggesting that male brains are more genetically influenced, i.e. functionally “hard-wired”, than females. This result suggests differences in genetic predisposition in males (more) vs. females (less) in terms of interplay of attending to task-oriented interactions with the environment (TPNs) vs. internal and external interactions with self and others (eDMN). This outcome may also have implications in terms of brain plasticity differences in males (less) versus females (more) in terms of ability to react or adapt/maladapt to environmental influences (e.g. task completion demands, psychosocial stressors, positive and negative feedback, meditation, cognitive behavioral therapy, pharmacotherapy) and their overall malleability.

##### Subcortical regions

Thalamus and basal ganglia (caudate) regions demonstrated a significant number of connections with each other as well as cortical regions in males which may be due to the cortico-basal-ganglia-thalamic-cortical feedback loops needed to maintain or sustain activity in these lateral cortical regions involved in various high-level and low-level cognitive processes (Alexander, 1986). Ventral diencephalon showed significant connections in males which control autonomic functions. Cortico-Cerebellar involvement was noted both in males and females which are involved in processes such as providing error correction and feedback to cortical processes (Habas et al., 2009).

### 4.3. Genetics and Environmental Factors

Fu et al. demonstrated in a pediatric population using rsfMRI (resting state eyes closed) data greater heritability among the posterior (visual), and lesser in the anterior brain regions (auditory, attention, executive control) resting state networks (Fu et al., 2015), which is consistent with our results that posterior compared to anterior regions are more genetically influenced. A study examining heritability in the HCP cohort involving twins, as well as nontwin siblings with rsfMRI data found low heritability of 20-40% throughout the brain including anterior and posterior brain regions (Adhikari et al., 2018). This is consistent with our findings which finds only 9/32 networks demonstrating significant genetic influence at the overall population level, although somewhat difficult to compare given that genetics and environmental factors as well as twins and nontwin siblings were taken into account in how heritability was calculated in their study, versus our approach which examines mainly the influence of genetics in twins exclusively.

Task fMRI studies have implicated brain regions involved in certain tasks as heritable (Jansen et al., 2015). One study investigating heritability of signal in the N-back fMRI task found heritability specific to the left supplementary motor area, inferior, middle, and superior frontal gyri, precentral and postcentral gyri, middle cingulate cortex, superior medial gyrus, angular gyrus, and the superior parietal lobule (Blokland et al., 2011). Another study examined the shared genetic etiology between cognitive performance and brain activations in language and math tasks using the HCP dataset. They found several parts of the language network along the superior temporal sulcus, as well as the angular gyrus belonging to the math processing network, are significantly genetically-correlated with these indicators of cognitive performance (Le Guen et al., 2018). The differences in heritability/genetic influence results between rsfMRI datasets (Adhikari et al., 2018; Elliott et al., 2019) and other results which use task fMRI datasets may be due to the differences in what is being assessed by resting state vs. task fMRI (Koten et al., 2009). Resting state fMRI data is commonly utilized to assess the functional connectivity of the brain regions during rest, while task fMRI data is commonly utilized to assess the activation of brain regions while performing a task. Moreover, factors such as the amount of rsfMRI data collected or the scan length also appear to influence the amount of variance in rsFC attributable to genetic influence (Elliott et al., 2019). Additionally, the specific parcellation used to define regions of interest, as well as primarily examining connections at the region level and not the connection level may lead to differences in our results. Overall, however, rsfMRI and task fMRI are measuring two different aspects of brain function or state which may show differences in heritability or genetic influence.

Our results suggest that rsFC involving anterior brain regions implicated in high-level cognitive and affective processes such as working memory, executive function, reasoning, attentional and impulse control, decision making and emotional judgements (FPNs); language and auditory processes (FTNs); action planning and movement (SMNs - premotor, somatosensory and motor regions) may have both environmental and genetic input which is suggested by more recent work (Fu et al., 2015) while rsFC involving posterior regions implicated in visual, perceptual, low-level cognitive and affective processes may be influenced more by genetics. Additionally, since the results for the connection analyses are Bonferroni corrected, they serve as robust findings that may not otherwise be found in typical ACE modelling approaches. These connections may in turn serve as specific endophenotypes in young, healthy individuals. Furthermore, these connections may also have implications in aging and development as well as in disease and other disorders.

### 4.4 Limitations

This work was limited by several factors. First, to determine genetic influence for the method proposed, it was assumed that the phenotypic difference in variances between MZ and DZ twin groups was primarily attributable to genetics. In other words, it was assumed that, on average, the shared and unshared environmental effects on the brain were similar between the MZ and DZ groups. Additionally, the data was limited to rsfMRI which focuses on functional connectivity involving various brain regions. Further analyses of other data types (e.g. task fMRI, DTI, and morphometrics) which have shown involvement of genetics and environment on other aspects of brain function and structure (Jansen et al., 2015; Jin et al., 2011) need to be investigated in conjunction with our method. The use of the F-Test is highly sensitive to non-normally distributed data, and therefore even slight deviations from normality may have influenced the resulting variances. Another limitation is that given the multitude of methods possible for normalizing the connections for the network-network interaction summarization, the raw data was discussed.

Although we are aware of correcting for test statistics in twin double entering methods such as in ACE model studies, in this study we have not implemented this correction since we did not make any assumptions about the direction of genetic influence on MZ compared to DZ twin groups. An additional limitation of this study is the low sample size due to education, age and gender matching. Future studies should investigate the effect of sample size on the results. We also did not incorporate other siblings or related family members in the analyses to minimize assumptions about genetics and environment. While there was a difference in age between males and females, all subjects were young healthy adults and brain maturation is thought to have plateaued at this time resulting in no significant changes in brain plasticity or connectivity during this age range. Furthermore, the direct statistical comparisons that were examined were between the DZ and MZ groups with no significant age difference in these groups, which would result in removal of any age effects.

## 5. Conclusions

Overall, our approach allowed for a parsimonious determination of genetic influence at the population and gender-specific levels. The method highlighted genetically-dependent resting state connections particularly across visual, parietal, temporal, and posterior cingulate cortices (i.e. visual, ventral stream, and dorsal stream regions, MT). These posterior brain regions which are more genetically influenced are implicated in visual, perceptual, low-level cognitive and affective processes. There was also a paucity of connections involving anterior regions of the brain which are less genetically-influenced, i.e., frontoparietal cortices, which are implicated in high-level cognitive and affective processes such as working memory, executive function, reasoning, attentional and impulse control, emotional judgement and decision making; frontotemporal cortices which are implicated in language and auditory processes; and sensorimotor cortices which are implicated for example in action planning and movement processes. These regional differences in genetic influence may have implications in terms of the brain”s ability to be influenced by environmental input (e.g. education, school and work environment, family and home environment, social interaction with peers, medications, nutrition, sports and physical exercise).

There were nearly five times more connections that are genetically influenced in males than females suggesting that males are more functionally “hard-wired” than females. There were also greater intra and internetwork interactions between TPNs and eDMN in males than females, which may lead to genetic predisposition of males compared to females being functionally “hardwired” in terms of interplay of interaction with environment and task-oriented processes vs. interaction with self and others. This observation may have implications in terms of brain plasticity differences in males (low) versus females (high) in terms of ability to react or adapt/maladapt to environmental influences (e.g. psychosocial stressors, positive feedback, meditation, cognitive behavioral therapy, pharmacotherapy). These results reveal the similarities and differences of genetics and environmental influences on different parcels, connections, and networks of the resting state functional brain in young healthy males and females with implications in development and later in life.

## Acknowledgments

The authors would like to thank all the participants involved in the Human Connectome Project and Washington University in St. Louis for data collection and grant support from NIH UF1AG051216, R01NS105646, and R01EB027087. The research presented was supported under NIH award TL1TR002375. The content is solely the responsibility of the authors and does not necessarily represent the official views of the National Institutes of Health.

