## Supplemental Information for "Genetic influence on resting state networks in young male and female adults"

Supplementary Information (SI) Follows

| Hemisphere Type | Network Interaction | Parcel Interaction | p-value |
|---|---|---|---|
| Intra (Left) | MT+ Complex and Neighbors -- MT+ Complex and Neighbors | Left Area PH -- Left Area V3CD | 1.59E-11 |
| Inter | MT+ Complex and Neighbors -- MT+ Complex and Neighbors | Left Area PH -- Right Area V3CD | 1.67E-09 |
| Intra (Right) | Ventral Stream -- Dorsal Stream | Right VentroMedial Visual Area 1 -- Right Area V3A | 1.12E-07 |
| Inter | MT+ Complex and Neighbors -- Dorsal Stream | Left Area PH -- Right IntraParietal Sulcus Area 1 | 3.59E-07 |
| Inter | MT+ Complex and Neighbors -- Dorsal Stream | Right Area Lateral Occipital 1 -- Left IntraParietal Sulcus Area 1 | 4.17E-07 |
| Intra (Left) | Inferior Frontal Cortex -- Inferior Parietal Cortex | Left Area 47l (47 lateral) -- Left Area PGi | 6.56E-07 |
| Intra (Right) | Ventral Stream -- Dorsal Stream | Right VentroMedial Visual Area 2 -- Right IntraParietal Sulcus Area 1 | 1.08E-06 |
| Inter | Inferior Parietal Cortex -- Medial Temporal Cortex | Left Area IntraParietal 0 -- Right ParaHippocampal Area 3 | 1.32E-06 |
| Intra (Right) | Posterior Cingulate Cortex -- Inferior Parietal Cortex | Right Area dorsal 23 a+b -- Right Area IntraParietal 1 | 2.22E-06 |
| Inter | Posterior Cingulate Cortex -- Posterior Cingulate Cortex | Left PreCuneus Visual Area -- Right ProStriate Area | 2.25E-06 |
| Inter | Ventral Stream -- Ventral Stream | Left VentroMedial Visual Area 2 -- Right VentroMedial Visual Area 1 | 4.63E-06 |
| Intra (Right) | Superior Parietal and IPS Cortex -- Dorsal Stream | Right Area Lateral IntraParietal ventral -- Right IntraParietal Sulcus Area 1 | 5.57E-06 |
| Inter | Inferior Parietal Cortex -- Sensorimotor Associated Paracentral Lobular and Mid Cingulate Cortex | Right Area PF opercular -- Left Area 5m ventral | 6.06E-06 |
| Inter | Ventral Stream -- Primary Visual Cortex (V1) | Left VentroMedial Visual Area 2 -- Right Primary Visual Cortex | 6.99E-06 |
| Intra (Right) | Temporal-Parietal-Occipital Junction -- Auditory Association Cortex | Right Area TemporoParietoOccipital Junction 2 -- Right Area STGa | 8.89E-06 |
| Inter | Posterior Cingulate Cortex -- Posterior Cingulate Cortex | Left ProStriate Area -- Right PreCuneus Visual Area | 9.78E-06 |
| Inter | Posterior Cingulate Cortex -- Ventral Stream | Right PreCuneus Visual Area -- Left Ventral Visual Complex | 1.15E-05 |
| Intra (Left) | MT+ Complex and Neighbors -- Dorsal Stream | Left Area Lateral Occipital 1 -- Left IntraParietal Sulcus Area 1 | 1.26E-05 |
| Intra (Left) | Posterior Cingulate Cortex -- Ventral Stream | Left PreCuneus Visual Area -- Left VentroMedial Visual Area 2 | 1.37E-05 |
| Intra (Left) | Posterior Cingulate Cortex -- Inferior Parietal Cortex | Left Dorsal Transitional Visual Area -- Left Area IntraParietal 0 | 1.57E-05 |
| Inter | MT+ Complex and Neighbors -- Dorsal Stream | Right Area PH -- Left IntraParietal Sulcus Area 1 | 1.80E-05 |
| Intra (Left) | Posterior Cingulate Cortex -- Superior Parietal and IPS Cortex | Left Area 31a -- Left Medial Area 7P | 1.89E-05 |
| Intra (Right) | Superior Parietal and IPS Cortex -- MT+ Complex and Neighbors | Right Area Lateral IntraParietal ventral -- Right Area PH | 1.92E-05 |
| Intra (Right) | Posterior Cingulate Cortex -- Inferior Parietal Cortex | Right Area ventral 23 a+b -- Right Area IntraParietal 1 | 2.75E-05 |
| Intra (Left) | Posterior Cingulate Cortex -- Ventral Stream | Left Dorsal Transitional Visual Area -- Left VentroMedial Visual Area 2 | 3.13E-05 |
| Intra (Left) | Ventral Stream -- Primary Visual Cortex (V1) | Left VentroMedial Visual Area 2 -- Left Primary Visual Cortex | 3.87E-05 |
| Inter | Sensorimotor Associated Paracentral Lobular and Mid Cingulate Cortex -- Somatosensory and Motor Cortex | Left Area 5m -- Right Primary Sensory Cortex | 4.13E-05 |
| Intra (Left) | Medial Temporal Cortex -- Dorsal Stream | Left ParaHippocampal Area 3 -- Left Area V3A | 4.43E-05 |
| Inter | Inferior Parietal Cortex -- Sensorimotor Associated Paracentral Lobular and Mid Cingulate Cortex | Left Area IntraParietal 0 -- Right Area 5m ventral | 4.43E-05 |
| Inter | Ventral Stream -- Dorsal Stream | Right VentroMedial Visual Area 1 -- Left Area V3A | 4.47E-05 |
| Intra (Right) | Inferior Frontal Cortex -- Inferior Frontal Cortex | Right Area 44 -- Right Area 45 | 4.50E-05 |
| Inter | Posterior Cingulate Cortex -- Superior Parietal and IPS Cortex | Right PreCuneus Visual Area -- Left Area Lateral IntraParietal dorsal | 4.89E-05 |
| Intra (Right) | Posterior Cingulate Cortex -- Inferior Parietal Cortex | Right Area dorsal 23 a+b -- Right Area IntraParietal 2 | 4.95E-05 |
| Inter | Posterior Cingulate Cortex -- Superior Parietal and IPS Cortex | Left ProStriate Area -- Right Area Lateral IntraParietal dorsal | 5.84E-05 |
| Intra (Left) | Posterior Cingulate Cortex -- Posterior Cingulate Cortex | Left PreCuneus Visual Area -- Left Dorsal Transitional Visual Area | 7.35E-05 |
| Inter | Ventral Stream -- Dorsal Stream | Right VentroMedial Visual Area 2 -- Left IntraParietal Sulcus Area 1 | 7.74E-05 |
| Intra (Left) | Posterior Opercular Cortex -- MT+ Complex and Neighbors | Left Area OP1/SII -- Left Middle Temporal Area | 8.13E-05 |
| Intra (Right) | Posterior Cingulate Cortex -- Ventral Stream | Right Dorsal Transitional Visual Area -- Right VentroMedial Visual Area 2 | 8.38E-05 |
| Intra (Right) | Inferior Frontal Cortex -- Posterior Cingulate Cortex | Right Area IFJp -- Right Area dorsal 23 a+b | 8.81E-05 |
| Intra (Left) | Sensorimotor Associated Paracentral Lobular and Mid Cingulate Cortex -- Somatosensory and Motor Cortex | Left Area 5m -- Left Primary Sensory Cortex | 9.08E-05 |
| Intra (Left) | Ventral Stream -- Dorsal Stream | Left VentroMedial Visual Area 2 -- Left Area V3A | 9.48E-05 |
| Intra (Left) | Posterior Cingulate Cortex -- Inferior Parietal Cortex | Left Area dorsal 23 a+b -- Left Area PGs | 9.54E-05 |
| Inter | Posterior Cingulate Cortex -- Medial Temporal Cortex | Right PreCuneus Visual Area -- Left ParaHippocampal Area 1 | 9.84E-05 |
| Intra (Right) | Caudate -- Posterior Cingulate Cortex | Right Caudate -- Right Parieto-Occipital Sulcus Area 2 | 9.84E-05 |
| Inter | Posterior Cingulate Cortex -- Inferior Parietal Cortex | Right Area ventral 23 a+b -- Left Area PGs | 0.00010409 |
| Intra (Right) | Ventral Stream -- Dorsal Stream | Right VentroMedial Visual Area 2 -- Right Area V3A | 0.000111815 |
| Intra (Right) | Posterior Cingulate Cortex -- Posterior Cingulate Cortex | Right PreCuneus Visual Area -- Right ProStriate Area | 0.000117273 |
| Inter | Ventral Stream -- Dorsal Stream | Right VentroMedial Visual Area 2 -- Left Sixth Visual Area | 0.00012523 |
| Intra (Right) | MT+ Complex and Neighbors -- Dorsal Stream | Right Area Lateral Occipital 1 -- Right Seventh Visual Area | 0.000129465 |
| Intra (Right) | Posterior Cingulate Cortex -- Early Visual Cortex | Right PreCuneus Visual Area -- Right Second Visual Area | 0.000134724 |
| Inter | Ventral Stream -- Ventral Stream | Left VentroMedial Visual Area 2 -- Right VentroMedial Visual Area 2 | 0.000149828 |
| Intra (Left) | Posterior Cingulate Cortex -- Temporal-Parietal-Occipital Junction | Left Dorsal Transitional Visual Area -- Left Area TemporoParietoOccipital Junction 1 | 0.000152978 |
| Intra (Left) | Ventral Stream -- Dorsal Stream | Left VentroMedial Visual Area 2 -- Left Area V3B | 0.000164691 |
| Inter | Posterior Cingulate Cortex -- Ventral Stream | Right PreCuneus Visual Area -- Left VentroMedial Visual Area 2 | 0.000194516 |
| Inter | Inferior Parietal Cortex -- Superior Parietal and IPS Cortex | Right Area PFt -- Left Medial IntraParietal Area | 0.000198113 |
| Intra (Right) | Posterior Cingulate Cortex -- Ventral Stream | Right PreCuneus Visual Area -- Right Fusiform Face Complex | 0.000200418 |
| Intra (Left) | Dorsal Stream -- Dorsal Stream | Left Seventh Visual Area -- Left IntraParietal Sulcus Area 1 | 0.000209226 |
| Inter | MT+ Complex and Neighbors -- Dorsal Stream | Left Area PH -- Right Area V3B | 0.000230166 |
| Inter | Inferior Parietal Cortex -- Lateral Temporal Cortex | Left Area IntraParietal 0 -- Right Area TE2 posterior | 0.00023231 |
| Inter | Ventral Stream -- Early Visual Cortex | Left VentroMedial Visual Area 2 -- Right Second Visual Area | 0.000246649 |
| Inter | Sensorimotor Associated Paracentral Lobular and Mid Cingulate Cortex -- Somatosensory and Motor Cortex | Left Area 5m -- Right Area 1 | 0.0002555 |
| Inter | Ventral Stream -- Dorsal Stream | Left VentroMedial Visual Area 2 -- Right Area V3A | 0.000273365 |
| Intra (Left) | Sensorimotor Associated Paracentral Lobular and Mid Cingulate Cortex -- Dorsal Stream | Left Area 5m -- Left Sixth Visual Area | 0.000276115 |
| Intra (Right) | Caudate -- Superior Parietal and IPS Cortex | Right Caudate -- Right Area Lateral IntraParietal dorsal | 0.000279406 |
| Intra (Right) | MT+ Complex and Neighbors -- Ventral Stream | Right Area Lateral Occipital 3 -- Right VentroMedial Visual Area 2 | 0.000282563 |
| Inter | Posterior Cingulate Cortex -- Superior Parietal and IPS Cortex | Right Area ventral 23 a+b -- Left Area Lateral IntraParietal dorsal | 0.000285529 |
| Intra (Right) | Inferior Frontal Cortex -- Posterior Cingulate Cortex | Right Area IFJp -- Right Area ventral 23 a+b | 0.000295256 |
| Inter | Ventral Stream -- Dorsal Stream | Right VentroMedial Visual Area 2 -- Left Area V3A | 0.000296662 |
| Intra (Left) | Ventral Stream -- Early Visual Cortex | Left Ventral Visual Complex -- Left Fourth Visual Area | 0.000299044 |
| Intra (Right) | Auditory Association Cortex -- Early Auditory Cortex | Right Area STGa -- Right Lateral Belt Complex | 0.000306856 |
| Intra (Right) | MT+ Complex and Neighbors -- Ventral Stream | Right Area PH -- Right VentroMedial Visual Area 2 | 0.000313668 |
| Inter | Inferior Frontal Cortex -- Inferior Parietal Cortex | Right Area 47l (47 lateral) -- Left Area IntraParietal 1 | 0.000316602 |
| Intra (Right) | Posterior Cingulate Cortex -- Ventral Stream | Right PreCuneus Visual Area -- Right Eighth Visual Area | 0.00032241 |
| Intra (Left) | Auditory Association Cortex -- Auditory Association Cortex | Left Area STSv posterior -- Left Area STSv anterior | 0.000324346 |
| Intra (Right) | Inferior Frontal Cortex -- Inferior Parietal Cortex | Right Area 47l (47 lateral) -- Right Area IntraParietal 1 | 0.000347919 |
| Inter | Insular and Frontal Opercular Cortex -- Sensorimotor Associated Paracentral Lobular and Mid Cingulate Cortex | Right Area Posterior Insular 1 -- Left Area 5m | 0.000366198 |
| Intra (Left) | Cerebellum -- Medial Temporal Cortex | Left Cerebellum -- Left ParaHippocampal Area 2 | 0.000397787 |
| Intra (Left) | Orbital and Polar Frontal Cortex -- Inferior Parietal Cortex | Left Area 13l -- Left Area PFm Complex | 0.000415172 |
| Inter | Posterior Cingulate Cortex -- Ventral Stream | Right Dorsal Transitional Visual Area -- Left VentroMedial Visual Area 2 | 0.000416222 |
| Inter | Ventral Stream -- Dorsal Stream | Right Eighth Visual Area -- Left IntraParietal Sulcus Area 1 | 0.000425409 |
| Inter | Hippocampus -- Posterior Opercular Cortex | Left Subcortical Hippocampus -- Right Area PFcm | 0.000441677 |
| Inter | Posterior Cingulate Cortex -- Inferior Parietal Cortex | Left Area 23c -- Right Area PF opercular | 0.000522828 |
| Inter | Medial Temporal Cortex -- Dorsal Stream | Left ParaHippocampal Area 3 -- Right Area V3B | 0.000529744 |
| Inter | Posterior Cingulate Cortex -- Early Visual Cortex | Right PreCuneus Visual Area -- Left Second Visual Area | 0.000533698 |
| Inter | Ventral Stream -- Primary Visual Cortex (V1) | Right VentroMedial Visual Area 2 -- Left Primary Visual Cortex | 0.000548093 |
| Inter | Posterior Cingulate Cortex -- Primary Visual Cortex (V1) | Right PreCuneus Visual Area -- Left Primary Visual Cortex | 0.000553327 |
| Intra (Left) | Inferior Frontal Cortex -- Anterior Cingulate and Medial Prefrontal Cortex | Left Area 47l (47 lateral) -- Left Area 9 Middle | 0.000556658 |
| Inter | MT+ Complex and Neighbors -- Ventral Stream | Left Area PH -- Right Ventral Visual Complex | 0.000578591 |
| Intra (Left) | Posterior Cingulate Cortex -- Medial Temporal Cortex | Left Dorsal Transitional Visual Area -- Left ParaHippocampal Area 1 | 0.000585635 |
| Inter | Superior Parietal and IPS Cortex -- Dorsal Stream | Right Area Lateral IntraParietal ventral -- Left IntraParietal Sulcus Area 1 | 0.00058599 |
| Intra (Left) | Posterior Cingulate Cortex -- Lateral Temporal Cortex | Left PreCuneus Visual Area -- Left Area TE2 posterior | 0.000594371 |
| Inter | Ventral Stream -- Dorsal Stream | Left VentroMedial Visual Area 2 -- Right Area V3B | 0.000599816 |
| Intra (Left) | Posterior Cingulate Cortex -- Ventral Stream | Left PreCuneus Visual Area -- Left Fusiform Face Complex | 0.000600085 |
| Inter | Anterior Cingulate and Medial Prefrontal Cortex -- Posterior Cingulate Cortex | Left Area 25 -- Right Area ventral 23 a+b | 0.000601965 |
| Inter | Ventral Stream -- Early Visual Cortex | Right VentroMedial Visual Area 2 -- Left Second Visual Area | 0.000605223 |
| Intra (Left) | MT+ Complex and Neighbors -- Early Visual Cortex | Left Area PH -- Left Fourth Visual Area | 0.000624431 |
| Intra (Right) | Inferior Frontal Cortex -- Posterior Cingulate Cortex | Right Area IFJp -- Right Parieto-Occipital Sulcus Area 1 | 0.000629959 |
| Intra (Right) | Superior Parietal and IPS Cortex -- Ventral Stream | Right Anterior IntraParietal Area -- Right VentroMedial Visual Area 2 | 0.000633516 |
| Intra (Right) | Caudate -- Ventral Stream | Right Caudate -- Right Ventral Visual Complex | 0.000637765 |
| Intra (Right) | Auditory Association Cortex -- Posterior Opercular Cortex | Right Area STSd posterior -- Right Area PFcm | 0.000663134 |
| Intra (Left) | Posterior Cingulate Cortex -- Premotor Cortex | Left PreCuneus Visual Area -- Left Frontal Eye Fields | 0.000677928 |
| Inter | Medial Temporal Cortex -- Primary Visual Cortex (V1) | Left ParaHippocampal Area 3 -- Right Primary Visual Cortex | 0.00075205 |
| Intra (Left) | Dorsal Stream -- Dorsal Stream | Left Area V3A -- Left IntraParietal Sulcus Area 1 | 0.000754428 |
| Inter | Cerebellum -- Orbital and Polar Frontal Cortex | Left Cerebellum -- Right Area posterior 10p | 0.000755239 |
| Intra (Left) | Superior Parietal and IPS Cortex -- MT+ Complex and Neighbors | Left Medial IntraParietal Area -- Left Area FST | 0.00081958 |
| Inter | Medial Temporal Cortex -- Sensorimotor Associated Paracentral Lobular and Mid Cingulate Cortex | Right Entorhinal Cortex -- Left Area 5m | 0.000823949 |
| Intra (Right) | Inferior Frontal Cortex -- Anterior Cingulate and Medial Prefrontal Cortex | Right Area 45 -- Right Area p32 | 0.000827409 |
| Inter | Posterior Cingulate Cortex -- Medial Temporal Cortex | Right PreCuneus Visual Area -- Left ParaHippocampal Area 2 | 0.000830415 |
| Intra (Left) | Inferior Parietal Cortex -- Superior Parietal and IPS Cortex | Left Area IntraParietal 1 -- Left Medial Area 7P | 0.00084532 |
| Inter | Medial Temporal Cortex -- Early Visual Cortex | Left ParaHippocampal Area 3 -- Right Second Visual Area | 0.000876294 |
| Intra (Right) | MT+ Complex and Neighbors -- MT+ Complex and Neighbors | Right Area Lateral Occipital 1 -- Right Area FST | 0.000878325 |
| Intra (Right) | Posterior Cingulate Cortex -- Insular and Frontal Opercular Cortex | Right Area 31pd -- Right Area Frontal Opercular 5 | 0.000888833 |
| Intra (Left) | Posterior Cingulate Cortex -- Inferior Parietal Cortex | Left PreCuneus Visual Area -- Left Area IntraParietal 0 | 0.000917799 |
| Intra (Right) | Posterior Cingulate Cortex -- Ventral Stream | Right PreCuneus Visual Area -- Right Ventral Visual Complex | 0.000931411 |
| Intra (Right) | Dorsolateral Prefrontal Cortex -- Anterior Cingulate and Medial Prefrontal Cortex | Right Area anterior 9-46v -- Right Area dorsal 32 | 0.000939089 |
| Intra (Left) | MT+ Complex and Neighbors -- Dorsal Stream | Left Area PH -- Left Seventh Visual Area | 0.000958205 |
| Intra (Right) | MT+ Complex and Neighbors -- Ventral Stream | Right Area Lateral Occipital 3 -- Right VentroMedial Visual Area 1 | 0.000975496 |
| Intra (Left) | Medial Temporal Cortex -- Early Visual Cortex | Left ParaHippocampal Area 3 -- Left Second Visual Area | 0.001005897 |
| Intra (Right) | MT+ Complex and Neighbors -- Dorsal Stream | Right Area PH -- Right IntraParietal Sulcus Area 1 | 0.001017105 |
| Inter | Posterior Cingulate Cortex -- Temporal-Parietal-Occipital Junction | Right PreCuneus Visual Area -- Left Superior Temporal Visual Area | 0.001074371 |
| Intra (Right) | Ventral Stream -- Dorsal Stream | Right VentroMedial Visual Area 2 -- Right Area V6A | 0.001117259 |
| Inter | MT+ Complex and Neighbors -- Dorsal Stream | Left Area PH -- Right Seventh Visual Area | 0.001123328 |
| Intra (Right) | Caudate -- Lateral Temporal Cortex | Right Caudate -- Right Area TE2 posterior | 0.001139336 |
| Intra (Left) | Superior Parietal and IPS Cortex -- MT+ Complex and Neighbors | Left Medial IntraParietal Area -- Left Area PH | 0.001158475 |
| Intra (Left) | MT+ Complex and Neighbors -- Dorsal Stream | Left Area FST -- Left Seventh Visual Area | 0.001202207 |
| Intra (Right) | Posterior Cingulate Cortex -- Medial Temporal Cortex | Right Area 31pd -- Right Entorhinal Cortex | 0.001206839 |
| Intra (Left) | Orbital and Polar Frontal Cortex -- Anterior Cingulate and Medial Prefrontal Cortex | Left Posterior OFC Complex -- Left Area 33 prime | 0.001229299 |
| Intra (Right) | Caudate -- MT+ Complex and Neighbors | Right Caudate -- Right Area PH | 0.001233124 |
| Inter | Posterior Cingulate Cortex -- Temporal-Parietal-Occipital Junction | Left PreCuneus Visual Area -- Right Area TemporoParietoOccipital Junction 3 | 0.001243974 |
| Inter | Ventral Stream -- Dorsal Stream | Right Ventral Visual Complex -- Left IntraParietal Sulcus Area 1 | 0.001248491 |
| Inter | Medial Temporal Cortex -- Dorsal Stream | Left ParaHippocampal Area 3 -- Right Area V3A | 0.001260516 |
| Inter | MT+ Complex and Neighbors -- Ventral Stream | Left Area PH -- Right VentroMedial Visual Area 3 | 0.001265687 |
| Intra (Left) | MT+ Complex and Neighbors -- Dorsal Stream | Left Area PH -- Left Area V3A | 0.001266948 |
| Inter | MT+ Complex and Neighbors -- Dorsal Stream | Right Area PH -- Left Area V3B | 0.001269996 |
| Inter | Superior Parietal and IPS Cortex -- Early Visual Cortex | Right Area Lateral IntraParietal ventral -- Left Fourth Visual Area | 0.001273737 |
| Inter | Sensorimotor Associated Paracentral Lobular and Mid Cingulate Cortex -- Somatosensory and Motor Cortex | Left Area 5m -- Right Area 2 | 0.001274153 |
| Inter | Inferior Frontal Cortex -- Posterior Cingulate Cortex | Right Area IFJp -- Left Parieto-Occipital Sulcus Area 1 | 0.001279014 |
| Intra (Right) | Caudate -- Ventral Stream | Right Caudate -- Right Fusiform Face Complex | 0.001288434 |
| Inter | Superior Parietal and IPS Cortex -- Ventral Stream | Right Area Lateral IntraParietal dorsal -- Left VentroMedial Visual Area 2 | 0.001299249 |
| Intra (Left) | Posterior Cingulate Cortex -- Temporal-Parietal-Occipital Junction | Left PreCuneus Visual Area -- Left Area TemporoParietoOccipital Junction 3 | 0.001369854 |
| Intra (Right) | Temporal-Parietal-Occipital Junction -- Ventral Stream | Right Superior Temporal Visual Area -- Right Ventral Visual Complex | 0.001497488 |
| Intra (Left) | Posterior Cingulate Cortex -- Posterior Cingulate Cortex | Left Area 23d -- Left Area 31pd | 0.001510046 |

| | | | |
|---|---|---|---|
| Intra (Left) | MT+ Complex and Neighbors -- Ventral Stream | Left Area V3CD -- Left Ventral Visual Complex | 0.001517711 |
| Intra (Left) | Medial Temporal Cortex -- Ventral Stream | Left ParaHippocampal Area 3 -- Left Eighth Visual Area | 0.001524909 |
| Intra (Left) | Orbital and Polar Frontal Cortex -- Inferior Parietal Cortex | Left Area anterior 47r -- Left Area PFm Complex | 0.001551416 |
| Intra (Right) | Posterior Cingulate Cortex -- Primary Visual Cortex (V1) | Right PreCuneus Visual Area -- Right Primary Visual Cortex | 0.001564277 |
| Intra (Right) | Hippocampus -- Posterior Opercular Cortex | Right Subcortical Hippocampus -- Right Area PFcm | 0.001579525 |
| Intra (Right) | Posterior Cingulate Cortex -- MT+ Complex and Neighbors | Right PreCuneus Visual Area -- Right Area Lateral Occipital 3 | 0.001623046 |
| Inter | Posterior Cingulate Cortex -- Temporal-Parietal-Occipital Junction | Right PreCuneus Visual Area -- Left Area TemporoParietoOccipital Junction 3 | 0.001645855 |
| Inter | Medial Temporal Cortex -- Sensorimotor Associated Paracentral Lobular and Mid Cingulate Cortex | Left PreSubiculum -- Right Area 5m ventral | 0.001691364 |
| Inter | Sensorimotor Associated Paracentral Lobular and Mid Cingulate Cortex -- MT+ Complex and Neighbors | Right Area 5m ventral -- Left Area PH | 0.001744974 |
| Intra (Right) | Posterior Cingulate Cortex -- Inferior Parietal Cortex | Right Area ventral 23 a+b -- Right Area IntraParietal 2 | 0.001770335 |
| Inter | Superior Parietal and IPS Cortex -- Sensorimotor Associated Paracentral Lobular and Mid Cingulate Cortex | Right Area Lateral IntraParietal dorsal -- Left Supplementary and Cingulate Eye Field | 0.001801392 |
| Intra (Left) | Dorsolateral Prefrontal Cortex -- Inferior Parietal Cortex | Left Area 9 Posterior -- Left Area PGs | 0.001802238 |
| Intra (Right) | Dorsolateral Prefrontal Cortex -- Inferior Frontal Cortex | Right Area 9 anterior -- Right Area 45 | 0.00181193 |
| Intra (Right) | Cerebellum -- Ventral Stream | Right Cerebellum -- Right VentroMedial Visual Area 3 | 0.001876735 |
| Inter | MT+ Complex and Neighbors -- Ventral Stream | Left Area PH -- Right VentroMedial Visual Area 2 | 0.001905823 |
| Inter | Ventral Stream -- Primary Visual Cortex (V1) | Right Ventral Visual Complex -- Left Primary Visual Cortex | 0.001938307 |
| Inter | Cerebellum -- Medial Temporal Cortex | Right Cerebellum -- Left ParaHippocampal Area 2 | 0.001970171 |
| Inter | Insular and Frontal Opercular Cortex -- Insular and Frontal Opercular Cortex | Left Insular Granular Complex -- Right Area Posterior Insular 1 | 0.001988764 |
| Intra (Left) | Medial Temporal Cortex -- Early Visual Cortex | Left ParaHippocampal Area 3 -- Left Fourth Visual Area | 0.00200505 |
| Intra (Right) | Inferior Frontal Cortex -- Posterior Cingulate Cortex | Right Area 47l (47 lateral) -- Right Area ventral 23 a+b | 0.002024861 |
| Intra (Left) | Hippocampus -- Sensorimotor Associated Paracentral Lobular and Mid Cingulate Cortex | Left Subcortical Hippocampus -- Left Area 5m | 0.002031811 |
| Intra (Left) | Medial Temporal Cortex -- Primary Visual Cortex (V1) | Left ParaHippocampal Area 3 -- Left Primary Visual Cortex | 0.002042228 |
| Intra (Left) | Ventral Stream -- Dorsal Stream | Left Ventral Visual Complex -- Left Area V3B | 0.002084147 |
| Intra (Right) | Ventral Diencephalon -- Auditory Association Cortex | Right VentralDiencephalon -- Right Area STSv posterior | 0.002144867 |
| Intra (Left) | Superior Parietal and IPS Cortex -- Dorsal Stream | Left Ventral IntraParietal Complex -- Left IntraParietal Sulcus Area 1 | 0.002166928 |
| Intra (Left) | Superior Parietal and IPS Cortex -- Temporal-Parietal-Occipital Junction | Left Lateral Area 7P -- Left Area TemporoParietoOccipital Junction 3 | 0.002228846 |
| Intra (Left) | Insular and Frontal Opercular Cortex -- Premotor Cortex | Left Area Frontal Opercular 5 -- Left Area 55b | 0.002249149 |
| Inter | MT+ Complex and Neighbors -- MT+ Complex and Neighbors | Left Area PH -- Right Area Lateral Occipital 1 | 0.002264469 |
| Inter | Ventral Diencephalon -- MT+ Complex and Neighbors | Right VentralDiencephalon -- Left Area Lateral Occipital 1 | 0.002294865 |
| Inter | Sensorimotor Associated Paracentral Lobular and Mid Cingulate Cortex -- Dorsal Stream | Left Area 5m -- Right Area V6A | 0.0023179 |
| Intra (Left) | Posterior Cingulate Cortex -- Insular and Frontal Opercular Cortex | Left Area 23c -- Left Area Posterior Insular 1 | 0.002347087 |
| Intra (Right) | Temporal-Parietal-Occipital Junction -- MT+ Complex and Neighbors | Right Area TemporoParietoOccipital Junction 2 -- Right Area FST | 0.002394636 |
| Intra (Right) | Hippocampus -- Superior Parietal and IPS Cortex | Right Subcortical Hippocampus -- Right Lateral Area 7A | 0.002416997 |
| Intra (Right) | Ventral Stream -- Early Visual Cortex | Right VentroMedial Visual Area 2 -- Right Second Visual Area | 0.00243113 |
| Inter | Posterior Cingulate Cortex -- Posterior Cingulate Cortex | Left PreCuneus Visual Area -- Right PreCuneus Visual Area | 0.00247636 |
| Intra (Right) | Posterior Cingulate Cortex -- Lateral Temporal Cortex | Right Parieto-Occipital Sulcus Area 1 -- Right Area TF | 0.002503703 |
| Inter | MT+ Complex and Neighbors -- Dorsal Stream | Right Area V3CD -- Left IntraParietal Sulcus Area 1 | 0.002539345 |
| Inter | Posterior Cingulate Cortex -- Posterior Cingulate Cortex | Left Parieto-Occipital Sulcus Area 1 -- Right Area dorsal 23 a+b | 0.0026052 |
| Intra (Right) | Hippocampus -- Insular and Frontal Opercular Cortex | Right Subcortical Hippocampus -- Right Posterior Insular Area 2 | 0.002621845 |
| Intra (Left) | Auditory Association Cortex -- Auditory Association Cortex | Left Area STSd posterior -- Left Area STSv anterior | 0.002634723 |
| Intra (Right) | Anterior Cingulate and Medial Prefrontal Cortex -- Lateral Temporal Cortex | Right Area 8BM -- Right Area TE1 posterior | 0.002676986 |
| Inter | Ventral Stream -- Dorsal Stream | Right VentroMedial Visual Area 1 -- Left Sixth Visual Area | 0.002691194 |
| Inter | MT+ Complex and Neighbors -- MT+ Complex and Neighbors | Left Area Lateral Occipital 2 -- Right Area PH | 0.002695466 |
| Inter | MT+ Complex and Neighbors -- Dorsal Stream | Left Medial Superior Temporal Area -- Right Seventh Visual Area | 0.002696346 |
| Intra (Right) | Superior Parietal and IPS Cortex -- Ventral Stream | Right Area Lateral IntraParietal ventral -- Right VentroMedial Visual Area 1 | 0.002712954 |
| Intra (Right) | Posterior Cingulate Cortex -- Posterior Cingulate Cortex | Right Parieto-Occipital Sulcus Area 2 -- Right Area ventral 23 a+b | 0.002728195 |
| Intra (Left) | Posterior Opercular Cortex -- MT+ Complex and Neighbors | Left Area OP1/SII -- Left Area V4t | 0.002748391 |
| Intra (Right) | MT+ Complex and Neighbors -- Dorsal Stream | Right Area FST -- Right Seventh Visual Area | 0.002849218 |
| Intra (Right) | Inferior Parietal Cortex -- Lateral Temporal Cortex | Right Area PFm Complex -- Right Area TE1 posterior | 0.00285679 |
| Intra (Right) | Sensorimotor Associated Paracentral Lobular and Mid Cingulate Cortex -- Somatosensory and Motor Cortex | Right Area 5m -- Right Area 2 | 0.002927145 |
| Intra (Right) | Lateral Temporal Cortex -- Medial Temporal Cortex | Right Area TF -- Right PreSubiculum | 0.002942681 |
| Inter | Sensorimotor Associated Paracentral Lobular and Mid Cingulate Cortex -- Dorsal Stream | Left Area 5m -- Right Sixth Visual Area | 0.003077403 |
| Inter | Superior Parietal and IPS Cortex -- Ventral Stream | Right Area Lateral IntraParietal dorsal -- Left VentroMedial Visual Area 1 | 0.003124033 |
| Inter | Temporal-Parietal-Occipital Junction -- MT+ Complex and Neighbors | Left Area TemporoParietoOccipital Junction 1 -- Right Area FST | 0.003128652 |
| Intra (Right) | Caudate -- Inferior Parietal Cortex | Right Caudate -- Right Area PFm Complex | 0.003217264 |
| Intra (Left) | Ventral Stream -- Dorsal Stream | Left VentroMedial Visual Area 2 -- Left Sixth Visual Area | 0.003284831 |
| Intra (Left) | Medial Temporal Cortex -- Early Visual Cortex | Left ParaHippocampal Area 3 -- Left Third Visual Area | 0.003296554 |
| Intra (Left) | Posterior Cingulate Cortex -- Temporal-Parietal-Occipital Junction | Left PreCuneus Visual Area -- Left Superior Temporal Visual Area | 0.003309116 |
| Inter | Hippocampus -- Sensorimotor Associated Paracentral Lobular and Mid Cingulate Cortex | Right Subcortical Hippocampus -- Left Area 5m | 0.003369655 |
| Intra (Left) | Inferior Frontal Cortex -- Posterior Cingulate Cortex | Left Area 45 -- Left Area 31p ventral | 0.00343202 |
| Inter | Posterior Cingulate Cortex -- Medial Temporal Cortex | Right Area dorsal 23 a+b -- Left ParaHippocampal Area 2 | 0.003440384 |
| Inter | Sensorimotor Associated Paracentral Lobular and Mid Cingulate Cortex -- Somatosensory and Motor Cortex | Right Area 5m -- Left Primary Sensory Cortex | 0.003513276 |
| Inter | Superior Parietal and IPS Cortex -- Medial Temporal Cortex | Right Area Lateral IntraParietal dorsal -- Left ParaHippocampal Area 2 | 0.003529481 |
| Intra (Right) | Inferior Frontal Cortex -- Anterior Cingulate and Medial Prefrontal Cortex | Right Area 47l (47 lateral) -- Right Area 9 Middle | 0.00353246 |
| Intra (Right) | Posterior Cingulate Cortex -- Early Auditory Cortex | Right Area ventral 23 a+b -- Right Lateral Belt Complex | 0.003543534 |
| Intra (Right) | Orbital and Polar Frontal Cortex -- Lateral Temporal Cortex | Right Area 47m -- Right Area PHT | 0.003576824 |
| Inter | MT+ Complex and Neighbors -- Dorsal Stream | Right Area FST -- Left Seventh Visual Area | 0.003585885 |
| Inter | Ventral Stream -- Early Visual Cortex | Right VentroMedial Visual Area 1 -- Left Second Visual Area | 0.003588681 |
| Inter | Insular and Frontal Opercular Cortex -- Insular and Frontal Opercular Cortex | Left Area Posterior Insular 1 -- Right Area Posterior Insular 1 | 0.003666512 |
| Intra (Left) | Inferior Frontal Cortex -- Inferior Parietal Cortex | Left Area 45 -- Left Area PGi | 0.003737115 |
| Intra (Right) | Posterior Cingulate Cortex -- Lateral Temporal Cortex | Right Area 7m -- Right Area TF | 0.003804432 |
| Intra (Right) | MT+ Complex and Neighbors -- Ventral Stream | Right Area FST -- Right VentroMedial Visual Area 2 | 0.003856338 |
| Intra (Right) | Inferior Parietal Cortex -- Ventral Stream | Right Area IntraParietal 0 -- Right VentroMedial Visual Area 2 | 0.003889477 |
| Intra (Right) | Ventral Stream -- Dorsal Stream | Right VentroMedial Visual Area 1 -- Right Area V3B | 0.00399932 |
| Inter | Temporal-Parietal-Occipital Junction -- Lateral Temporal Cortex | Left Area TemporoParietoOccipital Junction 1 -- Right Area TF | 0.004051422 |
| Intra (Right) | Inferior Frontal Cortex -- Premotor Cortex | Right Area IFSp -- Right Premotor Eye Field | 0.004070978 |
| Intra (Right) | Posterior Cingulate Cortex -- Inferior Parietal Cortex | Right PreCuneus Visual Area -- Right Area IntraParietal 0 | 0.004100849 |
| Intra (Left) | Posterior Cingulate Cortex -- Posterior Cingulate Cortex | Left Parieto-Occipital Sulcus Area 2 -- Left Area ventral 23 a+b | 0.004176303 |
| Intra (Right) | Caudate -- Ventral Stream | Right Caudate -- Right VentroMedial Visual Area 3 | 0.004280185 |
| Intra (Left) | Posterior Cingulate Cortex -- Inferior Parietal Cortex | Left Area dorsal 23 a+b -- Left Area IntraParietal 1 | 0.004484827 |
| Inter | Posterior Cingulate Cortex -- Early Visual Cortex | Right PreCuneus Visual Area -- Left Third Visual Area | 0.004634047 |
| Inter | MT+ Complex and Neighbors -- Dorsal Stream | Right Area V3CD -- Left Seventh Visual Area | 0.004719914 |
| Inter | Inferior Frontal Cortex -- Posterior Cingulate Cortex | Left Area 45 -- Right Area 31pd | 0.004760555 |
| Intra (Right) | Dorsolateral Prefrontal Cortex -- Posterior Cingulate Cortex | Right Area 46 -- Right Area 23c | 0.00481983 |
| Inter | Orbital and Polar Frontal Cortex -- Lateral Temporal Cortex | Left Area 47m -- Right Area PHT | 0.004869026 |
| Inter | Ventral Stream -- Dorsal Stream | Left Ventral Visual Complex -- Right Area V3B | 0.00488383 |
| Intra (Left) | Posterior Cingulate Cortex -- Insular and Frontal Opercular Cortex | Left Area 31p ventral -- Left Anterior Agranular Insula Complex | 0.004894672 |
| Intra (Right) | Medial Temporal Cortex -- Sensorimotor Associated Paracentral Lobular and Mid Cingulate Cortex | Right Entorhinal Cortex -- Right Area 5m | 0.005046491 |
| Inter | Ventral Stream -- Dorsal Stream | Left Ventral Visual Complex -- Right IntraParietal Sulcus Area 1 | 0.005051786 |
| Inter | Ventral Diencephalon -- Dorsolateral Prefrontal Cortex | Left VentralDiencephalon -- Right Area 9-46d | 0.005078513 |
| Intra (Left) | Ventral Stream -- Dorsal Stream | Left Eighth Visual Area -- Left IntraParietal Sulcus Area 1 | 0.005101079 |
| Intra (Left) | Posterior Cingulate Cortex -- Superior Parietal and IPS Cortex | Left PreCuneus Visual Area -- Left Area Lateral IntraParietal dorsal | 0.00530802 |
| Intra (Right) | Dorsolateral Prefrontal Cortex -- Superior Parietal and IPS Cortex | Right Area anterior 9-46v -- Right Lateral Area 7P | 0.005399405 |
| Intra (Right) | Posterior Cingulate Cortex -- Inferior Parietal Cortex | Right Area ventral 23 a+b -- Right Area PGs | 0.005422348 |
| Intra (Right) | Posterior Cingulate Cortex -- Lateral Temporal Cortex | Right Area ventral 23 a+b -- Right Area TF | 0.005480283 |
| Inter | Posterior Cingulate Cortex -- Inferior Parietal Cortex | Right PreCuneus Visual Area -- Left Area IntraParietal 0 | 0.005480496 |
| Inter | Hippocampus -- Dorsal Stream | Left Subcortical Hippocampus -- Right Area V6A | 0.00549733 |
| Intra (Left) | Medial Temporal Cortex -- Dorsal Stream | Left ParaHippocampal Area 3 -- Left Sixth Visual Area | 0.005623189 |
| Intra (Right) | Auditory Association Cortex -- MT+ Complex and Neighbors | Right Area STGa -- Right Area FST | 0.005686629 |
| Inter | Thalamus -- Medial Temporal Cortex | Right Thalamus -- Left ParaHippocampal Area 2 | 0.005689849 |
| Inter | MT+ Complex and Neighbors -- MT+ Complex and Neighbors | Left Area V3CD -- Right Area PH | 0.005734124 |
| Inter | Medial Temporal Cortex -- Sensorimotor Associated Paracentral Lobular and Mid Cingulate Cortex | Left ParaHippocampal Area 1 -- Right Area 5m ventral | 0.005771912 |
| Intra (Right) | Ventral Stream -- Primary Visual Cortex (V1) | Right VentroMedial Visual Area 2 -- Right Primary Visual Cortex | 0.005809395 |
| Intra (Right) | Posterior Cingulate Cortex -- Insular and Frontal Opercular Cortex | Right Area ventral 23 a+b -- Right Area Frontal Opercular 5 | 0.005917214 |
| Inter | Ventral Stream -- Dorsal Stream | Right VentroMedial Visual Area 1 -- Left IntraParietal Sulcus Area 1 | 0.005936656 |
| Intra (Left) | Hippocampus -- MT+ Complex and Neighbors | Left Subcortical Hippocampus -- Left Area Lateral Occipital 2 | 0.005967365 |
| Intra (Right) | Inferior Parietal Cortex -- Dorsal Stream | Right Area IntraParietal 0 -- Right Area V3B | 0.005971 |
| Intra (Left) | Thalamus -- Posterior Cingulate Cortex | Left Thalamus -- Left Area ventral 23 a+b | 0.006107972 |
| Intra (Left) | Posterior Cingulate Cortex -- Medial Temporal Cortex | Left Dorsal Transitional Visual Area -- Left ParaHippocampal Area 2 | 0.006168977 |
| Inter | MT+ Complex and Neighbors -- Dorsal Stream | Left Medial Superior Temporal Area -- Right Area V3A | 0.006215525 |
| Intra (Left) | Posterior Cingulate Cortex -- Inferior Parietal Cortex | Left Area ventral 23 a+b -- Left Area IntraParietal 2 | 0.006234715 |
| Intra (Right) | Dorsolateral Prefrontal Cortex -- Inferior Frontal Cortex | Right Area 8B Lateral -- Right Area 44 | 0.006614751 |
| Inter | MT+ Complex and Neighbors -- Early Visual Cortex | Left Area PH -- Right Fourth Visual Area | 0.00661844 |
| Intra (Right) | Posterior Cingulate Cortex -- Inferior Parietal Cortex | Right Area ventral 23 a+b -- Right Area IntraParietal 0 | 0.00682011 |
| Inter | Posterior Cingulate Cortex -- Inferior Parietal Cortex | Left Area ventral 23 a+b -- Right Area IntraParietal 1 | 0.006846181 |
| Intra (Right) | Posterior Cingulate Cortex -- Lateral Temporal Cortex | Right PreCuneus Visual Area -- Right Area TF | 0.006884868 |
| Inter | MT+ Complex and Neighbors -- Dorsal Stream | Right Area FST -- Left IntraParietal Sulcus Area 1 | 0.006900065 |
| Inter | Posterior Cingulate Cortex -- Inferior Parietal Cortex | Right Area ventral 23 a+b -- Left Area IntraParietal 2 | 0.006930823 |
| Intra (Left) | Hippocampus -- Posterior Cingulate Cortex | Left Subcortical Hippocampus -- Left Dorsal Transitional Visual Area | 0.00695599 |
| Intra (Left) | MT+ Complex and Neighbors -- MT+ Complex and Neighbors | Left Area FST -- Left Area V3CD | 0.007008602 |
| Intra (Left) | Auditory Association Cortex -- Sensorimotor Associated Paracentral Lobular and Mid Cingulate Cortex | Left Auditory 4 Complex -- Left Area 5m | 0.007104774 |
| Intra (Right) | Medial Temporal Cortex -- Ventral Stream | Right PreSubiculum -- Right Eighth Visual Area | 0.007128196 |
| Intra (Right) | Ventral Stream -- Dorsal Stream | Right VentroMedial Visual Area 2 -- Right Area V3B | 0.007164344 |
| Inter | MT+ Complex and Neighbors -- Ventral Stream | Right Area Lateral Occipital 3 -- Left VentroMedial Visual Area 2 | 0.007382646 |
| Intra (Right) | Inferior Parietal Cortex -- Lateral Temporal Cortex | Right Area IntraParietal 2 -- Right Area TE1 posterior | 0.007439326 |
| Inter | Ventral Stream -- Early Visual Cortex | Right VentroMedial Visual Area 2 -- Left Third Visual Area | 0.007521318 |
| Intra (Left) | Superior Parietal and IPS Cortex -- Medial Temporal Cortex | Left Area Lateral IntraParietal dorsal -- Left ParaHippocampal Area 3 | 0.007796366 |
| Intra (Right) | Thalamus -- Orbital and Polar Frontal Cortex | Right Thalamus -- Right Area anterior 47r | 0.00791286 |
| Intra (Left) | Posterior Cingulate Cortex -- MT+ Complex and Neighbors | Left PreCuneus Visual Area -- Left Area PH | 0.007993006 |
| Intra (Left) | Posterior Cingulate Cortex -- Early Auditory Cortex | Left Dorsal Transitional Visual Area -- Left RetroInsular Cortex | 0.00802979 |
| Intra (Left) | Sensorimotor Associated Paracentral Lobular and Mid Cingulate Cortex -- Somatosensory and Motor Cortex | Left Area 5m -- Left Area 1 | 0.008045605 |
| Inter | MT+ Complex and Neighbors -- Dorsal Stream | Right Area FST -- Left Area V3A | 0.008136207 |
| Inter | Posterior Cingulate Cortex -- Lateral Temporal Cortex | Left Area 31pd -- Right Area TF | 0.008269791 |
| Inter | Posterior Opercular Cortex -- Sensorimotor Associated Paracentral Lobular and Mid Cingulate Cortex | Right Area PFcm -- Left Area 5m | 0.00833285 |
| Inter | Posterior Cingulate Cortex -- Medial Temporal Cortex | Left Dorsal Transitional Visual Area -- Right PreSubiculum | 0.008402389 |
| Intra (Right) | MT+ Complex and Neighbors -- Dorsal Stream | Right Area Lateral Occipital 1 -- Right IntraParietal Sulcus Area 1 | 0.008829377 |
| Inter | MT+ Complex and Neighbors -- Ventral Stream | Right Area Lateral Occipital 1 -- Left Ventral Visual Complex | 0.008835251 |
| Intra (Left) | Sensorimotor Associated Paracentral Lobular and Mid Cingulate Cortex -- Somatosensory and Motor Cortex | Left Area 5m -- Left Area 2 | 0.009025428 |
| Inter | Posterior Cingulate Cortex -- Posterior Cingulate Cortex | Left Area 23d -- Right Area 31pd | 0.00918744 |
| Inter | MT+ Complex and Neighbors -- Dorsal Stream | Left Area Lateral Occipital 1 -- Right Seventh Visual Area | 0.009278661 |
| Inter | MT+ Complex and Neighbors -- Ventral Stream | Right Middle Temporal Area -- Left VentroMedial Visual Area 2 | 0.009332221 |
| Inter | Posterior Cingulate Cortex -- Inferior Parietal Cortex | Right Area ventral 23 a+b -- Left Area IntraParietal 0 | 0.00938037 |
| Intra (Left) | Posterior Cingulate Cortex -- Insular and Frontal Opercular Cortex | Left Dorsal Transitional Visual Area -- Left Insular Granular Complex | 0.009505686 |

| | | | |
|---|---|---|---|
| Inter | Insular and Frontal Opercular Cortex -- Sensorimotor Associated Paracentral Lobular and Mid Cingulate Cortex | Right Insular Granular Complex -- Left Area 5m | 0.009594184 |
| Intra (Right) | Lateral Temporal Cortex -- Ventral Stream | Right Area PHT -- Right VentroMedial Visual Area 2 | 0.009612815 |
| Intra (Right) | Superior Parietal and IPS Cortex -- Primary Visual Cortex (V1) | Right Area Lateral IntraParietal dorsal -- Right Primary Visual Cortex | 0.009636502 |
| Intra (Right) | Posterior Cingulate Cortex -- Early Visual Cortex | Right PreCuneus Visual Area -- Right Fourth Visual Area | 0.009742514 |
| Intra (Right) | Lateral Temporal Cortex -- Insular and Frontal Opercular Cortex | Right Area PHT -- Right Frontal OPercular Area 2 | 0.009750739 |
| Inter | Anterior Cingulate and Medial Prefrontal Cortex -- Posterior Cingulate Cortex | Left Area posterior 24 -- Right Area 31p ventral | 0.009816577 |
| Inter | Posterior Cingulate Cortex -- Primary Visual Cortex (V1) | Right Area 31pd -- Left Primary Visual Cortex | 0.009839548 |
| Intra (Right) | Caudate -- Posterior Cingulate Cortex | Right Caudate -- Right Dorsal Transitional Visual Area | 0.010234309 |
| Intra (Left) | Inferior Parietal Cortex -- Insular and Frontal Opercular Cortex | Left Area IntraParietal 0 -- Left Area Frontal Opercular 5 | 0.010234577 |
| Inter | Ventral Stream -- Dorsal Stream | Right Eighth Visual Area -- Left Area V3A | 0.01062931 |
| Inter | Sensorimotor Associated Paracentral Lobular and Mid Cingulate Cortex -- Somatosensory and Motor Cortex | Left Area 5m -- Right Area 3a | 0.010763268 |
| Intra (Left) | MT+ Complex and Neighbors -- Dorsal Stream | Left Area V4t -- Left IntraParietal Sulcus Area 1 | 0.010791935 |
| Inter | Temporal-Parietal-Occipital Junction -- Dorsal Stream | Left Area TemporoParietoOccipital Junction 1 -- Right Seventh Visual Area | 0.010871668 |
| Inter | Inferior Parietal Cortex -- Ventral Stream | Left Area IntraParietal 0 -- Right VentroMedial Visual Area 2 | 0.010879983 |
| Intra (Right) | Caudate -- Lateral Temporal Cortex | Right Caudate -- Right Area TE1 posterior | 0.011008277 |
| Intra (Left) | MT+ Complex and Neighbors -- Dorsal Stream | Left Area FST -- Left Area V6A | 0.011021981 |
| Intra (Left) | Posterior Cingulate Cortex -- MT+ Complex and Neighbors | Left PreCuneus Visual Area -- Left Area FST | 0.011051348 |
| Inter | Posterior Cingulate Cortex -- Lateral Temporal Cortex | Left Dorsal Transitional Visual Area -- Right Area TE2 posterior | 0.011055595 |
| Inter | Posterior Cingulate Cortex -- Inferior Parietal Cortex | Right Area 31p ventral -- Left Area IntraParietal 0 | 0.011114132 |
| Inter | Dorsolateral Prefrontal Cortex -- Medial Temporal Cortex | Right Area 46 -- Left ParaHippocampal Area 3 | 0.011187546 |
| Inter | Hippocampus -- Sensorimotor Associated Paracentral Lobular and Mid Cingulate Cortex | Right Subcortical Hippocampus -- Left Area 5L | 0.011190725 |
| Intra (Left) | Hippocampus -- Sensorimotor Associated Paracentral Lobular and Mid Cingulate Cortex | Left Subcortical Hippocampus -- Left Area 5m ventral | 0.011341982 |
| Intra (Right) | Posterior Cingulate Cortex -- MT+ Complex and Neighbors | Right PreCuneus Visual Area -- Right Area V3CD | 0.011437673 |
| Intra (Left) | Superior Parietal and IPS Cortex -- Superior Parietal and IPS Cortex | Left Lateral Area 7P -- Left Area 7PC | 0.011466738 |
| Inter | Hippocampus -- Somatosensory and Motor Cortex | Left Subcortical Hippocampus -- Right Area 2 | 0.011707036 |
| Inter | MT+ Complex and Neighbors -- Ventral Stream | Right Area PH -- Left Eighth Visual Area | 0.011720459 |
| Inter | Ventral Stream -- Ventral Stream | Left Ventral Visual Complex -- Right Eighth Visual Area | 0.011782789 |
| Inter | Temporal-Parietal-Occipital Junction -- Ventral Stream | Right Area TemporoParietoOccipital Junction 3 -- Left VentroMedial Visual Area 2 | 0.011851848 |
| Inter | Sensorimotor Associated Paracentral Lobular and Mid Cingulate Cortex -- Somatosensory and Motor Cortex | Right Area 5m -- Left Area 2 | 0.011858086 |
| Intra (Left) | Sensorimotor Associated Paracentral Lobular and Mid Cingulate Cortex -- Somatosensory and Motor Cortex | Left Area 5m -- Left Area 3a | 0.012018524 |
| Intra (Left) | Ventral Stream -- Early Visual Cortex | Left VentroMedial Visual Area 2 -- Left Second Visual Area | 0.01218065 |
| Intra (Right) | Caudate -- Lateral Temporal Cortex | Right Caudate -- Right Area TE2 anterior | 0.012323737 |
| Intra (Right) | Dorsal Stream -- Dorsal Stream | Right Seventh Visual Area -- Right Area V3B | 0.012492622 |
| Intra (Left) | Dorsal Stream -- Dorsal Stream | Left Area V3A -- Left Area V3B | 0.012547113 |
| Intra (Right) | Posterior Cingulate Cortex -- Superior Parietal and IPS Cortex | Right Area dorsal 23 a+b -- Right Medial IntraParietal Area | 0.012557847 |
| Intra (Right) | Posterior Cingulate Cortex -- Insular and Frontal Opercular Cortex | Right Area ventral 23 a+b -- Right Area Posterior Insular 1 | 0.012631361 |
| Inter | Superior Parietal and IPS Cortex -- Primary Visual Cortex (V1) | Right Area Lateral IntraParietal dorsal -- Left Primary Visual Cortex | 0.012634381 |
| Intra (Left) | Sensorimotor Associated Paracentral Lobular and Mid Cingulate Cortex -- Sensorimotor Associated Paracentral Lobular and Mid Cingulate Cortex | Left Area 5m -- Left Dorsal Area 24d | 0.012807734 |
| Inter | Sensorimotor Associated Paracentral Lobular and Mid Cingulate Cortex -- Ventral Stream | Left Area 5m -- Right VentroMedial Visual Area 1 | 0.012861957 |
| Inter | MT+ Complex and Neighbors -- Dorsal Stream | Right Area Lateral Occipital 1 -- Left Seventh Visual Area | 0.012987591 |
| Intra (Right) | MT+ Complex and Neighbors -- Ventral Stream | Right Area PH -- Right VentroMedial Visual Area 1 | 0.013099153 |
| Intra (Right) | Medial Temporal Cortex -- Dorsal Stream | Right ParaHippocampal Area 3 -- Right IntraParietal Sulcus Area 1 | 0.013288708 |
| Intra (Right) | Inferior Parietal Cortex -- MT+ Complex and Neighbors | Right Area IntraParietal 0 -- Right Area PH | 0.013304972 |
| Inter | Caudate -- Inferior Frontal Cortex | Left Caudate -- Right Area 44 | 0.013405823 |
| Inter | Posterior Cingulate Cortex -- Superior Parietal and IPS Cortex | Right Area ventral 23 a+b -- Left Anterior IntraParietal Area | 0.013476793 |
| Inter | Posterior Cingulate Cortex -- Ventral Stream | Right PreCuneus Visual Area -- Left Fusiform Face Complex | 0.013490496 |
| Intra (Left) | MT+ Complex and Neighbors -- Dorsal Stream | Left Area FST -- Left IntraParietal Sulcus Area 1 | 0.013570379 |
| Inter | Dorsolateral Prefrontal Cortex -- Posterior Cingulate Cortex | Left Area 8Av -- Right Area 7m | 0.013685696 |
| Intra (Right) | Sensorimotor Associated Paracentral Lobular and Mid Cingulate Cortex -- Somatosensory and Motor Cortex | Right Area 5m -- Right Area 1 | 0.01369284 |
| Intra (Right) | Caudate -- Superior Parietal and IPS Cortex | Right Caudate -- Right Area Lateral IntraParietal ventral | 0.013758424 |
| Intra (Right) | Hippocampus -- Sensorimotor Associated Paracentral Lobular and Mid Cingulate Cortex | Right Subcortical Hippocampus -- Right Area 5m ventral | 0.013869643 |
| Inter | MT+ Complex and Neighbors -- Dorsal Stream | Left Area PH -- Right Area V3A | 0.013921266 |
| Inter | Orbital and Polar Frontal Cortex -- Posterior Cingulate Cortex | Right Area 47s -- Left Area 31pd | 0.014174601 |
| Inter | Posterior Cingulate Cortex -- Lateral Temporal Cortex | Left PreCuneus Visual Area -- Right Area TF | 0.014189661 |
| Intra (Right) | Hippocampus -- MT+ Complex and Neighbors | Right Subcortical Hippocampus -- Right Area FST | 0.014425403 |
| Intra (Right) | Superior Parietal and IPS Cortex -- Lateral Temporal Cortex | Right Lateral Area 7P -- Right Area TE2 posterior | 0.014551068 |
| Inter | Orbital and Polar Frontal Cortex -- Superior Parietal and IPS Cortex | Right Area 47m -- Left Area Lateral IntraParietal dorsal | 0.014619016 |
| Intra (Right) | Hippocampus -- Somatosensory and Motor Cortex | Right Subcortical Hippocampus -- Right Area 2 | 0.015071406 |
| Inter | Posterior Cingulate Cortex -- Lateral Temporal Cortex | Right PreCuneus Visual Area -- Left Area TE2 posterior | 0.015105901 |
| Inter | Temporal-Parietal-Occipital Junction -- Ventral Stream | Left Area TemporoParietoOccipital Junction 2 -- Right VentroMedial Visual Area 3 | 0.015238768 |
| Inter | Posterior Cingulate Cortex -- Early Visual Cortex | Right PreCuneus Visual Area -- Left Fourth Visual Area | 0.015242641 |
| Inter | Ventral Stream -- Dorsal Stream | Left VentroMedial Visual Area 2 -- Right Seventh Visual Area | 0.015253175 |
| Intra (Right) | Superior Parietal and IPS Cortex -- Early Auditory Cortex | Right Area Lateral IntraParietal dorsal -- Right Lateral Belt Complex | 0.015318357 |
| Intra (Right) | Cerebellum -- Medial Temporal Cortex | Right Cerebellum -- Right ParaHippocampal Area 1 | 0.015366398 |
| Intra (Right) | Ventral Stream -- Early Visual Cortex | Right VentroMedial Visual Area 1 -- Right Second Visual Area | 0.01576334 |
| Intra (Left) | Temporal-Parietal-Occipital Junction -- Medial Temporal Cortex | Left Area TemporoParietoOccipital Junction 3 -- Left Perirhinal Ectorhinal Cortex | 0.015802863 |
| Inter | Lateral Temporal Cortex -- Lateral Temporal Cortex | Left Area TE2 anterior -- Right Area TE1 posterior | 0.015888472 |
| Inter | MT+ Complex and Neighbors -- MT+ Complex and Neighbors | Left Medial Superior Temporal Area -- Right Area FST | 0.01600123 |
| Inter | Posterior Cingulate Cortex -- MT+ Complex and Neighbors | Right PreCuneus Visual Area -- Left Area PH | 0.01615242 |
| Inter | Posterior Cingulate Cortex -- Primary Visual Cortex (V1) | Right Area ventral 23 a+b -- Left Primary Visual Cortex | 0.016300971 |
| Inter | Sensorimotor Associated Paracentral Lobular and Mid Cingulate Cortex -- Dorsal Stream | Right Area 5m -- Left Area V3A | 0.016425577 |
| Inter | Thalamus -- Posterior Cingulate Cortex | Left Thalamus -- Right Area ventral 23 a+b | 0.016439831 |
| Intra (Right) | Posterior Cingulate Cortex -- Superior Parietal and IPS Cortex | Right ProStriate Area -- Right Area Lateral IntraParietal dorsal | 0.016460719 |
| Inter | Hippocampus -- Superior Parietal and IPS Cortex | Left Subcortical Hippocampus -- Right Area 7PC | 0.016650916 |
| Intra (Left) | Orbital and Polar Frontal Cortex -- Posterior Cingulate Cortex | Left Polar 10p -- Left Parieto-Occipital Sulcus Area 1 | 0.016856237 |
| Intra (Right) | Posterior Cingulate Cortex -- Inferior Parietal Cortex | Right PreCuneus Visual Area -- Right Area PGp | 0.016935431 |
| Intra (Right) | Posterior Cingulate Cortex -- Lateral Temporal Cortex | Right Area 31pd -- Right Area TF | 0.017104464 |
| Inter | Posterior Cingulate Cortex -- Inferior Parietal Cortex | Right Area ventral 23 a+b -- Left Area IntraParietal 1 | 0.017261735 |
| Inter | MT+ Complex and Neighbors -- Ventral Stream | Left Area FST -- Right VentroMedial Visual Area 2 | 0.017393208 |
| Intra (Left) | Sensorimotor Associated Paracentral Lobular and Mid Cingulate Cortex -- Dorsal Stream | Left Area 5m -- Left Area V3A | 0.017515949 |
| Inter | Inferior Parietal Cortex -- Superior Parietal and IPS Cortex | Right Area PFt -- Left Anterior IntraParietal Area | 0.017911976 |
| Intra (Left) | Sensorimotor Associated Paracentral Lobular and Mid Cingulate Cortex -- Ventral Stream | Left Area 5m -- Left VentroMedial Visual Area 2 | 0.017930368 |
| Intra (Right) | Superior Parietal and IPS Cortex -- MT+ Complex and Neighbors | Right Area Lateral IntraParietal ventral -- Right Area Lateral Occipital 1 | 0.018121923 |
| Inter | Anterior Cingulate and Medial Prefrontal Cortex -- Posterior Cingulate Cortex | Left Area p32 -- Right Area ventral 23 a+b | 0.018155255 |
| Intra (Right) | Inferior Parietal Cortex -- Insular and Frontal Opercular Cortex | Right Area PF opercular -- Right Middle Insular Area | 0.018185122 |
| Intra (Right) | Ventral Diencephalon -- MT+ Complex and Neighbors | Right VentralDiencephalon -- Right Area FST | 0.018267798 |
| Intra (Right) | Lateral Temporal Cortex -- Medial Temporal Cortex | Right Area TE2 anterior -- Right PreSubiculum | 0.01830006 |
| Inter | Ventral Stream -- Dorsal Stream | Right VentroMedial Visual Area 3 -- Left IntraParietal Sulcus Area 1 | 0.018338242 |
| Inter | Lateral Temporal Cortex -- MT+ Complex and Neighbors | Right Area TF -- Left Area V4t | 0.018344069 |
| Intra (Left) | Posterior Cingulate Cortex -- Lateral Temporal Cortex | Left Parieto-Occipital Sulcus Area 1 -- Left Area TE1 posterior | 0.018419727 |
| Inter | Inferior Frontal Cortex -- Posterior Cingulate Cortex | Right Area IFJa -- Left PreCuneus Visual Area | 0.018539808 |
| Inter | Dorsolateral Prefrontal Cortex -- Inferior Parietal Cortex | Right Area 8B Lateral -- Left Area IntraParietal 1 | 0.018550846 |
| Inter | Ventral Stream -- Dorsal Stream | Left VentroMedial Visual Area 2 -- Right Sixth Visual Area | 0.018591147 |
| Inter | Anterior Cingulate and Medial Prefrontal Cortex -- Anterior Cingulate and Medial Prefrontal Cortex | Left Area 10v -- Right Area 10v | 0.018894418 |
| Inter | Inferior Frontal Cortex -- Medial Temporal Cortex | Right Area IFJa -- Left ParaHippocampal Area 1 | 0.019091362 |
| Intra (Left) | Medial Temporal Cortex -- MT+ Complex and Neighbors | Left ParaHippocampal Area 3 -- Left Area V3CD | 0.019155537 |
| Intra (Right) | Temporal-Parietal-Occipital Junction -- MT+ Complex and Neighbors | Right Area TemporoParietoOccipital Junction 2 -- Right Area V4t | 0.019419631 |
| Inter | Insular and Frontal Opercular Cortex -- Sensorimotor Associated Paracentral Lobular and Mid Cingulate Cortex | Right Area Posterior Insular 1 -- Left Area 5m ventral | 0.019500933 |
| Intra (Right) | Hippocampus -- Superior Parietal and IPS Cortex | Right Subcortical Hippocampus -- Right Area 7PC | 0.019513614 |
| Inter | Superior Parietal and IPS Cortex -- Dorsal Stream | Right Area Lateral IntraParietal dorsal -- Left Sixth Visual Area | 0.019560224 |
| Intra (Right) | MT+ Complex and Neighbors -- MT+ Complex and Neighbors | Right Area Lateral Occipital 1 -- Right Area PH | 0.019665588 |
| Inter | Cerebellum -- Dorsolateral Prefrontal Cortex | Right Cerebellum -- Left Area 8Ad | 0.019839296 |
| Intra (Right) | Posterior Cingulate Cortex -- Early Visual Cortex | Right PreCuneus Visual Area -- Right Third Visual Area | 0.019886306 |
| Inter | Superior Parietal and IPS Cortex -- Ventral Stream | Right Anterior IntraParietal Area -- Left VentroMedial Visual Area 2 | 0.020033974 |
| Intra (Left) | MT+ Complex and Neighbors -- Dorsal Stream | Left Medial Superior Temporal Area -- Left Area V3A | 0.020279319 |
| Inter | Posterior Cingulate Cortex -- Inferior Parietal Cortex | Right PreCuneus Visual Area -- Left Area IntraParietal 1 | 0.020297769 |
| Intra (Right) | Posterior Cingulate Cortex -- Inferior Parietal Cortex | Right Parieto-Occipital Sulcus Area 1 -- Right Area IntraParietal 1 | 0.020530924 |
| Intra (Right) | Anterior Cingulate and Medial Prefrontal Cortex -- Inferior Parietal Cortex | Right Area s32 -- Right Area IntraParietal 1 | 0.020542969 |
| Inter | Dorsolateral Prefrontal Cortex -- Auditory Association Cortex | Right Area 9 anterior -- Left Area STSv posterior | 0.02066324 |
| Intra (Left) | Inferior Frontal Cortex -- Posterior Cingulate Cortex | Left Area 45 -- Left Area 31pd | 0.020690735 |
| Intra (Left) | Superior Parietal and IPS Cortex -- Lateral Temporal Cortex | Left Medial IntraParietal Area -- Left Area TE2 posterior | 0.021594912 |
| Intra (Left) | Sensorimotor Associated Paracentral Lobular and Mid Cingulate Cortex -- Somatosensory and Motor Cortex | Left Area 5m -- Left Primary Motor Cortex | 0.021661625 |
| Intra (Right) | Posterior Cingulate Cortex -- Medial Temporal Cortex | Right Area 7m -- Right Entorhinal Cortex | 0.021713674 |
| Inter | MT+ Complex and Neighbors -- Dorsal Stream | Left Area FST -- Right IntraParietal Sulcus Area 1 | 0.021875108 |
| Intra (Right) | Caudate -- Inferior Parietal Cortex | Right Caudate -- Right Area PFt | 0.021908703 |
| Inter | Lateral Temporal Cortex -- MT+ Complex and Neighbors | Right Area TF -- Left Middle Temporal Area | 0.022060715 |
| Intra (Left) | Posterior Opercular Cortex -- MT+ Complex and Neighbors | Left Area OP1/SII -- Left Medial Superior Temporal Area | 0.022089624 |
| Inter | Sensorimotor Associated Paracentral Lobular and Mid Cingulate Cortex -- Dorsal Stream | Left Area 5m -- Right Area V3A | 0.022110143 |
| Intra (Left) | Inferior Parietal Cortex -- Inferior Parietal Cortex | Left Area IntraParietal 1 -- Left Area PF Complex | 0.02272323 |
| Intra (Left) | Inferior Frontal Cortex -- Inferior Parietal Cortex | Left Area 44 -- Left Area PGi | 0.022971246 |
| Inter | Posterior Cingulate Cortex -- Medial Temporal Cortex | Right Area 23c -- Left ParaHippocampal Area 2 | 0.023062838 |
| Intra (Right) | Superior Parietal and IPS Cortex -- Ventral Stream | Right Area Lateral IntraParietal ventral -- Right VentroMedial Visual Area 2 | 0.023082967 |
| Inter | Ventral Stream -- Ventral Stream | Left VentroMedial Visual Area 2 -- Right Ventral Visual Complex | 0.02339479 |
| Inter | Inferior Parietal Cortex -- Premotor Cortex | Left Area IntraParietal 1 -- Right Premotor Eye Field | 0.023396533 |
| Intra (Right) | Inferior Frontal Cortex -- Inferior Parietal Cortex | Right Area IFJp -- Right Area PGs | 0.023456201 |
| Inter | Posterior Cingulate Cortex -- Ventral Stream | Left PreCuneus Visual Area -- Right Ventral Visual Complex | 0.023540705 |
| Inter | Posterior Opercular Cortex -- MT+ Complex and Neighbors | Right Area OP1/SII -- Left Middle Temporal Area | 0.023742337 |
| Intra (Left) | MT+ Complex and Neighbors -- Early Visual Cortex | Left Area FST -- Left Fourth Visual Area | 0.023759247 |
| Intra (Right) | Posterior Cingulate Cortex -- Superior Parietal and IPS Cortex | Right ProStriate Area -- Right Area Lateral IntraParietal ventral | 0.024111757 |
| Intra (Left) | Posterior Cingulate Cortex -- Posterior Cingulate Cortex | Left Parieto-Occipital Sulcus Area 1 -- Left Area 31a | 0.024533477 |
| Inter | Anterior Cingulate and Medial Prefrontal Cortex -- Auditory Association Cortex | Left Area 10v -- Right Area STGa | 0.024648295 |
| Intra (Right) | Dorsolateral Prefrontal Cortex -- Orbital and Polar Frontal Cortex | Right Area posterior 9-46v -- Right Area posterior 10p | 0.024902022 |
| Intra (Right) | Superior Parietal and IPS Cortex -- Ventral Stream | Right Area Lateral IntraParietal dorsal -- Right VentroMedial Visual Area 1 | 0.024905753 |
| Intra (Left) | Thalamus -- Medial Temporal Cortex | Left Thalamus -- Left ParaHippocampal Area 2 | 0.025533432 |
| Inter | Ventral Stream -- Dorsal Stream | Left VentroMedial Visual Area 2 -- Right Area V6A | 0.025549365 |
| Inter | Posterior Cingulate Cortex -- Ventral Stream | Left PreCuneus Visual Area -- Right VentroMedial Visual Area 1 | 0.025606227 |
| Inter | Dorsal Stream -- Primary Visual Cortex (V1) | Right Area V3B -- Left Primary Visual Cortex | 0.025725839 |
| Intra (Right) | Anterior Cingulate and Medial Prefrontal Cortex -- Posterior Cingulate Cortex | Right Anterior 24 prime -- Right Area ventral 23 a+b | 0.025743545 |
| Intra (Left) | Ventral Stream -- Ventral Stream | Left Fusiform Face Complex -- Left VentroMedial Visual Area 2 | 0.025761363 |
| Intra (Right) | Superior Parietal and IPS Cortex -- Lateral Temporal Cortex | Right Medial IntraParietal Area -- Right Area TE2 posterior | 0.025846463 |
| Intra (Left) | Inferior Frontal Cortex -- Posterior Cingulate Cortex | Left Area 47l (47 lateral) -- Left Area 31pd | 0.026078489 |
| Inter | Inferior Parietal Cortex -- MT+ Complex and Neighbors | Right Area IntraParietal 0 -- Left Area PH | 0.026496165 |
| Intra (Left) | MT+ Complex and Neighbors -- Ventral Stream | Left Area Lateral Occipital 1 -- Left Ventral Visual Complex | 0.027061106 |

| | | | |
|---|---|---|---|
| Inter | Superior Parietal and IPS Cortex -- Posterior Opercular Cortex | Right Area Lateral IntraParietal ventral -- Left Area OP2-3/VS | 0.027088092 |
| Intra (Right) | Lateral Temporal Cortex -- Insular and Frontal Opercular Cortex | Right Area TE1 posterior -- Right Middle Insular Area | 0.027138571 |
| Inter | Lateral Temporal Cortex -- Premotor Cortex | Left Area TE1 posterior -- Right Premotor Eye Field | 0.027167963 |
| Intra (Right) | Superior Parietal and IPS Cortex -- Ventral Stream | Right Ventral IntraParietal Complex -- Right VentroMedial Visual Area 2 | 0.027697698 |
| Inter | Medial Temporal Cortex -- Sensorimotor Associated Paracentral Lobular and Mid Cingulate Cortex | Left ParaHippocampal Area 2 -- Right Area 5m ventral | 0.027865389 |
| Intra (Right) | Insular and Frontal Opercular Cortex -- Sensorimotor Associated Paracentral Lobular and Mid Cingulate Cortex | Right Area Posterior Insular 1 -- Right Area 5m | 0.028499953 |
| Intra (Left) | Ventral Stream -- Dorsal Stream | Left Eighth Visual Area -- Left Seventh Visual Area | 0.028535814 |
| Intra (Right) | MT+ Complex and Neighbors -- Ventral Stream | Right Area Lateral Occipital 1 -- Right VentroMedial Visual Area 2 | 0.028643393 |
| Inter | MT+ Complex and Neighbors -- Dorsal Stream | Left Medial Superior Temporal Area -- Right Area V6A | 0.028837451 |
| Inter | Dorsolateral Prefrontal Cortex -- Lateral Temporal Cortex | Left Area 8C -- Right Area TE2 anterior | 0.02885756 |
| Inter | Cerebellum -- Medial Temporal Cortex | Right Cerebellum -- Left ParaHippocampal Area 1 | 0.028904646 |
| Inter | Inferior Frontal Cortex -- Inferior Parietal Cortex | Right Area IFSa -- Left Area IntraParietal 1 | 0.028968178 |
| Intra (Left) | Temporal-Parietal-Occipital Junction -- Lateral Temporal Cortex | Left Area TemporoParietoOccipital Junction 3 -- Left Area TE2 posterior | 0.029486298 |
| Inter | MT+ Complex and Neighbors -- Dorsal Stream | Right Area PH -- Left Area V3A | 0.029504705 |
| Inter | Inferior Frontal Cortex -- Orbital and Polar Frontal Cortex | Left Area IFSp -- Right Polar 10p | 0.030146133 |
| Inter | Orbital and Polar Frontal Cortex -- Posterior Cingulate Cortex | Right Area 47m -- Left PreCuneus Visual Area | 0.030177635 |
| Inter | MT+ Complex and Neighbors -- MT+ Complex and Neighbors | Left Area V4t -- Right Area FST | 0.030379409 |
| Intra (Right) | Inferior Parietal Cortex -- Posterior Opercular Cortex | Right Area PF opercular -- Right Frontal Opercular Area 1 | 0.031132453 |
| Inter | Accumbens -- Anterior Cingulate and Medial Prefrontal Cortex | Left Accumbens -- Right Area s32 | 0.031346434 |
| Inter | Insular and Frontal Opercular Cortex -- Insular and Frontal Opercular Cortex | Left Insular Granular Complex -- Right Middle Insular Area | 0.031567127 |
| Inter | Auditory Association Cortex -- MT+ Complex and Neighbors | Right Area STGa -- Left Area V4t | 0.031570869 |
| Inter | Superior Parietal and IPS Cortex -- Ventral Stream | Right Area Lateral IntraParietal ventral -- Left Eighth Visual Area | 0.032084621 |
| Inter | Inferior Frontal Cortex -- Posterior Cingulate Cortex | Left Area 47l (47 lateral) -- Right Area 31pd | 0.032267533 |
| Intra (Left) | Sensorimotor Associated Paracentral Lobular and Mid Cingulate Cortex -- Early Visual Cortex | Left Area 5m -- Left Second Visual Area | 0.032387846 |
| Intra (Right) | Hippocampus -- Temporal-Parietal-Occipital Junction | Right Subcortical Hippocampus -- Right Area TemporoParietoOccipital Junction 2 | 0.032627625 |
| Intra (Right) | Superior Parietal and IPS Cortex -- MT+ Complex and Neighbors | Right Area Lateral IntraParietal ventral -- Right Area Lateral Occipital 3 | 0.032707492 |
| Inter | Inferior Frontal Cortex -- Posterior Cingulate Cortex | Left Area IFSp -- Right Area 31pd | 0.032769797 |
| Inter | Hippocampus -- Sensorimotor Associated Paracentral Lobular and Mid Cingulate Cortex | Right Subcortical Hippocampus -- Left Area 5m ventral | 0.032812564 |
| Intra (Right) | Dorsolateral Prefrontal Cortex -- Inferior Frontal Cortex | Right Area 8Av -- Right Area 47l (47 lateral) | 0.032867023 |
| Intra (Left) | Superior Parietal and IPS Cortex -- Medial Temporal Cortex | Left Medial Area 7A -- Left ParaHippocampal Area 2 | 0.032948766 |
| Intra (Right) | Ventral Stream -- Dorsal Stream | Right VentroMedial Visual Area 1 -- Right Sixth Visual Area | 0.033576999 |
| Intra (Left) | Medial Temporal Cortex -- Insular and Frontal Opercular Cortex | Left ParaHippocampal Area 3 -- Left Area Posterior Insular 1 | 0.033645338 |
| Inter | Sensorimotor Associated Paracentral Lobular and Mid Cingulate Cortex -- Ventral Stream | Right Area 5m ventral -- Left Ventral Visual Complex | 0.033723695 |
| Inter | Auditory Association Cortex -- MT+ Complex and Neighbors | Right Area STGa -- Left Middle Temporal Area | 0.033922611 |
| Intra (Right) | Inferior Parietal Cortex -- Sensorimotor Associated Paracentral Lobular and Mid Cingulate Cortex | Right Area PF opercular -- Right Area 5m ventral | 0.034059413 |
| Intra (Left) | Inferior Parietal Cortex -- Lateral Temporal Cortex | Left Area IntraParietal 1 -- Left Area PHT | 0.034554882 |
| Intra (Left) | Early Auditory Cortex -- Sensorimotor Associated Paracentral Lobular and Mid Cingulate Cortex | Left ParaBelt Complex -- Left Area 5m | 0.034595102 |
| Inter | Orbital and Polar Frontal Cortex -- Superior Parietal and IPS Cortex | Right Area 47m -- Left Medial Area 7A | 0.03477932 |
| Inter | Orbital and Polar Frontal Cortex -- Posterior Cingulate Cortex | Left Area posterior 10p -- Right Area ventral 23 a+b | 0.034829282 |
| Inter | Cerebellum -- Posterior Cingulate Cortex | Left Cerebellum -- Right Area 7m | 0.035186897 |
| Intra (Left) | Ventral Stream -- Ventral Stream | Left VentroMedial Visual Area 2 -- Left Ventral Visual Complex | 0.035258981 |
| Intra (Left) | MT+ Complex and Neighbors -- Dorsal Stream | Left Medial Superior Temporal Area -- Left Sixth Visual Area | 0.035532761 |
| Intra (Right) | Caudate -- Ventral Stream | Right Caudate -- Right Eighth Visual Area | 0.035775192 |
| Intra (Right) | Lateral Temporal Cortex -- Premotor Cortex | Right Area TE2 posterior -- Right Premotor Eye Field | 0.035989778 |
| Intra (Right) | Temporal-Parietal-Occipital Junction -- Ventral Stream | Right Area TemporoParietoOccipital Junction 2 -- Right Ventral Visual Complex | 0.036067919 |
| Inter | Posterior Cingulate Cortex -- Posterior Cingulate Cortex | Left Dorsal Transitional Visual Area -- Right PreCuneus Visual Area | 0.036089323 |
| Inter | Sensorimotor Associated Paracentral Lobular and Mid Cingulate Cortex -- Dorsal Stream | Right Area 5m -- Left Sixth Visual Area | 0.036322457 |
| Inter | Temporal-Parietal-Occipital Junction -- Insular and Frontal Opercular Cortex | Right Area TemporoParietoOccipital Junction 1 -- Left Insular Granular Complex | 0.036329512 |
| Inter | Posterior Cingulate Cortex -- Posterior Cingulate Cortex | Left RetroSplenial Complex -- Right Parieto-Occipital Sulcus Area 2 | 0.036362185 |
| Intra (Right) | Sensorimotor Associated Paracentral Lobular and Mid Cingulate Cortex -- MT+ Complex and Neighbors | Right Area 5m -- Right Area FST | 0.036413482 |
| Inter | Hippocampus -- MT+ Complex and Neighbors | Left Subcortical Hippocampus -- Right Area FST | 0.036422658 |
| Inter | Sensorimotor Associated Paracentral Lobular and Mid Cingulate Cortex -- Ventral Stream | Right Area 5m ventral -- Left VentroMedial Visual Area 2 | 0.036482587 |
| Inter | Superior Parietal and IPS Cortex -- Medial Temporal Cortex | Right Medial Area 7A -- Left ParaHippocampal Area 3 | 0.03652644 |
| Inter | Inferior Parietal Cortex -- Ventral Stream | Right Area IntraParietal 0 -- Left VentroMedial Visual Area 2 | 0.036880144 |
| Intra (Right) | MT+ Complex and Neighbors -- Ventral Stream | Right Middle Temporal Area -- Right VentroMedial Visual Area 2 | 0.037736439 |
| Intra (Right) | Posterior Cingulate Cortex -- Medial Temporal Cortex | Right Dorsal Transitional Visual Area -- Right PreSubiculum | 0.037743739 |
| Intra (Left) | Inferior Parietal Cortex -- Auditory Association Cortex | Left Area PGi -- Left Area STSv anterior | 0.038118922 |
| Intra (Right) | Temporal-Parietal-Occipital Junction -- Ventral Stream | Right Area TemporoParietoOccipital Junction 2 -- Right Fusiform Face Complex | 0.038180698 |
| Intra (Right) | Inferior Frontal Cortex -- Inferior Parietal Cortex | Right Area IFSa -- Right Area IntraParietal 1 | 0.038619156 |
| Inter | Orbital and Polar Frontal Cortex -- Posterior Cingulate Cortex | Left Area 13l -- Right Area ventral 23 a+b | 0.038880124 |
| Inter | Superior Parietal and IPS Cortex -- Lateral Temporal Cortex | Left Area Lateral IntraParietal dorsal -- Right Area TE2 posterior | 0.039320439 |
| Intra (Left) | MT+ Complex and Neighbors -- Dorsal Stream | Left Area Lateral Occipital 1 -- Left Area V3B | 0.039448985 |
| Inter | Temporal-Parietal-Occipital Junction -- Early Visual Cortex | Left Area TemporoParietoOccipital Junction 1 -- Right Second Visual Area | 0.039497986 |
| Inter | MT+ Complex and Neighbors -- MT+ Complex and Neighbors | Left Area FST -- Right Area Lateral Occipital 1 | 0.039522584 |
| Inter | Posterior Opercular Cortex -- Sensorimotor Associated Paracentral Lobular and Mid Cingulate Cortex | Right Area OP1/SII -- Left Area 5m | 0.039900142 |
| Intra (Right) | Temporal-Parietal-Occipital Junction -- Ventral Stream | Right Area TemporoParietoOccipital Junction 1 -- Right Ventral Visual Complex | 0.040126209 |
| Intra (Right) | MT+ Complex and Neighbors -- Ventral Stream | Right Area V3CD -- Right VentroMedial Visual Area 1 | 0.040198082 |
| Intra (Right) | Hippocampus -- MT+ Complex and Neighbors | Right Subcortical Hippocampus -- Right Middle Temporal Area | 0.040296478 |
| Intra (Left) | Temporal-Parietal-Occipital Junction -- Early Visual Cortex | Left Area TemporoParietoOccipital Junction 1 -- Left Second Visual Area | 0.040526199 |
| Intra (Left) | Inferior Parietal Cortex -- Ventral Stream | Left Area IntraParietal 0 -- Left VentroMedial Visual Area 2 | 0.040536457 |
| Inter | Anterior Cingulate and Medial Prefrontal Cortex -- Posterior Cingulate Cortex | Left Area 10r -- Right Area ventral 23 a+b | 0.040558446 |
| Intra (Right) | Temporal-Parietal-Occipital Junction -- Auditory Association Cortex | Right Area TemporoParietoOccipital Junction 2 -- Right Area STSd posterior | 0.040585055 |
| Inter | Sensorimotor Associated Paracentral Lobular and Mid Cingulate Cortex -- Somatosensory and Motor Cortex | Right Area 5m -- Left Area 3a | 0.040817463 |
| Intra (Right) | Inferior Frontal Cortex -- Temporal-Parietal-Occipital Junction | Right Area IFSa -- Right Area TemporoParietoOccipital Junction 3 | 0.041008118 |
| Inter | Hippocampus -- Dorsal Stream | Left Subcortical Hippocampus -- Right Area V3A | 0.041347394 |
| Intra (Left) | Superior Parietal and IPS Cortex -- Medial Temporal Cortex | Left Area Lateral IntraParietal dorsal -- Left ParaHippocampal Area 2 | 0.041487203 |
| Inter | Superior Parietal and IPS Cortex -- Early Visual Cortex | Right Area Lateral IntraParietal dorsal -- Left Second Visual Area | 0.041851397 |
| Intra (Left) | MT+ Complex and Neighbors -- Dorsal Stream | Left Area PH -- Left Area V3B | 0.042145555 |
| Inter | Superior Parietal and IPS Cortex -- Temporal-Parietal-Occipital Junction | Right Area Lateral IntraParietal ventral -- Left Superior Temporal Visual Area | 0.042372074 |
| Inter | Medial Temporal Cortex -- Ventral Stream | Right PreSubiculum -- Left Fusiform Face Complex | 0.042413185 |
| Intra (Left) | MT+ Complex and Neighbors -- Ventral Stream | Left Area PH -- Left VentroMedial Visual Area 2 | 0.042570338 |
| Inter | Orbital and Polar Frontal Cortex -- Superior Parietal and IPS Cortex | Right Area 47m -- Left Medial Area 7P | 0.042634855 |
| Intra (Right) | Hippocampus -- Auditory Association Cortex | Right Subcortical Hippocampus -- Right Area STSv anterior | 0.042667001 |
| Inter | MT+ Complex and Neighbors -- Ventral Stream | Left Area FST -- Right VentroMedial Visual Area 3 | 0.043136279 |
| Intra (Left) | MT+ Complex and Neighbors -- Dorsal Stream | Left Area FST -- Left Area V3A | 0.043247358 |
| Intra (Right) | Caudate -- Inferior Parietal Cortex | Right Caudate -- Right Area PGp | 0.0440125 |
| Intra (Left) | MT+ Complex and Neighbors -- Dorsal Stream | Left Area PH -- Left Sixth Visual Area | 0.044093869 |
| Intra (Right) | Inferior Parietal Cortex -- Ventral Stream | Right Area PFt -- Right VentroMedial Visual Area 2 | 0.044440635 |
| Inter | Orbital and Polar Frontal Cortex -- Posterior Cingulate Cortex | Right Area 47s -- Left Area 31p ventral | 0.044489491 |
| Inter | Posterior Cingulate Cortex -- Ventral Stream | Left PreCuneus Visual Area -- Right Fusiform Face Complex | 0.044722343 |
| Intra (Left) | MT+ Complex and Neighbors -- Early Visual Cortex | Left Medial Superior Temporal Area -- Left Third Visual Area | 0.044816935 |
| Intra (Right) | Posterior Cingulate Cortex -- Sensorimotor Associated Paracentral Lobular and Mid Cingulate Cortex | Right ProStriate Area -- Right Area 5m | 0.044916655 |
| Inter | Orbital and Polar Frontal Cortex -- Posterior Cingulate Cortex | Left Area 10d -- Right Area ventral 23 a+b | 0.044966579 |
| Intra (Right) | Caudate -- Ventral Stream | Right Caudate -- Right VentroMedial Visual Area 1 | 0.045088249 |
| Intra (Right) | Caudate -- Inferior Frontal Cortex | Right Caudate -- Right Area IFJp | 0.045216338 |
| Inter | Posterior Cingulate Cortex -- Inferior Parietal Cortex | Right Area dorsal 23 a+b -- Left Area IntraParietal 2 | 0.045873955 |
| Inter | Superior Parietal and IPS Cortex -- Ventral Stream | Right Area Lateral IntraParietal ventral -- Left Ventral Visual Complex | 0.046131335 |
| Intra (Right) | Cerebellum -- Ventral Stream | Right Cerebellum -- Right VentroMedial Visual Area 2 | 0.046175301 |
| Intra (Right) | Posterior Cingulate Cortex -- Insular and Frontal Opercular Cortex | Right Area ventral 23 a+b -- Right Insular Granular Complex | 0.046267807 |
| Intra (Left) | Posterior Cingulate Cortex -- Dorsal Stream | Left Area 23c -- Left Seventh Visual Area | 0.046274894 |
| Intra (Right) | Sensorimotor Associated Paracentral Lobular and Mid Cingulate Cortex -- MT+ Complex and Neighbors | Right Area 5m ventral -- Right Area FST | 0.046296678 |
| Inter | Posterior Cingulate Cortex -- Lateral Temporal Cortex | Left Parieto-Occipital Sulcus Area 1 -- Right Area TF | 0.046821034 |
| Inter | Insular and Frontal Opercular Cortex -- Posterior Opercular Cortex | Left Insular Granular Complex -- Right Area 43 | 0.047097595 |
| Inter | Medial Temporal Cortex -- Sensorimotor Associated Paracentral Lobular and Mid Cingulate Cortex | Left Hippocampus -- Right Area 5m | 0.047281025 |
| Inter | Auditory Association Cortex -- Sensorimotor Associated Paracentral Lobular and Mid Cingulate Cortex | Right Auditory 4 Complex -- Left Area 5m | 0.047477816 |
| Intra (Left) | Posterior Opercular Cortex -- MT+ Complex and Neighbors | Left Area PFcm -- Left Middle Temporal Area | 0.047630683 |
| Inter | Posterior Cingulate Cortex -- Inferior Parietal Cortex | Left PreCuneus Visual Area -- Right Area PGp | 0.04771779 |
| Intra (Right) | Posterior Cingulate Cortex -- Ventral Stream | Right PreCuneus Visual Area -- Right VentroMedial Visual Area 1 | 0.04775633 |
| Intra (Left) | MT+ Complex and Neighbors -- Ventral Stream | Left Area Lateral Occipital 1 -- Left VentroMedial Visual Area 2 | 0.047991657 |
| Intra (Right) | Caudate -- Early Visual Cortex | Right Caudate -- Right Second Visual Area | 0.048535852 |
| Inter | Ventral Stream -- Ventral Stream | Left VentroMedial Visual Area 2 -- Right Fusiform Face Complex | 0.048606198 |
| Inter | MT+ Complex and Neighbors -- MT+ Complex and Neighbors | Left Area Lateral Occipital 2 -- Right Area FST | 0.049547258 |
| Intra (Right) | Caudate -- Medial Temporal Cortex | Right Caudate -- Right PreSubiculum | 0.04979455 |
| Intra (Left) | Insular and Frontal Opercular Cortex -- Dorsal Stream | Left Insular Granular Complex -- Left Sixth Visual Area | 0.049866788 |

**Table S1a:** Connections which were found to be significant across the group (MZ vs DZ) analysis. All connections were significant with the MZ variance smaller than DZ variance demonstrating genetic influence.

| Hemisphere Type | Network Interaction | Parcel Interaction | p-value |
|---|---|---|---|
| Intra (Left) | Posterior Cingulate Cortex -- Superior Parietal and IPS Cortex | Left Area 31a -- Left Medial Area 7P | 1.10E-09 |
| Inter | Ventral Stream -- Dorsal Stream | Right Eighth Visual Area -- Left IntraParietal Sulcus Area 1 | 9.00E-07 |
| Intra (Left) | Thalamus -- Inferior Parietal Cortex | Left Thalamus -- Left Area PGp | 1.15E-06 |
| Inter | MT+ Complex and Neighbors -- Dorsal Stream | Right Area Lateral Occipital 1 -- Left IntraParietal Sulcus Area 1 | 1.68E-06 |
| Inter | Thalamus -- Inferior Parietal Cortex | Left Thalamus -- Right Area PGp | 3.16E-06 |
| Inter | MT+ Complex and Neighbors -- Dorsal Stream | Right Area V3CD -- Left IntraParietal Sulcus Area 1 | 9.75E-06 |
| Intra (Left) | MT+ Complex and Neighbors -- MT+ Complex and Neighbors | Left Area PH -- Left Area V3CD | 1.18E-05 |
| Inter | Posterior Cingulate Cortex -- Lateral Temporal Cortex | Left Area ventral 23 a+b -- Right Area TG Ventral | 1.73E-05 |
| Intra (Right) | Posterior Cingulate Cortex -- Ventral Stream | Right PreCuneus Visual Area -- Right Fusiform Face Complex | 1.75E-05 |
| Intra (Right) | Posterior Cingulate Cortex -- Early Auditory Cortex | Right Area 7m -- Right Lateral Belt Complex | 2.78E-05 |
| Intra (Left) | Posterior Cingulate Cortex -- Superior Parietal and IPS Cortex | Left PreCuneus Visual Area -- Left Area Lateral IntraParietal dorsal | 3.09E-05 |
| Intra (Right) | Ventral Diencephalon -- Caudate | Right VentralDiencephalon -- Right Caudate | 3.21E-05 |
| Inter | Cerebellum -- Superior Parietal and IPS Cortex | Right Cerebellum -- Left Lateral Area 7A | 3.45E-05 |
| Inter | Pallidum -- Ventral Stream | Right Pallidum -- Left VentroMedial Visual Area 3 | 3.51E-05 |
| Inter | Thalamus -- Inferior Parietal Cortex | Left Thalamus -- Right Area IntraParietal 1 | 4.12E-05 |
| Inter | Pallidum -- MT+ Complex and Neighbors | Right Pallidum -- Left Area Lateral Occipital 1 | 6.04E-05 |
| Intra (Right) | Superior Parietal and IPS Cortex -- Premotor Cortex | Right Area Lateral IntraParietal dorsal -- Right Area 55b | 6.49E-05 |
| Intra (Left) | Thalamus -- Inferior Parietal Cortex | Left Thalamus -- Left Area IntraParietal 0 | 8.39E-05 |
| Inter | Dorsal Stream -- Early Visual Cortex | Left IntraParietal Sulcus Area 1 -- Right Fourth Visual Area | 0.000107303 |
| Inter | Thalamus -- Inferior Parietal Cortex | Left Thalamus -- Right Area IntraParietal 0 | 0.000109643 |
| Inter | MT+ Complex and Neighbors -- Dorsal Stream | Right Area PH -- Left IntraParietal Sulcus Area 1 | 0.000117248 |
| Intra (Right) | Auditory Association Cortex -- Early Auditory Cortex | Right Area STSd posterior -- Right Lateral Belt Complex | 0.000165819 |
| Intra (Left) | Posterior Cingulate Cortex -- Posterior Cingulate Cortex | Left Parieto-Occipital Sulcus Area 2 -- Left Area ventral 23 a+b | 0.000169432 |
| Inter | Superior Parietal and IPS Cortex -- Sensorimotor Associated Paracentral Lobular and Mid Cingulate Cortex | Left Area Lateral IntraParietal dorsal -- Right Area 5m | 0.000175083 |
| Intra (Left) | Dorsolateral Prefrontal Cortex -- Temporal-Parietal-Occipital Junction | Left Area 46 -- Left Area TemporoParietoOccipital Junction 3 | 0.000191932 |
| Intra (Left) | Thalamus -- Medial Temporal Cortex | Left Thalamus -- Left ParaHippocampal Area 3 | 0.000247183 |
| Intra (Left) | Thalamus -- Dorsal Stream | Left Thalamus -- Left IntraParietal Sulcus Area 1 | 0.000304739 |
| Inter | Hippocampus -- Temporal-Parietal-Occipital Junction | Right Subcortical Hippocampus -- Left PeriSylvian Language Area | 0.000332129 |
| Inter | Posterior Cingulate Cortex -- Posterior Cingulate Cortex | Left Parieto-Occipital Sulcus Area 1 -- Right Area dorsal 23 a+b | 0.000364955 |
| Intra (Left) | Superior Parietal and IPS Cortex -- MT+ Complex and Neighbors | Left Medial IntraParietal Area -- Left Area PH | 0.000400386 |
| Inter | Thalamus -- Dorsal Stream | Left Thalamus -- Right Area V3B | 0.000426118 |
| Inter | Superior Parietal and IPS Cortex -- Lateral Temporal Cortex | Left Area Lateral IntraParietal dorsal -- Right Area TE2 anterior | 0.000444364 |
| Inter | Temporal-Parietal-Occipital Junction -- Premotor Cortex | Right Area TemporoParietoOccipital Junction 1 -- Left Area 55b | 0.000447231 |
| Inter | Thalamus -- Ventral Stream | Left Thalamus -- Right Fusiform Face Complex | 0.000449746 |
| Intra (Left) | Superior Parietal and IPS Cortex -- Superior Parietal and IPS Cortex | Left Area 7PC -- Left Anterior IntraParietal Area | 0.000497132 |
| Inter | Posterior Cingulate Cortex -- MT+ Complex and Neighbors | Right PreCuneus Visual Area -- Left Area Lateral Occipital 2 | 0.000530186 |
| Intra (Right) | Cerebellum -- Ventral Stream | Right Cerebellum -- Right VentroMedial Visual Area 3 | 0.000605072 |
| Intra (Left) | Posterior Cingulate Cortex -- Temporal-Parietal-Occipital Junction | Left Area 23c -- Left Area TemporoParietoOccipital Junction 2 | 0.000607323 |
| Intra (Right) | Thalamus -- Dorsal Stream | Right Thalamus -- Right Area V3B | 0.000714895 |
| Inter | Orbital and Polar Frontal Cortex -- Posterior Cingulate Cortex | Right Area 47s -- Left Area 31pd | 0.000764092 |
| Inter | Posterior Cingulate Cortex -- Ventral Stream | Right PreCuneus Visual Area -- Left Posterior InferoTemporalComplex | 0.000814769 |
| Intra (Left) | Thalamus -- MT+ Complex and Neighbors | Left Thalamus -- Left Area Lateral Occipital 1 | 0.000886626 |
| Inter | Posterior Cingulate Cortex -- Superior Parietal and IPS Cortex | Right PreCuneus Visual Area -- Left Area Lateral IntraParietal dorsal | 0.000933154 |
| Intra (Left) | Insular and Frontal Opercular Cortex -- Auditory Association Cortex | Left Insular Granular Complex -- Left Area STSv anterior | 0.000943551 |
| Inter | Ventral Stream -- Dorsal Stream | Right VentroMedial Visual Area 2 -- Left Sixth Visual Area | 0.001051217 |
| Inter | Thalamus -- Superior Parietal and IPS Cortex | Left Thalamus -- Right Area Lateral IntraParietal dorsal | 0.001110455 |
| Inter | Posterior Cingulate Cortex -- Early Visual Cortex | Right PreCuneus Visual Area -- Left Fourth Visual Area | 0.001117524 |
| Intra (Right) | Auditory Association Cortex -- Early Auditory Cortex | Right Area STGa -- Right Lateral Belt Complex | 0.001151586 |
| Inter | Caudate -- MT+ Complex and Neighbors | Right Caudate -- Left Area PH | 0.001155994 |
| Intra (Left) | Inferior Parietal Cortex -- Temporal-Parietal-Occipital Junction | Left Area PF Complex -- Left Area TemporoParietoOccipital Junction 3 | 0.001212562 |
| Intra (Right) | Posterior Cingulate Cortex -- MT+ Complex and Neighbors | Right PreCuneus Visual Area -- Right Area V3CD | 0.001248706 |
| Intra (Left) | Temporal-Parietal-Occipital Junction -- Sensorimotor Associated Paracentral Lobular and Mid Cingulate Cortex | Left Area TemporoParietoOccipital Junction 2 -- Left Area 5m ventral | 0.001347971 |
| Intra (Right) | Caudate -- Dorsal Stream | Right Caudate -- Right Area V3B | 0.001406314 |
| Intra (Right) | Posterior Cingulate Cortex -- Temporal-Parietal-Occipital Junction | Right Parieto-Occipital Sulcus Area 1 -- Right Area TemporoParietoOccipital Junction 3 | 0.001439871 |
| Inter | Posterior Cingulate Cortex -- Insular and Frontal Opercular Cortex | Right Area 31pd -- Left Area 52 | 0.001445096 |
| Inter | Posterior Cingulate Cortex -- Posterior Cingulate Cortex | Left Area 23d -- Right Area 31pd | 0.001513966 |
| Inter | Inferior Frontal Cortex -- Premotor Cortex | Right Area 45 -- Left Area 55b | 0.001585721 |
| Inter | Caudate -- Inferior Parietal Cortex | Right Caudate -- Left Area IntraParietal 0 | 0.00171203 |
| Inter | Inferior Frontal Cortex -- Lateral Temporal Cortex | Left Area IFSp -- Right Area TE2 anterior | 0.001734764 |
| Intra (Left) | Superior Parietal and IPS Cortex -- Temporal-Parietal-Occipital Junction | Left Medial IntraParietal Area -- Left Area TemporoParietoOccipital Junction 3 | 0.001744028 |
| Intra (Left) | Inferior Frontal Cortex -- Inferior Parietal Cortex | Left Area 47l (47 lateral) -- Left Area PGi | 0.001763679 |
| Intra (Left) | Thalamus -- Lateral Temporal Cortex | Left Thalamus -- Left Area TF | 0.001895652 |
| Intra (Left) | Posterior Cingulate Cortex -- Insular and Frontal Opercular Cortex | Left Area 31p ventral -- Left Anterior Agranular Insula Complex | 0.001903497 |
| Intra (Right) | Posterior Cingulate Cortex -- Inferior Parietal Cortex | Right Area dorsal 23 a+b -- Right Area IntraParietal 1 | 0.001912874 |
| Intra (Left) | Posterior Cingulate Cortex -- Temporal-Parietal-Occipital Junction | Left Area 23c -- Left Area TemporoParietoOccipital Junction 3 | 0.001954754 |
| Intra (Right) | Posterior Cingulate Cortex -- Ventral Stream | Right PreCuneus Visual Area -- Right Posterior InferoTemporalComplex | 0.00199484 |
| Inter | Posterior Cingulate Cortex -- MT+ Complex and Neighbors | Right PreCuneus Visual Area -- Left Area PH | 0.002031828 |
| Inter | Ventral Diencephalon -- Superior Parietal and IPS Cortex | Left VentralDiencephalon -- Right Area Lateral IntraParietal dorsal | 0.002034244 |
| Inter | Anterior Cingulate and Medial Prefrontal Cortex -- Posterior Cingulate Cortex | Right Area a24 -- Left Parieto-Occipital Sulcus Area 2 | 0.002078719 |
| Inter | Sensorimotor Associated Paracentral Lobular and Mid Cingulate Cortex -- Dorsal Stream | Left Area 5m -- Right Area V6A | 0.002118687 |
| Intra (Right) | Posterior Cingulate Cortex -- Posterior Opercular Cortex | Right Area 7m -- Right Area OP2-3/VS | 0.00212665 |
| Inter | MT+ Complex and Neighbors -- MT+ Complex and Neighbors | Left Area PH -- Right Area V3CD | 0.002353499 |
| Inter | Caudate -- Medial Temporal Cortex | Right Caudate -- Left PreSubiculum | 0.002473829 |
| Intra (Left) | Posterior Cingulate Cortex -- Dorsal Stream | Left Area 23c -- Left Seventh Visual Area | 0.002484608 |
| Inter | Thalamus -- Temporal-Parietal-Occipital Junction | Left Thalamus -- Right Area TemporoParietoOccipital Junction 1 | 0.002486936 |
| Inter | Temporal-Parietal-Occipital Junction -- Premotor Cortex | Right Superior Temporal Visual Area -- Left Area 55b | 0.002627273 |
| Intra (Right) | Temporal-Parietal-Occipital Junction -- Lateral Temporal Cortex | Right Area TemporoParietoOccipital Junction 2 -- Right Area TG Ventral | 0.002661369 |
| Inter | Thalamus -- MT+ Complex and Neighbors | Left Thalamus -- Right Middle Temporal Area | 0.002742856 |
| Inter | Hippocampus -- Superior Parietal and IPS Cortex | Right Subcortical Hippocampus -- Left Medial Area 7A | 0.002807414 |
| Intra (Right) | Auditory Association Cortex -- Early Auditory Cortex | Right Area STSd posterior -- Right RetroInsular Cortex | 0.002846206 |
| Inter | Cerebellum -- Medial Temporal Cortex | Right Cerebellum -- Left ParaHippocampal Area 2 | 0.002880719 |
| Intra (Right) | Lateral Temporal Cortex -- Medial Temporal Cortex | Right Area TE2 anterior -- Right PreSubiculum | 0.003017235 |
| Intra (Left) | Insular and Frontal Opercular Cortex -- Premotor Cortex | Left Frontal Opercular Area 3 -- Left Rostral Area 6 | 0.003017671 |
| Inter | Thalamus -- MT+ Complex and Neighbors | Right Thalamus -- Left Area Lateral Occipital 1 | 0.003018886 |
| Intra (Left) | Orbital and Polar Frontal Cortex -- Inferior Parietal Cortex | Left Area 13l -- Left Area PFm Complex | 0.003063243 |
| Inter | Thalamus -- Dorsal Stream | Left Thalamus -- Right IntraParietal Sulcus Area 1 | 0.00307056 |
| Inter | Superior Parietal and IPS Cortex -- Sensorimotor Associated Paracentral Lobular and Mid Cingulate Cortex | Left Area Lateral IntraParietal dorsal -- Right Area 5L | 0.003095949 |
| Intra (Left) | Superior Parietal and IPS Cortex -- Dorsal Stream | Left Area Lateral IntraParietal ventral -- Left IntraParietal Sulcus Area 1 | 0.003097067 |
| Intra (Right) | Superior Parietal and IPS Cortex -- Early Auditory Cortex | Right Area Lateral IntraParietal dorsal -- Right Lateral Belt Complex | 0.003197894 |
| Intra (Right) | Caudate -- Superior Parietal and IPS Cortex | Right Caudate -- Right Area Lateral IntraParietal ventral | 0.003386304 |
| Intra (Left) | Temporal-Parietal-Occipital Junction -- Sensorimotor Associated Paracentral Lobular and Mid Cingulate Cortex | Left Area TemporoParietoOccipital Junction 3 -- Left Area 5m ventral | 0.00343619 |
| Intra (Left) | Thalamus -- Inferior Parietal Cortex | Left Thalamus -- Left Area PFm Complex | 0.003487907 |
| Inter | Posterior Cingulate Cortex -- Ventral Stream | Right PreCuneus Visual Area -- Left Fusiform Face Complex | 0.003524046 |
| Inter | Ventral Diencephalon -- Superior Parietal and IPS Cortex | Left VentralDiencephalon -- Right Anterior IntraParietal Area | 0.003572208 |
| Intra (Right) | Posterior Cingulate Cortex -- Inferior Parietal Cortex | Right Area ventral 23 a+b -- Right Area IntraParietal 1 | 0.003590461 |
| Intra (Left) | Medial Temporal Cortex -- Early Visual Cortex | Left ParaHippocampal Area 3 -- Left Fourth Visual Area | 0.003692144 |
| Intra (Left) | Auditory Association Cortex -- Auditory Association Cortex | Left Area STSd posterior -- Left Area STSv anterior | 0.003719235 |
| Inter | Posterior Cingulate Cortex -- Inferior Parietal Cortex | Right ProStriate Area -- Left Area IntraParietal 0 | 0.003776163 |
| Intra (Right) | Posterior Cingulate Cortex -- Early Visual Cortex | Right PreCuneus Visual Area -- Right Second Visual Area | 0.003805634 |
| Intra (Right) | Cerebellum -- Medial Temporal Cortex | Right Cerebellum -- Right ParaHippocampal Area 1 | 0.003837067 |
| Intra (Right) | Posterior Cingulate Cortex -- Posterior Cingulate Cortex | Right Area 7m -- Right Area dorsal 23 a+b | 0.003851689 |
| Intra (Left) | Posterior Cingulate Cortex -- Insular and Frontal Opercular Cortex | Left Area 23c -- Left Area Posterior Insular 1 | 0.003943413 |
| Inter | Pallidum -- Auditory Association Cortex | Right Pallidum -- Left Area STSd posterior | 0.003983215 |
| Inter | Early Auditory Cortex -- Dorsal Stream | Right RetroInsular Cortex -- Left IntraParietal Sulcus Area 1 | 0.004111043 |
| Inter | Posterior Cingulate Cortex -- MT+ Complex and Neighbors | Left Area 23c -- Right Area FST | 0.004213074 |
| Intra (Right) | Posterior Cingulate Cortex -- Lateral Temporal Cortex | Right Area 7m -- Right Area TG Ventral | 0.004367373 |
| Inter | Thalamus -- Lateral Temporal Cortex | Left Thalamus -- Right Area TF | 0.004414402 |
| Inter | Inferior Parietal Cortex -- Inferior Parietal Cortex | Left Area IntraParietal 2 -- Right Area PGs | 0.004456431 |
| Inter | Pallidum -- Temporal-Parietal-Occipital Junction | Right Pallidum -- Left Area TemporoParietoOccipital Junction 2 | 0.004544059 |
| Inter | Thalamus -- MT+ Complex and Neighbors | Left Thalamus -- Right Area V3CD | 0.004557493 |
| Intra (Left) | MT+ Complex and Neighbors -- Dorsal Stream | Left Area PH -- Left IntraParietal Sulcus Area 1 | 0.004561469 |
| Inter | MT+ Complex and Neighbors -- Dorsal Stream | Left Area PH -- Right IntraParietal Sulcus Area 1 | 0.004564829 |
| Inter | Thalamus -- MT+ Complex and Neighbors | Left Thalamus -- Right Area PH | 0.004575991 |
| Inter | Posterior Cingulate Cortex -- Premotor Cortex | Right PreCuneus Visual Area -- Left Area 55b | 0.004713629 |
| Intra (Right) | Posterior Cingulate Cortex -- Inferior Parietal Cortex | Right Area ventral 23 a+b -- Right Area PFm Complex | 0.004830782 |
| Inter | Posterior Cingulate Cortex -- Early Visual Cortex | Right PreCuneus Visual Area -- Left Second Visual Area | 0.005263715 |
| Intra (Right) | Caudate -- Superior Parietal and IPS Cortex | Right Caudate -- Right Area Lateral IntraParietal dorsal | 0.005270293 |
| Intra (Right) | Superior Parietal and IPS Cortex -- MT+ Complex and Neighbors | Right Area Lateral IntraParietal ventral -- Right Area PH | 0.005408494 |
| Intra (Left) | Dorsal Stream -- Dorsal Stream | Left Seventh Visual Area -- Left IntraParietal Sulcus Area 1 | 0.005408787 |
| Inter | Dorsolateral Prefrontal Cortex -- Inferior Parietal Cortex | Left Area 9 Posterior -- Right Area PFm Complex | 0.005411051 |
| Inter | Posterior Cingulate Cortex -- MT+ Complex and Neighbors | Left Area 23c -- Right Area Lateral Occipital 1 | 0.005451688 |
| Inter | Ventral Diencephalon -- Dorsolateral Prefrontal Cortex | Left VentralDiencephalon -- Right Area 9-46d | 0.005473637 |
| Inter | Posterior Cingulate Cortex -- MT+ Complex and Neighbors | Left Area 23c -- Right Area Lateral Occipital 3 | 0.0055608 |
| Inter | Sensorimotor Associated Paracentral Lobular and Mid Cingulate Cortex -- Sensorimotor Associated Paracentral Lobular and Mid Cingulate Cortex | Left Area 5m -- Right Ventral Area 24d | 0.005593593 |
| Intra (Right) | Inferior Frontal Cortex -- Inferior Frontal Cortex | Right Area 44 -- Right Area 45 | 0.005810723 |
| Intra (Left) | Thalamus -- Inferior Parietal Cortex | Left Thalamus -- Left Area IntraParietal 1 | 0.005860667 |
| Intra (Right) | Orbital and Polar Frontal Cortex -- Inferior Parietal Cortex | Right Area 47m -- Right Area IntraParietal 1 | 0.005981579 |
| Intra (Left) | Auditory Association Cortex -- Auditory Association Cortex | Left Area STSv posterior -- Left Area STSv anterior | 0.006019871 |
| Inter | Posterior Cingulate Cortex -- Temporal-Parietal-Occipital Junction | Right PreCuneus Visual Area -- Left Area TemporoParietoOccipital Junction 3 | 0.006060468 |
| Inter | Auditory Association Cortex -- Early Auditory Cortex | Right Area STSd posterior -- Left Lateral Belt Complex | 0.006073867 |
| Intra (Right) | Posterior Cingulate Cortex -- Dorsal Stream | Right PreCuneus Visual Area -- Right Area V3B | 0.006243662 |
| Intra (Left) | Temporal-Parietal-Occipital Junction -- Early Auditory Cortex | Left PeriSylvian Language Area -- Left ParaBelt Complex | 0.006302779 |
| Intra (Left) | Posterior Cingulate Cortex -- Medial Temporal Cortex | Left Area ventral 23 a+b -- Left Perirhinal Ectorhinal Cortex | 0.006452943 |
| Intra (Right) | Orbital and Polar Frontal Cortex -- Lateral Temporal Cortex | Right Area 47m -- Right Area PHT | 0.00668358 |
| Inter | Posterior Cingulate Cortex -- Inferior Parietal Cortex | Right Area ventral 23 a+b -- Left Area IntraParietal 0 | 0.006870612 |
| Inter | Inferior Parietal Cortex -- Sensorimotor Associated Paracentral Lobular and Mid Cingulate Cortex | Left Area IntraParietal 0 -- Right Area 5m ventral | 0.007213346 |
| Intra (Right) | Posterior Cingulate Cortex -- Ventral Stream | Right PreCuneus Visual Area -- Right Eighth Visual Area | 0.007287201 |
| Intra (Left) | Temporal-Parietal-Occipital Junction -- Early Auditory Cortex | Left PeriSylvian Language Area -- Left Lateral Belt Complex | 0.007582741 |
| Inter | Orbital and Polar Frontal Cortex -- Posterior Cingulate Cortex | Right Area 47s -- Left Area 31p ventral | 0.007583559 |
| Inter | Dorsal Stream -- Early Visual Cortex | Right Area V6A -- Left Third Visual Area | 0.007658276 |
| Inter | Cerebellum -- Sensorimotor Associated Paracentral Lobular and Mid Cingulate Cortex | Right Cerebellum -- Left Dorsal Area 24d | 0.007730005 |
| Intra (Right) | Auditory Association Cortex -- Early Auditory Cortex | Right Area STSd posterior -- Right Primary Auditory Cortex | 0.007837311 |
| Intra (Left) | Amygdala -- Inferior Parietal Cortex | Left Amygdala -- Left Area IntraParietal 2 | 0.008002046 |
| Intra (Right) | Caudate -- MT+ Complex and Neighbors | Right Caudate -- Right Area PH | 0.008368838 |
| Inter | Posterior Cingulate Cortex -- Insular and Frontal Opercular Cortex | Right Area ventral 23 a+b -- Left Anterior Agranular Insula Complex | 0.008727748 |
| Inter | Posterior Cingulate Cortex -- Early Visual Cortex | Right PreCuneus Visual Area -- Left Third Visual Area | 0.008793927 |
| Inter | Temporal-Parietal-Occipital Junction -- Insular and Frontal Opercular Cortex | Right Area TemporoParietoOccipital Junction 1 -- Left Insular Granular Complex | 0.008972572 |
| Inter | Orbital and Polar Frontal Cortex -- Posterior Cingulate Cortex | Right Area 47m -- Left Parieto-Occipital Sulcus Area 2 | 0.009461957 |
| Intra (Right) | Inferior Parietal Cortex -- MT+ Complex and Neighbors | Right Area IntraParietal 0 -- Right Area PH | 0.009525786 |
| Intra (Left) | Thalamus -- Superior Parietal and IPS Cortex | Left Thalamus -- Left Lateral Area 7A | 0.009586041 |
| Intra (Right) | Posterior Cingulate Cortex -- Inferior Parietal Cortex | Right Parieto-Occipital Sulcus Area 1 -- Right Area IntraParietal 1 | 0.009709635 |
| Intra (Right) | Ventral Diencephalon -- Posterior Cingulate Cortex | Right VentralDiencephalon -- Right Area 23d | 0.009786527 |
| Intra (Right) | Superior Parietal and IPS Cortex -- Posterior Opercular Cortex | Right Area Lateral IntraParietal ventral -- Right Area PFcm | 0.010192614 |
| Inter | Lateral Temporal Cortex -- MT+ Complex and Neighbors | Right Area TG Ventral -- Left Area V4t | 0.010299265 |
| Inter | Thalamus -- MT+ Complex and Neighbors | Left Thalamus -- Right Area FST | 0.010308352 |
| Inter | Anterior Cingulate and Medial Prefrontal Cortex -- Posterior Cingulate Cortex | Left Area 10r -- Right Area ventral 23 a+b | 0.010831069 |
| Inter | Posterior Cingulate Cortex -- Posterior Cingulate Cortex | Left Parieto-Occipital Sulcus Area 2 -- Right Area ventral 23 a+b | 0.011041223 |
| Inter | Posterior Cingulate Cortex -- Inferior Parietal Cortex | Right Area dorsal 23 a+b -- Left Area PGs | 0.011123266 |
| Inter | Thalamus -- Lateral Temporal Cortex | Left Thalamus -- Right Area PHT | 0.011160759 |
| Intra (Left) | Dorsolateral Prefrontal Cortex -- Premotor Cortex | Left Superior 6-8 Translational Area -- Left Frontal Eye Fields | 0.011232641 |
| Intra (Left) | Inferior Parietal Cortex -- Lateral Temporal Cortex | Left Area IntraParietal 1 -- Left Area PHT | 0.011234083 |
| Intra (Right) | Caudate -- Posterior Cingulate Cortex | Right Caudate -- Right Dorsal Transitional Visual Area | 0.011306273 |

| | | | |
|---|---|---|---|
| Inter | Thalamus -- Inferior Parietal Cortex | Left Thalamus -- Right Area PFm Complex | 0.011372148 |
| Inter | Caudate -- Sensorimotor Associated Paracentral Lobular and Mid Cingulate Cortex | Right Caudate -- Left Dorsal Area 24d | 0.011585256 |
| Inter | Dorsal Stream -- Early Visual Cortex | Right Area V6A -- Left Second Visual Area | 0.01165694 |
| Inter | Posterior Cingulate Cortex -- Auditory Association Cortex | Right Area dorsal 23 a+b -- Left Area STSd posterior | 0.012234247 |
| Inter | Anterior Cingulate and Medial Prefrontal Cortex -- Posterior Cingulate Cortex | Left Area 25 -- Right Area ventral 23 a+b | 0.01240639 |
| Intra (Left) | Sensorimotor Associated Paracentral Lobular and Mid Cingulate Cortex -- Dorsal Stream | Left Area 5m -- Left Area V6A | 0.012643651 |
| Inter | Ventral Stream -- Dorsal Stream | Right VentroMedial Visual Area 3 -- Left IntraParietal Sulcus Area 1 | 0.012651615 |
| Intra (Right) | Orbital and Polar Frontal Cortex -- Inferior Parietal Cortex | Right Area 47m -- Right Area PFm Complex | 0.012902874 |
| Inter | Ventral Stream -- Early Visual Cortex | Right VentroMedial Visual Area 2 -- Left Second Visual Area | 0.013047731 |
| Inter | Posterior Cingulate Cortex -- Auditory Association Cortex | Right Area dorsal 23 a+b -- Left Area STSv posterior | 0.013132418 |
| Intra (Left) | Ventral Stream -- Dorsal Stream | Left Eighth Visual Area -- Left IntraParietal Sulcus Area 1 | 0.013438881 |
| Intra (Left) | Thalamus -- Auditory Association Cortex | Left Thalamus -- Left Area STSd anterior | 0.013575532 |
| Intra (Left) | Dorsolateral Prefrontal Cortex -- Lateral Temporal Cortex | Left Superior 6-8 Translational Area -- Left Area TE2 posterior | 0.013736694 |
| Intra (Right) | Lateral Temporal Cortex -- MT+ Complex and Neighbors | Right Area TF -- Right Area FST | 0.013921216 |
| Inter | Orbital and Polar Frontal Cortex -- Lateral Temporal Cortex | Right Area 13l -- Left Area PHT | 0.013949749 |
| Intra (Right) | Posterior Cingulate Cortex -- MT+ Complex and Neighbors | Right PreCuneus Visual Area -- Right Area V4t | 0.013959835 |
| Inter | Anterior Cingulate and Medial Prefrontal Cortex -- Inferior Parietal Cortex | Left Area Posterior 24 prime -- Right Area PFt | 0.014347654 |
| Inter | Posterior Cingulate Cortex -- Superior Parietal and IPS Cortex | Right Area ventral 23 a+b -- Left Area Lateral IntraParietal dorsal | 0.014411649 |
| Inter | MT+ Complex and Neighbors -- Dorsal Stream | Right Area Lateral Occipital 1 -- Left Seventh Visual Area | 0.014552691 |
| Intra (Left) | Medial Temporal Cortex -- Premotor Cortex | Left ParaHippocampal Area 3 -- Left Premotor Eye Field | 0.014656579 |
| Intra (Left) | Dorsolateral Prefrontal Cortex -- Orbital and Polar Frontal Cortex | Left Inferior 6-8 Transitional Area -- Left Area 13l | 0.014682655 |
| Intra (Left) | Superior Parietal and IPS Cortex -- Temporal-Parietal-Occipital Junction | Left Medial Area 7A -- Left Area TemporoParietoOccipital Junction 2 | 0.014684371 |
| Intra (Right) | Temporal-Parietal-Occipital Junction -- Auditory Association Cortex | Right Area TemporoParietoOccipital Junction 2 -- Right Area STGa | 0.014761189 |
| Intra (Right) | Superior Parietal and IPS Cortex -- Medial Temporal Cortex | Right Area Lateral IntraParietal dorsal -- Right PreSubiculum | 0.014905791 |
| Intra (Right) | Posterior Cingulate Cortex -- Inferior Parietal Cortex | Right Area 7m -- Right Area PGi | 0.015184775 |
| Inter | Dorsolateral Prefrontal Cortex -- Temporal-Parietal-Occipital Junction | Left Area 8Ad -- Right Area TemporoParietoOccipital Junction 3 | 0.015569795 |
| Intra (Left) | Anterior Cingulate and Medial Prefrontal Cortex -- Inferior Parietal Cortex | Left Area dorsal 32 -- Left Area IntraParietal 2 | 0.015839619 |
| Inter | Sensorimotor Associated Paracentral Lobular and Mid Cingulate Cortex -- MT+ Complex and Neighbors | Right Area 5m ventral -- Left Area PH | 0.015919509 |
| Intra (Right) | Sensorimotor Associated Paracentral Lobular and Mid Cingulate Cortex -- Ventral Stream | Right Area 5m ventral -- Right Fusiform Face Complex | 0.01598703 |
| Intra (Right) | Posterior Cingulate Cortex -- MT+ Complex and Neighbors | Right PreCuneus Visual Area -- Right Area Lateral Occipital 3 | 0.016243892 |
| Intra (Left) | Posterior Cingulate Cortex -- Posterior Cingulate Cortex | Left Area 23d -- Left Area 31pd | 0.016274221 |
| Inter | Ventral Diencephalon -- Inferior Frontal Cortex | Left VentralDiencephalon -- Right Area posterior 47r | 0.016299218 |
| Inter | Cerebellum -- Medial Temporal Cortex | Right Cerebellum -- Left ParaHippocampal Area 1 | 0.016442314 |
| Inter | Ventral Diencephalon -- Inferior Parietal Cortex | Left VentralDiencephalon -- Right Area PF Complex | 0.016516537 |
| Intra (Left) | Thalamus -- Posterior Cingulate Cortex | Left Thalamus -- Left ProStriate Area | 0.016613717 |
| Intra (Left) | Posterior Cingulate Cortex -- Temporal-Parietal-Occipital Junction | Left PreCuneus Visual Area -- Left Area TemporoParietoOccipital Junction 3 | 0.016781432 |
| Inter | Auditory Association Cortex -- Early Auditory Cortex | Left Area STGa -- Right Lateral Belt Complex | 0.016799603 |
| Intra (Left) | Sensorimotor Associated Paracentral Lobular and Mid Cingulate Cortex -- Dorsal Stream | Left Area 5m -- Left Sixth Visual Area | 0.017160641 |
| Inter | MT+ Complex and Neighbors -- Dorsal Stream | Right Area FST -- Left IntraParietal Sulcus Area 1 | 0.017324574 |
| Intra (Right) | Hippocampus -- Posterior Opercular Cortex | Right Subcortical Hippocampus -- Right Area PFcm | 0.017563295 |
| Intra (Right) | Posterior Cingulate Cortex -- Insular and Frontal Opercular Cortex | Right Parieto-Occipital Sulcus Area 1 -- Right Area Frontal Opercular 5 | 0.017622447 |
| Intra (Left) | Superior Parietal and IPS Cortex -- MT+ Complex and Neighbors | Left Area Lateral IntraParietal ventral -- Left Area PH | 0.017902281 |
| Intra (Right) | Posterior Cingulate Cortex -- Premotor Cortex | Right ProStriate Area -- Right Area 6 anterior | 0.017994703 |
| Inter | Temporal-Parietal-Occipital Junction -- Insular and Frontal Opercular Cortex | Right Area TemporoParietoOccipital Junction 2 -- Left Pirform Cortex | 0.018119811 |
| Intra (Right) | Dorsolateral Prefrontal Cortex -- Posterior Cingulate Cortex | Right Area posterior 9-46v -- Right Area ventral 23 a+b | 0.019934818 |
| Intra (Right) | Ventral Diencephalon -- Auditory Association Cortex | Right VentralDiencephalon -- Right Area STSv posterior | 0.020017355 |
| Intra (Left) | Hippocampus -- Temporal-Parietal-Occipital Junction | Left Subcortical Hippocampus -- Left PeriSylvian Language Area | 0.020224821 |
| Intra (Left) | Insular and Frontal Opercular Cortex -- Posterior Opercular Cortex | Left Insular Granular Complex -- Left Area OP4/PV | 0.020331729 |
| Intra (Right) | MT+ Complex and Neighbors -- Ventral Stream | Right Area PH -- Right VentroMedial Visual Area 2 | 0.021366779 |
| Intra (Right) | Posterior Cingulate Cortex -- Early Auditory Cortex | Right Area ventral 23 a+b -- Right Lateral Belt Complex | 0.021582484 |
| Inter | Orbital and Polar Frontal Cortex -- Lateral Temporal Cortex | Right Area 47m -- Left Area PHT | 0.021709832 |
| Inter | Cerebellum -- Dorsal Stream | Right Cerebellum -- Left IntraParietal Sulcus Area 1 | 0.021961341 |
| Inter | Superior Parietal and IPS Cortex -- Temporal-Parietal-Occipital Junction | Right Anterior IntraParietal Area -- Left Area TemporoParietoOccipital Junction 3 | 0.022307546 |
| Intra (Right) | Caudate -- Lateral Temporal Cortex | Right Caudate -- Right Area TE2 posterior | 0.022458101 |
| Intra (Left) | Dorsal Stream -- Early Visual Cortex | Left IntraParietal Sulcus Area 1 -- Left Fourth Visual Area | 0.022600379 |
| Inter | Anterior Cingulate and Medial Prefrontal Cortex -- Superior Parietal and IPS Cortex | Left Anterior 24 prime -- Right Area Lateral IntraParietal dorsal | 0.022711856 |
| Intra (Left) | Superior Parietal and IPS Cortex -- Lateral Temporal Cortex | Left Medial Area 7A -- Left Area PHT | 0.023135698 |
| Inter | Posterior Cingulate Cortex -- Ventral Stream | Right PreCuneus Visual Area -- Left Ventral Visual Complex | 0.023285294 |
| Intra (Left) | Temporal-Parietal-Occipital Junction -- Sensorimotor Associated Paracentral Lobular and Mid Cingulate Cortex | Left Area TemporoParietoOccipital Junction 1 -- Left Area 5m ventral | 0.023309898 |
| Intra (Left) | Thalamus -- Auditory Association Cortex | Left Thalamus -- Left Area STSd posterior | 0.023821801 |
| Intra (Right) | Posterior Cingulate Cortex -- Early Auditory Cortex | Right Area 7m -- Right RetroInsular Cortex | 0.024304683 |
| Intra (Right) | Inferior Parietal Cortex -- Early Visual Cortex | Right Area IntraParietal 0 -- Right Fourth Visual Area | 0.024478398 |
| Intra (Right) | Inferior Parietal Cortex -- Ventral Stream | Right Area IntraParietal 0 -- Right Eighth Visual Area | 0.024502995 |
| Intra (Right) | Lateral Temporal Cortex -- Sensorimotor Associated Paracentral Lobular and Mid Cingulate Cortex | Right Area TG Ventral -- Right Area 5L | 0.024716371 |
| Inter | Orbital and Polar Frontal Cortex -- Inferior Parietal Cortex | Left Area 13l -- Right Area PFm Complex | 0.024762329 |
| Inter | Thalamus -- Lateral Temporal Cortex | Left Thalamus -- Right Area TE2 anterior | 0.024838596 |
| Intra (Left) | Medial Temporal Cortex -- MT+ Complex and Neighbors | Left ParaHippocampal Area 3 -- Left Area Lateral Occipital 2 | 0.025052215 |
| Intra (Right) | Pallidum -- MT+ Complex and Neighbors | Right Pallidum -- Right Area V3CD | 0.025243193 |
| Intra (Left) | Medial Temporal Cortex -- Ventral Stream | Left ParaHippocampal Area 3 -- Left Posterior InferoTemporalComplex | 0.025264265 |
| Intra (Right) | Temporal-Parietal-Occipital Junction -- Ventral Stream | Right Superior Temporal Visual Area -- Right Ventral Visual Complex | 0.025480383 |
| Intra (Right) | Inferior Frontal Cortex -- Posterior Cingulate Cortex | Right Area 45 -- Right Area 7m | 0.02597475 |
| Inter | Posterior Cingulate Cortex -- Auditory Association Cortex | Right RetroSplenial Complex -- Left Area STSd posterior | 0.026511419 |
| Intra (Left) | Ventral Diencephalon -- Dorsal Stream | Left VentralDiencephalon -- Left IntraParietal Sulcus Area 1 | 0.027192238 |
| Intra (Left) | Auditory Association Cortex -- Sensorimotor Associated Paracentral Lobular and Mid Cingulate Cortex | Left Auditory 4 Complex -- Left Area 5m ventral | 0.027469692 |
| Intra (Right) | Posterior Cingulate Cortex -- Posterior Cingulate Cortex | Right RetroSplenial Complex -- Right Area 7m | 0.027603751 |
| Intra (Right) | Orbital and Polar Frontal Cortex -- Inferior Parietal Cortex | Right Area 10d -- Right Area IntraParietal 1 | 0.027645309 |
| Intra (Right) | Posterior Cingulate Cortex -- Early Auditory Cortex | Right Parieto-Occipital Sulcus Area 1 -- Right Lateral Belt Complex | 0.028297557 |
| Inter | Posterior Cingulate Cortex -- Inferior Parietal Cortex | Right Area 23d -- Left Area IntraParietal 0 | 0.028340699 |
| Inter | Orbital and Polar Frontal Cortex -- Lateral Temporal Cortex | Right Area 11l -- Left Area TE1 posterior | 0.028577579 |
| Intra (Left) | Ventral Diencephalon -- Superior Parietal and IPS Cortex | Left VentralDiencephalon -- Left Lateral Area 7P | 0.029222853 |
| Inter | Posterior Cingulate Cortex -- Early Auditory Cortex | Right Parieto-Occipital Sulcus Area 1 -- Left Lateral Belt Complex | 0.029803093 |
| Intra (Right) | Posterior Cingulate Cortex -- Medial Temporal Cortex | Right Area 7m -- Right Entorhinal Cortex | 0.030222154 |
| Inter | Posterior Cingulate Cortex -- MT+ Complex and Neighbors | Left Area 23c -- Right Area Lateral Occipital 2 | 0.030520986 |
| Intra (Right) | Posterior Cingulate Cortex -- Early Visual Cortex | Right PreCuneus Visual Area -- Right Fourth Visual Area | 0.030672605 |
| Intra (Left) | Superior Parietal and IPS Cortex -- Sensorimotor Associated Paracentral Lobular and Mid Cingulate Cortex | Left Area Lateral IntraParietal dorsal -- Left Ventral Area 24d | 0.030757974 |
| Inter | Ventral Stream -- Dorsal Stream | Right Posterior InferoTemporalComplex -- Left IntraParietal Sulcus Area 1 | 0.030868837 |
| Intra (Right) | Superior Parietal and IPS Cortex -- Primary Visual Cortex (V1) | Right Area Lateral IntraParietal dorsal -- Right Primary Visual Cortex | 0.03093699 |
| Intra (Right) | Accumbens -- Anterior Cingulate and Medial Prefrontal Cortex | Right Accumbens -- Right Anterior 24 prime | 0.031009553 |
| Inter | Orbital and Polar Frontal Cortex -- Inferior Parietal Cortex | Right Area 13l -- Left Area PF Complex | 0.031051585 |
| Inter | Insular and Frontal Opercular Cortex -- Posterior Opercular Cortex | Left Insular Granular Complex -- Right Area OP1/SII | 0.031494142 |
| Inter | Medial Temporal Cortex -- Sensorimotor Associated Paracentral Lobular and Mid Cingulate Cortex | Left PreSubiculum -- Right Area 5m | 0.031769476 |
| Intra (Left) | Ventral Diencephalon -- Premotor Cortex | Left VentralDiencephalon -- Left Area 6 anterior | 0.032027985 |
| Inter | Dorsal Stream -- Early Visual Cortex | Left IntraParietal Sulcus Area 1 -- Right Third Visual Area | 0.032232861 |
| Intra (Right) | Posterior Cingulate Cortex -- Temporal-Parietal-Occipital Junction | Right Area ventral 23 a+b -- Right Area TemporoParietoOccipital Junction 3 | 0.032264873 |
| Intra (Right) | Posterior Cingulate Cortex -- MT+ Complex and Neighbors | Right Area ventral 23 a+b -- Right Area PH | 0.03234816 |
| Inter | Ventral Stream -- Dorsal Stream | Right Ventral Visual Complex -- Left IntraParietal Sulcus Area 1 | 0.032899783 |
| Inter | Lateral Temporal Cortex -- Auditory Association Cortex | Right Area TG Ventral -- Left Auditory 4 Complex | 0.033058729 |
| Intra (Right) | Dorsolateral Prefrontal Cortex -- Orbital and Polar Frontal Cortex | Right Area 8Av -- Right Area 47m | 0.033131167 |
| Intra (Right) | Auditory Association Cortex -- Premotor Cortex | Right Area STGa -- Right Area 6 anterior | 0.033160915 |
| Inter | Posterior Opercular Cortex -- Premotor Cortex | Right Area PFcm -- Left Frontal Eye Fields | 0.033598481 |
| Intra (Right) | Dorsolateral Prefrontal Cortex -- Inferior Parietal Cortex | Right Area 9 Posterior -- Right Area IntraParietal 1 | 0.033756536 |
| Intra (Right) | Dorsolateral Prefrontal Cortex -- Inferior Frontal Cortex | Right Area 46 -- Right Area IFSa | 0.0337673 |
| Inter | Orbital and Polar Frontal Cortex -- Posterior Cingulate Cortex | Right Area 10d -- Left Parieto-Occipital Sulcus Area 2 | 0.034081121 |
| Intra (Right) | Cerebellum -- MT+ Complex and Neighbors | Right Cerebellum -- Right Area PH | 0.034190762 |
| Inter | Posterior Cingulate Cortex -- Inferior Parietal Cortex | Right Area 7m -- Left Area IntraParietal 2 | 0.034697437 |
| Intra (Left) | Auditory Association Cortex -- Sensorimotor Associated Paracentral Lobular and Mid Cingulate Cortex | Left Auditory 4 Complex -- Left Area 5m | 0.03480785 |
| Intra (Right) | Posterior Cingulate Cortex -- Dorsal Stream | Right RetroSplenial Complex -- Right Area V3B | 0.034944286 |
| Intra (Right) | Early Auditory Cortex -- Ventral Stream | Right RetroInsular Cortex -- Right Posterior InferoTemporalComplex | 0.035006596 |
| Inter | Hippocampus -- Sensorimotor Associated Paracentral Lobular and Mid Cingulate Cortex | Right Subcortical Hippocampus -- Left Area 5L | 0.035051122 |
| Inter | Posterior Cingulate Cortex -- Early Auditory Cortex | Left Dorsal Transitional Visual Area -- Right Lateral Belt Complex | 0.036491573 |
| Intra (Right) | Cerebellum -- Dorsal Stream | Right Cerebellum -- Right Area V3B | 0.037147604 |
| Intra (Left) | Superior Parietal and IPS Cortex -- MT+ Complex and Neighbors | Left Area 7PC -- Left Area PH | 0.037428896 |
| Intra (Right) | Temporal-Parietal-Occipital Junction -- MT+ Complex and Neighbors | Right Superior Temporal Visual Area -- Right Area PH | 0.037769928 |
| Inter | Posterior Cingulate Cortex -- Posterior Cingulate Cortex | Left RetroSplenial Complex -- Right Area 31p ventral | 0.037892478 |
| Inter | Posterior Cingulate Cortex -- Inferior Parietal Cortex | Right Area ventral 23 a+b -- Left Area IntraParietal 2 | 0.037926356 |
| Intra (Right) | Inferior Parietal Cortex -- Lateral Temporal Cortex | Right Area PFm Complex -- Right Area TE1 posterior | 0.039073033 |
| Intra (Right) | Posterior Cingulate Cortex -- MT+ Complex and Neighbors | Right PreCuneus Visual Area -- Right Area PH | 0.039598511 |
| Intra (Right) | Posterior Cingulate Cortex -- Insular and Frontal Opercular Cortex | Right Area 7m -- Right Area Frontal Opercular 5 | 0.03970822 |
| Intra (Right) | Cerebellum -- Inferior Parietal Cortex | Right Cerebellum -- Right Area PGp | 0.039837506 |
| Inter | Caudate -- Lateral Temporal Cortex | Left Caudate -- Right Area PHT | 0.039885317 |
| Intra (Right) | Posterior Cingulate Cortex -- Inferior Parietal Cortex | Right Area ventral 23 a+b -- Right Area PGs | 0.040784972 |
| Intra (Left) | Temporal-Parietal-Occipital Junction -- Early Auditory Cortex | Left PeriSylvian Language Area -- Left Primary Auditory Cortex | 0.040831072 |
| Intra (Left) | Posterior Cingulate Cortex -- Ventral Stream | Left PreCuneus Visual Area -- Left Posterior InferoTemporalComplex | 0.040876804 |
| Intra (Right) | Inferior Frontal Cortex -- Posterior Cingulate Cortex | Right Area 45 -- Right Area 31p ventral | 0.040880509 |
| Intra (Right) | Posterior Cingulate Cortex -- Insular and Frontal Opercular Cortex | Right Area ventral 23 a+b -- Right Area Frontal Opercular 5 | 0.041102019 |
| Inter | Posterior Cingulate Cortex -- Temporal-Parietal-Occipital Junction | Left Parieto-Occipital Sulcus Area 1 -- Right Area TemporoParietoOccipital Junction 3 | 0.041176521 |
| Inter | Dorsolateral Prefrontal Cortex -- Temporal-Parietal-Occipital Junction | Right Area 46 -- Left Area TemporoParietoOccipital Junction 2 | 0.042251214 |
| Inter | Caudate -- Posterior Cingulate Cortex | Right Caudate -- Left Dorsal Transitional Visual Area | 0.042849326 |
| Intra (Right) | Auditory Association Cortex -- Sensorimotor Associated Paracentral Lobular and Mid Cingulate Cortex | Right Area STGa -- Right Area 5L | 0.043497074 |
| Inter | Sensorimotor Associated Paracentral Lobular and Mid Cingulate Cortex -- Dorsal Stream | Right Area 5m -- Left Area V6A | 0.044011955 |
| Inter | Orbital and Polar Frontal Cortex -- Lateral Temporal Cortex | Left Area 47m -- Right Area PHT | 0.044828746 |
| Inter | Dorsal Stream -- Dorsal Stream | Left IntraParietal Sulcus Area 1 -- Right Seventh Visual Area | 0.044844105 |
| Inter | Caudate -- MT+ Complex and Neighbors | Left Caudate -- Right Area PH | 0.045081539 |
| Intra (Right) | Posterior Cingulate Cortex -- Medial Temporal Cortex | Right Area 7m -- Right Perirhinal Ectorhinal Cortex | 0.045170862 |
| Intra (Right) | Early Auditory Cortex -- MT+ Complex and Neighbors | Right Lateral Belt Complex -- Right Area V4t | 0.045911868 |
| Intra (Left) | Superior Parietal and IPS Cortex -- Sensorimotor Associated Paracentral Lobular and Mid Cingulate Cortex | Left Area Lateral IntraParietal dorsal -- Left Area 5L | 0.046147338 |
| Inter | Pallidum -- Ventral Stream | Right Pallidum -- Left Eighth Visual Area | 0.046495914 |
| Inter | Posterior Cingulate Cortex -- Superior Parietal and IPS Cortex | Left PreCuneus Visual Area -- Right Area Lateral IntraParietal dorsal | 0.04666147 |
| Inter | Insular and Frontal Opercular Cortex -- MT+ Complex and Neighbors | Left Pirform Cortex -- Right Area FST | 0.046866113 |
| Intra (Left) | Insular and Frontal Opercular Cortex -- Sensorimotor Associated Paracentral Lobular and Mid Cingulate Cortex | Left Pirform Cortex -- Left Area 5m | 0.047485667 |
| Intra (Right) | Posterior Cingulate Cortex -- MT+ Complex and Neighbors | Right PreCuneus Visual Area -- Right Medial Superior Temporal Area | 0.048085377 |
| Inter | Dorsolateral Prefrontal Cortex -- Lateral Temporal Cortex | Left Area 8C -- Right Area TE2 anterior | 0.048274857 |
| Intra (Right) | Caudate -- Dorsal Stream | Right Caudate -- Right Sixth Visual Area | 0.048277459 |
| Intra (Left) | Temporal-Parietal-Occipital Junction -- Premotor Cortex | Left Superior Temporal Visual Area -- Left Area 55b | 0.048358672 |
| Intra (Right) | Orbital and Polar Frontal Cortex -- Posterior Cingulate Cortex | Right Area 47m -- Right Area dorsal 23 a+b | 0.048671829 |
| Inter | Inferior Parietal Cortex -- Lateral Temporal Cortex | Left Area IntraParietal 2 -- Right Area TE2 anterior | 0.049588374 |
| Inter | Ventral Diencephalon -- MT+ Complex and Neighbors | Left VentralDiencephalon -- Right Area Lateral Occipital 1 | 0.049783673 |
| Intra (Left) | Thalamus -- MT+ Complex and Neighbors | Left Thalamus -- Left Area PH | 0.049824485 |

**Table S1b:** Connections which were found to be significant across the male (MMZ vs MDZ) analysis. All connections were significant with the MZ variance smaller than DZ variance demonstrating genetic influence.

| Hemisphere Type | Network Interaction | Parcel Interaction | p-value |
|---|---|---|---|
| Intra (Right) | Posterior Cingulate Cortex -- Inferior Parietal Cortex | Right Area dorsal 23 a+b -- Right Area IntraParietal 2 | 1.14E-06 |
| Intra (Right) | Superior Parietal and IPS Cortex -- Dorsal Stream | Right Area Lateral IntraParietal ventral -- Right IntraParietal Sulcus Area 1 | 1.60E-05 |
| Intra (Left) | Ventral Stream -- Dorsal Stream | Left VentroMedial Visual Area 2 -- Left Area V3A | 7.60E-05 |
| Inter | Inferior Parietal Cortex -- Medial Temporal Cortex | Left Area IntraParietal 0 -- Right ParaHippocampal Area 3 | 0.000126685 |
| Intra (Left) | MT+ Complex and Neighbors -- MT+ Complex and Neighbors | Left Area PH -- Left Area V3CD | 0.000172811 |
| Intra (Right) | Ventral Stream -- Dorsal Stream | Right VentroMedial Visual Area 1 -- Right Area V3A | 0.000203618 |
| Intra (Left) | Ventral Stream -- Early Visual Cortex | Left Eighth Visual Area -- Left Second Visual Area | 0.000266175 |
| Intra (Right) | Anterior Cingulate and Medial Prefrontal Cortex -- Auditory Association Cortex | Right Area p32 -- Right Area STSv anterior | 0.000268557 |
| Intra (Left) | Ventral Stream -- Dorsal Stream | Left VentroMedial Visual Area 2 -- Left Area V3B | 0.000808759 |
| Intra (Left) | Ventral Stream -- Early Visual Cortex | Left Eighth Visual Area -- Left Third Visual Area | 0.001263647 |
| Inter | Ventral Stream -- Dorsal Stream | Right VentroMedial Visual Area 1 -- Left Area V3A | 0.001354631 |
| Intra (Right) | MT+ Complex and Neighbors -- Ventral Stream | Right Area Lateral Occipital 3 -- Right VentroMedial Visual Area 1 | 0.001621632 |
| Intra (Right) | Anterior Cingulate and Medial Prefrontal Cortex -- Posterior Cingulate Cortex | Right Area 33 prime -- Right Area 23c | 0.001704 |
| Inter | Anterior Cingulate and Medial Prefrontal Cortex -- Anterior Cingulate and Medial Prefrontal Cortex | Left Area 10v -- Right Area 10v | 0.001976739 |
| Intra (Right) | Orbital and Polar Frontal Cortex -- Insular and Frontal Opercular Cortex | Right Posterior OFC Complex -- Right Frontal Opercular Area 3 | 0.002686339 |
| Inter | Inferior Parietal Cortex -- Sensorimotor Associated Paracentral Lobular and Mid Cingulate Cortex | Right Area PF opercular -- Left Area 5m ventral | 0.002779225 |
| Inter | Sensorimotor Associated Paracentral Lobular and Mid Cingulate Cortex -- Somatosensory and Motor Cortex | Left Area 5m -- Right Area 1 | 0.003579041 |
| Intra (Right) | Superior Parietal and IPS Cortex -- Lateral Temporal Cortex | Right Medial IntraParietal Area -- Right Area TE2 posterior | 0.00393788 |
| Intra (Right) | Anterior Cingulate and Medial Prefrontal Cortex -- Superior Parietal and IPS Cortex | Right Area 33 prime -- Right Anterior IntraParietal Area | 0.004957444 |
| Intra (Right) | MT+ Complex and Neighbors -- MT+ Complex and Neighbors | Right Area Lateral Occipital 1 -- Right Area FST | 0.005589921 |
| Inter | Insular and Frontal Opercular Cortex -- Insular and Frontal Opercular Cortex | Left Posterior Insular Area 2 -- Right Area Posterior Insular 1 | 0.005650928 |
| Intra (Left) | Dorsolateral Prefrontal Cortex -- Anterior Cingulate and Medial Prefrontal Cortex | Left Area 9-46d -- Left Area anterior 32 prime | 0.005898452 |
| Intra (Right) | Lateral Temporal Cortex -- Insular and Frontal Opercular Cortex | Right Area PHT -- Right Frontal OPercular Area 2 | 0.0060415 |
| Intra (Left) | Orbital and Polar Frontal Cortex -- Auditory Association Cortex | Left Area anterior 10p -- Left Area STSd anterior | 0.006326714 |
| Intra (Left) | MT+ Complex and Neighbors -- Ventral Stream | Left Area Lateral Occipital 1 -- Left VentroMedial Visual Area 2 | 0.006495927 |
| Intra (Left) | MT+ Complex and Neighbors -- Dorsal Stream | Left Area FST -- Left Area V3A | 0.00737451 |
| Intra (Left) | Cerebellum -- Medial Temporal Cortex | Left Cerebellum -- Left ParaHippocampal Area 2 | 0.007836581 |
| Inter | Cerebellum -- Orbital and Polar Frontal Cortex | Left Cerebellum -- Right Area posterior 10p | 0.008361728 |
| Intra (Left) | Ventral Stream -- Dorsal Stream | Left Eighth Visual Area -- Left Area V3A | 0.009118175 |
| Intra (Left) | Inferior Parietal Cortex -- Inferior Parietal Cortex | Left Area IntraParietal 2 -- Left Area IntraParietal 1 | 0.00935093 |
| Inter | Insular and Frontal Opercular Cortex -- Sensorimotor Associated Paracentral Lobular and Mid Cingulate Cortex | Right Area Posterior Insular 1 -- Left Area 5m | 0.009931853 |
| Inter | Ventral Stream -- Dorsal Stream | Left Ventral Visual Complex -- Right Seventh Visual Area | 0.010451933 |
| Inter | Cerebellum -- Posterior Cingulate Cortex | Left Cerebellum -- Right Area 7m | 0.01078148 |
| Inter | MT+ Complex and Neighbors -- Ventral Stream | Left Area PH -- Right Ventral Visual Complex | 0.010961106 |
| Inter | MT+ Complex and Neighbors -- Ventral Stream | Left Area FST -- Right Fusiform Face Complex | 0.012014434 |
| Inter | Cerebellum -- Inferior Parietal Cortex | Left Cerebellum -- Right Area PGi | 0.01525176 |
| Intra (Right) | Inferior Frontal Cortex -- Premotor Cortex | Right Area IFSp -- Right Premotor Eye Field | 0.015253318 |
| Intra (Right) | Lateral Temporal Cortex -- Ventral Stream | Right Area PHT -- Right VentroMedial Visual Area 2 | 0.015621275 |
| Left BrainStem | Brain Stem -- Accumbens | Bilateral BrainStem -- Left Accumbens | 0.016331142 |
| Inter | MT+ Complex and Neighbors -- MT+ Complex and Neighbors | Left Area PH -- Right Area PH | 0.01890601 |
| Inter | MT+ Complex and Neighbors -- MT+ Complex and Neighbors | Left Area Lateral Occipital 2 -- Right Area PH | 0.019294092 |
| Inter | Ventral Stream -- Early Visual Cortex | Left Eighth Visual Area -- Right Second Visual Area | 0.020711616 |
| Inter | Ventral Stream -- Dorsal Stream | Left VentroMedial Visual Area 2 -- Right Area V3A | 0.022128203 |
| Intra (Left) | MT+ Complex and Neighbors -- MT+ Complex and Neighbors | Left Area FST -- Left Area V3CD | 0.023463234 |
| Intra (Right) | Dorsolateral Prefrontal Cortex -- Inferior Parietal Cortex | Right Area 8C -- Right Area IntraParietal 2 | 0.023514784 |
| Intra (Left) | Sensorimotor Associated Paracentral Lobular and Mid Cingulate Cortex -- Somatosensory and Motor Cortex | Left Area 5m -- Left Primary Sensory Cortex | 0.026442445 |
| Inter | Ventral Stream -- Dorsal Stream | Left VentroMedial Visual Area 2 -- Right Seventh Visual Area | 0.026609617 |
| Inter | Posterior Cingulate Cortex -- Inferior Parietal Cortex | Right Area 31p ventral -- Left Area IntraParietal 1 | 0.026783282 |
| Intra (Left) | Posterior Cingulate Cortex -- Ventral Stream | Left Dorsal Transitional Visual Area -- Left VentroMedial Visual Area 2 | 0.027962836 |
| Inter | MT+ Complex and Neighbors -- Ventral Stream | Left Area Lateral Occipital 1 -- Right VentroMedial Visual Area 3 | 0.028462145 |
| Inter | Inferior Frontal Cortex -- Anterior Cingulate and Medial Prefrontal Cortex | Left Area IFSa -- Right Area 33 prime | 0.029364455 |
| Intra (Left) | Posterior Opercular Cortex -- MT+ Complex and Neighbors | Left Area OP1/SII -- Left Middle Temporal Area | 0.032423712 |
| Inter | Ventral Stream -- Primary Visual Cortex (V1) | Right Ventral Visual Complex -- Left Primary Visual Cortex | 0.032740965 |
| Inter | Inferior Parietal Cortex -- Medial Temporal Cortex | Right Area PGp -- Left ParaHippocampal Area 3 | 0.032840109 |
| Inter | Ventral Stream -- Primary Visual Cortex (V1) | Left VentroMedial Visual Area 2 -- Right Primary Visual Cortex | 0.033173097 |
| Intra (Right) | Ventral Stream -- Dorsal Stream | Right VentroMedial Visual Area 2 -- Right IntraParietal Sulcus Area 1 | 0.033286427 |
| Inter | Orbital and Polar Frontal Cortex -- Temporal-Parietal-Occipital Junction | Right Polar 10p -- Left PeriSylvian Language Area | 0.038093104 |
| Inter | Posterior Cingulate Cortex -- Posterior Cingulate Cortex | Left PreCuneus Visual Area -- Right ProStriate Area | 0.039762221 |
| Intra (Left) | Posterior Cingulate Cortex -- Inferior Parietal Cortex | Left Dorsal Transitional Visual Area -- Left Area IntraParietal 0 | 0.039963865 |
| Intra (Right) | Superior Parietal and IPS Cortex -- Ventral Stream | Right Area Lateral IntraParietal ventral -- Right VentroMedial Visual Area 1 | 0.040415466 |
| Intra (Left) | Superior Parietal and IPS Cortex -- Ventral Stream | Left Lateral Area 7P -- Left VentroMedial Visual Area 2 | 0.040878915 |
| Inter | MT+ Complex and Neighbors -- MT+ Complex and Neighbors | Left Area PH -- Right Area Lateral Occipital 1 | 0.04387106 |
| Intra (Left) | Inferior Parietal Cortex -- Insular and Frontal Opercular Cortex | Left Area PF Complex -- Left Area Frontal Opercular 5 | 0.046144573 |
| Inter | Orbital and Polar Frontal Cortex -- Medial Temporal Cortex | Left Polar 10p -- Right ParaHippocampal Area 1 | 0.049635214 |
| Intra (Left) | Amygdala -- Orbital and Polar Frontal Cortex | Left Amygdala -- Left Area 11l* | 0.027727097 |

**Table S1c:** Connections which were found to be significant across the female (FMZ vs FDZ) analysis. All connections were significant with the MZ variance smaller than DZ variance demonstrating genetic influence.

*The Left Amygdala -- Left Area 11l connection was the only connection which was significant but had variance greater in the MZ group as opposed to the DZ group.

| Parcel | Frequency | Network Name | Network Number |
|---|---|---|---|
| Primary Visual Cortex | 14 | Primary Visual Cortex (V1) | 1 |
| Second Visual Area | 15 | Early Visual Cortex | 2 |
| Fourth Visual Area | 8 | Early Visual Cortex | 2 |
| Third Visual Area | 5 | Early Visual Cortex | 2 |
| IntraParietal Sulcus Area 1 | 25 | Dorsal Stream | 3 |
| Area V3A | 22 | Dorsal Stream | 3 |
| Seventh Visual Area | 16 | Dorsal Stream | 3 |
| Area V3B | 15 | Dorsal Stream | 3 |
| Sixth Visual Area | 13 | Dorsal Stream | 3 |
| Area V6A | 6 | Dorsal Stream | 3 |
| VentroMedial Visual Area 2 | 62 | Ventral Stream | 4 |
| Ventral Visual Complex | 22 | Ventral Stream | 4 |
| VentroMedial Visual Area 1 | 19 | Ventral Stream | 4 |
| Eighth Visual Area | 11 | Ventral Stream | 4 |
| Fusiform Face Complex | 9 | Ventral Stream | 4 |
| VentroMedial Visual Area 3 | 6 | Ventral Stream | 4 |
| Area PH | 35 | MT+ Complex and Neighbors | 5 |
| Area FST | 29 | MT+ Complex and Neighbors | 5 |
| Area Lateral Occipital 1 | 17 | MT+ Complex and Neighbors | 5 |
| Area V3CD | 10 | MT+ Complex and Neighbors | 5 |
| Middle Temporal Area | 8 | MT+ Complex and Neighbors | 5 |
| Medial Superior Temporal Area | 8 | MT+ Complex and Neighbors | 5 |
| Area V4t | 6 | MT+ Complex and Neighbors | 5 |
| Area Lateral Occipital 3 | 5 | MT+ Complex and Neighbors | 5 |
| Area Lateral Occipital 2 | 3 | MT+ Complex and Neighbors | 5 |
| Area 2 | 6 | Somatosensory and Motor Cortex | 6 |
| Primary Sensory Cortex | 3 | Somatosensory and Motor Cortex | 6 |
| Area 1 | 3 | Somatosensory and Motor Cortex | 6 |
| Area 3a | 3 | Somatosensory and Motor Cortex | 6 |
| Primary Motor Cortex | 1 | Somatosensory and Motor Cortex | 6 |
| Area 5m | 40 | Sensorimotor Associated Paracentral Lobular and Mid Cingulate Cortex | 7 |
| Area 5m ventral | 14 | Sensorimotor Associated Paracentral Lobular and Mid Cingulate Cortex | 7 |
| Supplementary and Cingulate Eye Field | 1 | Sensorimotor Associated Paracentral Lobular and Mid Cingulate Cortex | 7 |
| Area 5L | 1 | Sensorimotor Associated Paracentral Lobular and Mid Cingulate Cortex | 7 |
| Dorsal Area 24d | 1 | Sensorimotor Associated Paracentral Lobular and Mid Cingulate Cortex | 7 |
| Premotor Eye Field | 4 | Premotor Cortex | 8 |
| Frontal Eye Fields | 1 | Premotor Cortex | 8 |
| Area 55b | 1 | Premotor Cortex | 8 |
| Area OP1/SII | 5 | Posterior Opercular Cortex | 9 |
| Area PFcm | 5 | Posterior Opercular Cortex | 9 |
| Area OP2-3/VS | 1 | Posterior Opercular Cortex | 9 |
| Frontal Opercular Area 1 | 1 | Posterior Opercular Cortex | 9 |
| Area 43 | 1 | Posterior Opercular Cortex | 9 |
| Lateral Belt Complex | 3 | Early Auditory Cortex | 10 |
| RetroInsular Cortex | 1 | Early Auditory Cortex | 10 |
| ParaBelt Complex | 1 | Early Auditory Cortex | 10 |
| Area STGa | 6 | Auditory Association Cortex | 11 |
| Area STSv anterior | 4 | Auditory Association Cortex | 11 |
| Area STSv posterior | 3 | Auditory Association Cortex | 11 |
| Area STSd posterior | 3 | Auditory Association Cortex | 11 |
| Auditory 4 Complex | 2 | Auditory Association Cortex | 11 |
| Area Posterior Insular 1 | 9 | Insular and Frontal Opercular Cortex | 12 |
| Insular Granular Complex | 8 | Insular and Frontal Opercular Cortex | 12 |
| Area Frontal Opercular 5 | 4 | Insular and Frontal Opercular Cortex | 12 |
| Middle Insular Area | 3 | Insular and Frontal Opercular Cortex | 12 |
| Posterior Insular Area 2 | 1 | Insular and Frontal Opercular Cortex | 12 |
| Anterior Agranular Insula Complex | 1 | Insular and Frontal Opercular Cortex | 12 |
| Frontal OPercular Area 2 | 1 | Insular and Frontal Opercular Cortex | 12 |
| ParaHippocampal Area 3 | 18 | Medial Temporal Cortex | 13 |
| ParaHippocampal Area 2 | 12 | Medial Temporal Cortex | 13 |
| PreSubiculum | 8 | Medial Temporal Cortex | 13 |
| ParaHippocampal Area 1 | 6 | Medial Temporal Cortex | 13 |
| Entorhinal Cortex | 4 | Medial Temporal Cortex | 13 |
| Perirhinal Ectorhinal Cortex | 1 | Medial Temporal Cortex | 13 |
| Hippocampus | 1 | Medial Temporal Cortex | 13 |
| Area TF | 12 | Lateral Temporal Cortex | 14 |
| Area TE2 posterior | 11 | Lateral Temporal Cortex | 14 |
| Area TE1 posterior | 8 | Lateral Temporal Cortex | 14 |

| | | | |
|---|---|---|---|
| Area PHT | 5 | Lateral Temporal Cortex | 14 |
| Area TE2 anterior | 4 | Lateral Temporal Cortex | 14 |
| Area TemporoParietoOccipital Junction 2 | 8 | Temporal-Parietal-Occipital Junction | 15 |
| Area TemporoParietoOccipital Junction 1 | 8 | Temporal-Parietal-Occipital Junction | 15 |
| Area TemporoParietoOccipital Junction 3 | 8 | Temporal-Parietal-Occipital Junction | 15 |
| Superior Temporal Visual Area | 4 | Temporal-Parietal-Occipital Junction | 15 |
| Area Lateral IntraParietal dorsal | 20 | Superior Parietal and IPS Cortex | 16 |
| Area Lateral IntraParietal ventral | 14 | Superior Parietal and IPS Cortex | 16 |
| Medial IntraParietal Area | 6 | Superior Parietal and IPS Cortex | 16 |
| Anterior IntraParietal Area | 4 | Superior Parietal and IPS Cortex | 16 |
| Lateral Area 7P | 4 | Superior Parietal and IPS Cortex | 16 |
| Medial Area 7P | 3 | Superior Parietal and IPS Cortex | 16 |
| Area 7PC | 3 | Superior Parietal and IPS Cortex | 16 |
| Medial Area 7A | 3 | Superior Parietal and IPS Cortex | 16 |
| Ventral IntraParietal Complex | 2 | Superior Parietal and IPS Cortex | 16 |
| Lateral Area 7A | 1 | Superior Parietal and IPS Cortex | 16 |
| Area IntraParietal 0 | 18 | Inferior Parietal Cortex | 17 |
| Area IntraParietal 1 | 17 | Inferior Parietal Cortex | 17 |
| Area IntraParietal 2 | 6 | Inferior Parietal Cortex | 17 |
| Area PF opercular | 5 | Inferior Parietal Cortex | 17 |
| Area PGs | 5 | Inferior Parietal Cortex | 17 |
| Area PGi | 4 | Inferior Parietal Cortex | 17 |
| Area PFt | 4 | Inferior Parietal Cortex | 17 |
| Area PFm Complex | 4 | Inferior Parietal Cortex | 17 |
| Area PGp | 3 | Inferior Parietal Cortex | 17 |
| Area PF Complex | 1 | Inferior Parietal Cortex | 17 |
| PreCuneus Visual Area | 54 | Posterior Cingulate Cortex | 18 |
| Area ventral 23 a+b | 31 | Posterior Cingulate Cortex | 18 |
| Dorsal Transitional Visual Area | 16 | Posterior Cingulate Cortex | 18 |
| Area 31pd | 13 | Posterior Cingulate Cortex | 18 |
| Area dorsal 23 a+b | 9 | Posterior Cingulate Cortex | 18 |
| Parieto-Occipital Sulcus Area 1 | 9 | Posterior Cingulate Cortex | 18 |
| ProStriate Area | 7 | Posterior Cingulate Cortex | 18 |
| Area 23c | 5 | Posterior Cingulate Cortex | 18 |
| Area 31p ventral | 5 | Posterior Cingulate Cortex | 18 |
| Parieto-Occipital Sulcus Area 2 | 4 | Posterior Cingulate Cortex | 18 |
| Area 7m | 4 | Posterior Cingulate Cortex | 18 |
| Area 31a | 2 | Posterior Cingulate Cortex | 18 |
| Area 23d | 2 | Posterior Cingulate Cortex | 18 |
| RetroSplenial Complex | 1 | Posterior Cingulate Cortex | 18 |
| Area 10v | 3 | Anterior Cingulate and Medial Prefrontal Cortex | 19 |
| Area 9 Middle | 2 | Anterior Cingulate and Medial Prefrontal Cortex | 19 |
| Area p32 | 2 | Anterior Cingulate and Medial Prefrontal Cortex | 19 |
| Area s32 | 2 | Anterior Cingulate and Medial Prefrontal Cortex | 19 |
| Area 25 | 1 | Anterior Cingulate and Medial Prefrontal Cortex | 19 |
| Area dorsal 32 | 1 | Anterior Cingulate and Medial Prefrontal Cortex | 19 |
| Area 33 prime | 1 | Anterior Cingulate and Medial Prefrontal Cortex | 19 |
| Area 8BM | 1 | Anterior Cingulate and Medial Prefrontal Cortex | 19 |
| Area posterior 24 | 1 | Anterior Cingulate and Medial Prefrontal Cortex | 19 |
| Anterior 24 prime | 1 | Anterior Cingulate and Medial Prefrontal Cortex | 19 |
| Area 10r | 1 | Anterior Cingulate and Medial Prefrontal Cortex | 19 |
| Area 47m | 6 | Orbital and Polar Frontal Cortex | 20 |
| Area posterior 10p | 3 | Orbital and Polar Frontal Cortex | 20 |
| Area 13l | 2 | Orbital and Polar Frontal Cortex | 20 |
| Area anterior 47r | 2 | Orbital and Polar Frontal Cortex | 20 |
| Area 47s | 2 | Orbital and Polar Frontal Cortex | 20 |
| Polar 10p | 2 | Orbital and Polar Frontal Cortex | 20 |
| Posterior OFC Complex | 1 | Orbital and Polar Frontal Cortex | 20 |
| Area 10d | 1 | Orbital and Polar Frontal Cortex | 20 |
| Area 47l (47 lateral) | 9 | Inferior Frontal Cortex | 21 |
| Area 45 | 7 | Inferior Frontal Cortex | 21 |
| Area IFJp | 6 | Inferior Frontal Cortex | 21 |
| Area 44 | 4 | Inferior Frontal Cortex | 21 |
| Area IFSp | 3 | Inferior Frontal Cortex | 21 |
| Area IFSa | 3 | Inferior Frontal Cortex | 21 |
| Area IFJa | 2 | Inferior Frontal Cortex | 21 |
| Area anterior 9-46v | 2 | Dorsolateral Prefrontal Cortex | 22 |
| Area 9 anterior | 2 | Dorsolateral Prefrontal Cortex | 22 |
| Area 46 | 2 | Dorsolateral Prefrontal Cortex | 22 |

| | | | |
|---|---|---|---|
| Area 8B Lateral | 2 | Dorsolateral Prefrontal Cortex | 22 |
| Area 8Av | 2 | Dorsolateral Prefrontal Cortex | 22 |
| Area 9 Posterior | 1 | Dorsolateral Prefrontal Cortex | 22 |
| Area 9-46d | 1 | Dorsolateral Prefrontal Cortex | 22 |
| Area 8Ad | 1 | Dorsolateral Prefrontal Cortex | 22 |
| Area posterior 9-46v | 1 | Dorsolateral Prefrontal Cortex | 22 |
| Area 8C | 1 | Dorsolateral Prefrontal Cortex | 22 |
| Subcortical Hippocampus | 23 | Hippocampus | 24 |
| Accumbens | 1 | Accumbens | 25 |
| Caudate | 20 | Caudate | 26 |
| Thalamus | 5 | Thalamus | 29 |
| VentralDiencephalon | 4 | Ventral Diencephalon | 31 |
| Cerebellum | 9 | Cerebellum | 32 |

**Table S2a:** Parcel frequencies demonstrating genetic influence from significant connections broken into parcels (MZ vs DZ analysis).

| Parcel | Frequency | Network Name | Network Number |
|---|---|---|---|
| Primary Visual Cortex | 1 | Primary Visual Cortex (V1) | 1 |
| Fourth Visual Area | 6 | Early Visual Cortex | 2 |
| Second Visual Area | 4 | Early Visual Cortex | 2 |
| Third Visual Area | 3 | Early Visual Cortex | 2 |
| IntraParietal Sulcus Area 1 | 22 | Dorsal Stream | 3 |
| Area V3B | 6 | Dorsal Stream | 3 |
| Area V6A | 5 | Dorsal Stream | 3 |
| Seventh Visual Area | 4 | Dorsal Stream | 3 |
| Sixth Visual Area | 3 | Dorsal Stream | 3 |
| Posterior InferoTemporalComplex | 6 | Ventral Stream | 4 |
| Eighth Visual Area | 5 | Ventral Stream | 4 |
| Fusiform Face Complex | 4 | Ventral Stream | 4 |
| VentroMedial Visual Area 3 | 3 | Ventral Stream | 4 |
| VentroMedial Visual Area 2 | 3 | Ventral Stream | 4 |
| Ventral Visual Complex | 3 | Ventral Stream | 4 |
| Area PH | 22 | MT+ Complex and Neighbors | 5 |
| Area Lateral Occipital 1 | 7 | MT+ Complex and Neighbors | 5 |
| Area V3CD | 6 | MT+ Complex and Neighbors | 5 |
| Area FST | 5 | MT+ Complex and Neighbors | 5 |
| Area Lateral Occipital 2 | 3 | MT+ Complex and Neighbors | 5 |
| Area V4t | 3 | MT+ Complex and Neighbors | 5 |
| Area Lateral Occipital 3 | 2 | MT+ Complex and Neighbors | 5 |
| Middle Temporal Area | 1 | MT+ Complex and Neighbors | 5 |
| Medial Superior Temporal Area | 1 | MT+ Complex and Neighbors | 5 |
| Area 5m | 9 | Sensorimotor Associated Paracentral Lobular and Mid Cingulate Cortex | 7 |
| Area 5m ventral | 7 | Sensorimotor Associated Paracentral Lobular and Mid Cingulate Cortex | 7 |
| Area 5L | 5 | Sensorimotor Associated Paracentral Lobular and Mid Cingulate Cortex | 7 |
| Ventral Area 24d | 2 | Sensorimotor Associated Paracentral Lobular and Mid Cingulate Cortex | 7 |
| Dorsal Area 24d | 2 | Sensorimotor Associated Paracentral Lobular and Mid Cingulate Cortex | 7 |
| Area 55b | 6 | Premotor Cortex | 8 |
| Area 6 anterior | 3 | Premotor Cortex | 8 |
| Frontal Eye Fields | 2 | Premotor Cortex | 8 |
| Rostral Area 6 | 1 | Premotor Cortex | 8 |
| Premotor Eye Field | 1 | Premotor Cortex | 8 |
| Area PFcm | 3 | Posterior Opercular Cortex | 9 |
| Area OP2-3/VS | 1 | Posterior Opercular Cortex | 9 |
| Area OP4/PV | 1 | Posterior Opercular Cortex | 9 |
| Area OP1/SII | 1 | Posterior Opercular Cortex | 9 |
| Lateral Belt Complex | 12 | Early Auditory Cortex | 10 |
| RetroInsular Cortex | 4 | Early Auditory Cortex | 10 |
| Primary Auditory Cortex | 2 | Early Auditory Cortex | 10 |
| ParaBelt Complex | 1 | Early Auditory Cortex | 10 |
| Area STSd posterior | 9 | Auditory Association Cortex | 11 |
| Area STGa | 5 | Auditory Association Cortex | 11 |
| Area STSv anterior | 3 | Auditory Association Cortex | 11 |
| Area STSv posterior | 3 | Auditory Association Cortex | 11 |
| Auditory 4 Complex | 3 | Auditory Association Cortex | 11 |
| Area STSd anterior | 1 | Auditory Association Cortex | 11 |
| Insular Granular Complex | 4 | Insular and Frontal Opercular Cortex | 12 |
| Area Frontal Opercular 5 | 3 | Insular and Frontal Opercular Cortex | 12 |
| Pirform Cortex | 3 | Insular and Frontal Opercular Cortex | 12 |
| Anterior Agranular Insula Complex | 2 | Insular and Frontal Opercular Cortex | 12 |
| Area 52 | 1 | Insular and Frontal Opercular Cortex | 12 |
| Frontal Opercular Area 3 | 1 | Insular and Frontal Opercular Cortex | 12 |
| Area Posterior Insular 1 | 1 | Insular and Frontal Opercular Cortex | 12 |
| ParaHippocampal Area 3 | 5 | Medial Temporal Cortex | 13 |
| PreSubiculum | 4 | Medial Temporal Cortex | 13 |
| ParaHippocampal Area 1 | 2 | Medial Temporal Cortex | 13 |
| Perirhinal Ectorhinal Cortex | 2 | Medial Temporal Cortex | 13 |
| ParaHippocampal Area 2 | 1 | Medial Temporal Cortex | 13 |
| Entorhinal Cortex | 1 | Medial Temporal Cortex | 13 |
| Area PHT | 8 | Lateral Temporal Cortex | 14 |
| Area TG Ventral | 6 | Lateral Temporal Cortex | 14 |
| Area TE2 anterior | 6 | Lateral Temporal Cortex | 14 |
| Area TF | 3 | Lateral Temporal Cortex | 14 |
| Area TE2 posterior | 2 | Lateral Temporal Cortex | 14 |
| Area TE1 posterior | 2 | Lateral Temporal Cortex | 14 |
| Area TemporoParietoOccipital Junction 3 | 12 | Temporal-Parietal-Occipital Junction | 15 |
| Area TemporoParietoOccipital Junction 2 | 8 | Temporal-Parietal-Occipital Junction | 15 |
| PeriSylvian Language Area | 5 | Temporal-Parietal-Occipital Junction | 15 |

| | | | |
|---|---|---|---|
| Area TemporoParietoOccipital Junction 1 | 4 | Temporal-Parietal-Occipital Junction | 15 |
| Superior Temporal Visual Area | 4 | Temporal-Parietal-Occipital Junction | 15 |
| Area Lateral IntraParietal dorsal | 17 | Superior Parietal and IPS Cortex | 16 |
| Area Lateral IntraParietal ventral | 5 | Superior Parietal and IPS Cortex | 16 |
| Anterior IntraParietal Area | 3 | Superior Parietal and IPS Cortex | 16 |
| Medial Area 7A | 3 | Superior Parietal and IPS Cortex | 16 |
| Lateral Area 7A | 2 | Superior Parietal and IPS Cortex | 16 |
| Medial IntraParietal Area | 2 | Superior Parietal and IPS Cortex | 16 |
| Area 7PC | 2 | Superior Parietal and IPS Cortex | 16 |
| Medial Area 7P | 1 | Superior Parietal and IPS Cortex | 16 |
| Lateral Area 7P | 1 | Superior Parietal and IPS Cortex | 16 |
| Area IntraParietal 0 | 10 | Inferior Parietal Cortex | 17 |
| Area IntraParietal 1 | 9 | Inferior Parietal Cortex | 17 |
| Area PFm Complex | 8 | Inferior Parietal Cortex | 17 |
| Area IntraParietal 2 | 6 | Inferior Parietal Cortex | 17 |
| Area PGp | 3 | Inferior Parietal Cortex | 17 |
| Area PF Complex | 3 | Inferior Parietal Cortex | 17 |
| Area PGs | 3 | Inferior Parietal Cortex | 17 |
| Area PGi | 2 | Inferior Parietal Cortex | 17 |
| Area PFt | 1 | Inferior Parietal Cortex | 17 |
| PreCuneus Visual Area | 26 | Posterior Cingulate Cortex | 18 |
| Area ventral 23 a+b | 18 | Posterior Cingulate Cortex | 18 |
| Area 7m | 12 | Posterior Cingulate Cortex | 18 |
| Area 23c | 8 | Posterior Cingulate Cortex | 18 |
| Parieto-Occipital Sulcus Area 1 | 7 | Posterior Cingulate Cortex | 18 |
| Area dorsal 23 a+b | 7 | Posterior Cingulate Cortex | 18 |
| Parieto-Occipital Sulcus Area 2 | 5 | Posterior Cingulate Cortex | 18 |
| Area 31pd | 4 | Posterior Cingulate Cortex | 18 |
| Area 23d | 4 | Posterior Cingulate Cortex | 18 |
| Area 31p ventral | 4 | Posterior Cingulate Cortex | 18 |
| RetroSplenial Complex | 4 | Posterior Cingulate Cortex | 18 |
| ProStriate Area | 3 | Posterior Cingulate Cortex | 18 |
| Dorsal Transitional Visual Area | 3 | Posterior Cingulate Cortex | 18 |
| Area 31a | 1 | Posterior Cingulate Cortex | 18 |
| Anterior 24 prime | 2 | Anterior Cingulate and Medial Prefrontal Cortex | 19 |
| Area a24 | 1 | Anterior Cingulate and Medial Prefrontal Cortex | 19 |
| Area 10r | 1 | Anterior Cingulate and Medial Prefrontal Cortex | 19 |
| Area 25 | 1 | Anterior Cingulate and Medial Prefrontal Cortex | 19 |
| Area Posterior 24 prime | 1 | Anterior Cingulate and Medial Prefrontal Cortex | 19 |
| Area dorsal 32 | 1 | Anterior Cingulate and Medial Prefrontal Cortex | 19 |
| Area 47m | 8 | Orbital and Polar Frontal Cortex | 20 |
| Area 13l | 5 | Orbital and Polar Frontal Cortex | 20 |
| Area 47s | 2 | Orbital and Polar Frontal Cortex | 20 |
| Area 10d | 2 | Orbital and Polar Frontal Cortex | 20 |
| Area 11l | 1 | Orbital and Polar Frontal Cortex | 20 |
| Area 45 | 4 | Inferior Frontal Cortex | 21 |
| Area IFSp | 1 | Inferior Frontal Cortex | 21 |
| Area 47l (47 lateral) | 1 | Inferior Frontal Cortex | 21 |
| Area 44 | 1 | Inferior Frontal Cortex | 21 |
| Area posterior 47r | 1 | Inferior Frontal Cortex | 21 |
| Area IFSa | 1 | Inferior Frontal Cortex | 21 |
| Area 46 | 3 | Dorsolateral Prefrontal Cortex | 22 |
| Area 9 Posterior | 2 | Dorsolateral Prefrontal Cortex | 22 |
| Superior 6-8 Translational Area | 2 | Dorsolateral Prefrontal Cortex | 22 |
| Area 9-46d | 1 | Dorsolateral Prefrontal Cortex | 22 |
| Inferior 6-8 Transitional Area | 1 | Dorsolateral Prefrontal Cortex | 22 |
| Area 8Ad | 1 | Dorsolateral Prefrontal Cortex | 22 |
| Area posterior 9-46v | 1 | Dorsolateral Prefrontal Cortex | 22 |
| Area 8Av | 1 | Dorsolateral Prefrontal Cortex | 22 |
| Area 8C | 1 | Dorsolateral Prefrontal Cortex | 22 |
| Amygdala | 1 | Amygdala | 23 |
| Subcortical Hippocampus | 5 | Hippocampus | 24 |
| Accumbens | 1 | Accumbens | 25 |
| Caudate | 15 | Caudate | 26 |
| Pallidum | 6 | Pallidum | 27 |
| Thalamus | 31 | Thalamus | 29 |
| VentralDiencephalon | 12 | Ventral Diencephalon | 31 |
| Cerebellum | 10 | Cerebellum | 32 |

**Table S2b:** Parcel frequencies demonstrating genetic influence from significant connections broken into parcels for males (MMZ vs MDZ analysis).

| Parcel | Frequency | Network Name | Network Number |
|---|---|---|---|
| Primary Visual Cortex | 2 | Primary Visual Cortex (V1) | 1 |
| Second Visual Area | 2 | Early Visual Cortex | 2 |
| Third Visual Area | 1 | Early Visual Cortex | 2 |
| Area V3A | 6 | Dorsal Stream | 3 |
| IntraParietal Sulcus Area 1 | 2 | Dorsal Stream | 3 |
| Seventh Visual Area | 2 | Dorsal Stream | 3 |
| Area V3B | 1 | Dorsal Stream | 3 |
| VentroMedial Visual Area 2 | 10 | Ventral Stream | 4 |
| VentroMedial Visual Area 1 | 4 | Ventral Stream | 4 |
| Eighth Visual Area | 4 | Ventral Stream | 4 |
| Ventral Visual Complex | 3 | Ventral Stream | 4 |
| Fusiform Face Complex | 1 | Ventral Stream | 4 |
| VentroMedial Visual Area 3 | 1 | Ventral Stream | 4 |
| Area PH | 5 | MT+ Complex and Neighbors | 5 |
| Area Lateral Occipital 1 | 4 | MT+ Complex and Neighbors | 5 |
| Area FST | 4 | MT+ Complex and Neighbors | 5 |
| Area V3CD | 3 | MT+ Complex and Neighbors | 5 |
| Area Lateral Occipital 3 | 1 | MT+ Complex and Neighbors | 5 |
| Area Lateral Occipital 2 | 1 | MT+ Complex and Neighbors | 5 |
| Middle Temporal Area | 1 | MT+ Complex and Neighbors | 5 |
| Area 1 | 1 | Somatosensory and Motor Cortex | 6 |
| Primary Sensory Cortex | 1 | Somatosensory and Motor Cortex | 6 |
| Area 5m | 3 | Sensorimotor Associated Paracentral Lobular and Mid Cingulate Cortex | 7 |
| Area 5m ventral | 1 | Sensorimotor Associated Paracentral Lobular and Mid Cingulate Cortex | 7 |
| Premotor Eye Field | 1 | Premotor Cortex | 8 |
| Area OP1/SII | 1 | Posterior Opercular Cortex | 9 |
| Area STSv anterior | 1 | Auditory Association Cortex | 11 |
| Area STSd anterior | 1 | Auditory Association Cortex | 11 |
| Area Posterior Insular 1 | 2 | Insular and Frontal Opercular Cortex | 12 |
| Frontal Opercular Area 3 | 1 | Insular and Frontal Opercular Cortex | 12 |
| Posterior Insular Area 2 | 1 | Insular and Frontal Opercular Cortex | 12 |
| Frontal OPercular Area 2 | 1 | Insular and Frontal Opercular Cortex | 12 |
| Area Frontal Opercular 5 | 1 | Insular and Frontal Opercular Cortex | 12 |
| ParaHippocampal Area 3 | 2 | Medial Temporal Cortex | 13 |
| ParaHippocampal Area 2 | 1 | Medial Temporal Cortex | 13 |
| ParaHippocampal Area 1 | 1 | Medial Temporal Cortex | 13 |
| Area PHT | 2 | Lateral Temporal Cortex | 14 |
| Area TE2 posterior | 1 | Lateral Temporal Cortex | 14 |
| PeriSylvian Language Area | 1 | Temporal-Parietal-Occipital Junction | 15 |
| Area Lateral IntraParietal ventral | 2 | Superior Parietal and IPS Cortex | 16 |
| Medial IntraParietal Area | 1 | Superior Parietal and IPS Cortex | 16 |
| Anterior IntraParietal Area | 1 | Superior Parietal and IPS Cortex | 16 |
| Lateral Area 7P | 1 | Superior Parietal and IPS Cortex | 16 |
| Area IntraParietal 2 | 3 | Inferior Parietal Cortex | 17 |
| Area IntraParietal 0 | 2 | Inferior Parietal Cortex | 17 |
| Area IntraParietal 1 | 2 | Inferior Parietal Cortex | 17 |
| Area PF opercular | 1 | Inferior Parietal Cortex | 17 |
| Area PGi | 1 | Inferior Parietal Cortex | 17 |
| Area PGp | 1 | Inferior Parietal Cortex | 17 |
| Area PF Complex | 1 | Inferior Parietal Cortex | 17 |
| Dorsal Transitional Visual Area | 2 | Posterior Cingulate Cortex | 18 |
| Area dorsal 23 a+b | 1 | Posterior Cingulate Cortex | 18 |
| Area 23c | 1 | Posterior Cingulate Cortex | 18 |
| Area 7m | 1 | Posterior Cingulate Cortex | 18 |
| Area 31p ventral | 1 | Posterior Cingulate Cortex | 18 |
| PreCuneus Visual Area | 1 | Posterior Cingulate Cortex | 18 |
| ProStriate Area | 1 | Posterior Cingulate Cortex | 18 |
| Area 33 prime | 3 | Anterior Cingulate and Medial Prefrontal Cortex | 19 |
| Area 10v | 2 | Anterior Cingulate and Medial Prefrontal Cortex | 19 |
| Area p32 | 1 | Anterior Cingulate and Medial Prefrontal Cortex | 19 |
| Area anterior 32 prime | 1 | Anterior Cingulate and Medial Prefrontal Cortex | 19 |
| Polar 10p | 2 | Orbital and Polar Frontal Cortex | 20 |
| Posterior OFC Complex | 1 | Orbital and Polar Frontal Cortex | 20 |
| Area anterior 10p | 1 | Orbital and Polar Frontal Cortex | 20 |
| Area posterior 10p | 1 | Orbital and Polar Frontal Cortex | 20 |
| Area IFSp | 1 | Inferior Frontal Cortex | 21 |
| Area IFSa | 1 | Inferior Frontal Cortex | 21 |
| Area 9-46d | 1 | Dorsolateral Prefrontal Cortex | 22 |
| Area 8C | 1 | Dorsolateral Prefrontal Cortex | 22 |
| Accumbens | 1 | Accumbens | 25 |
| BrainStem | 1 | Brain Stem | 30 |
| Cerebellum | 4 | Cerebellum | 32 |

**Table S2c:** Parcel frequencies demonstrating genetic influence from significant connections broken into parcels for females (FMZ vs FDZ analysis).
